# Virus-free continuous directed evolution in human cells using somatic hypermutation

**DOI:** 10.1101/2024.12.24.629435

**Authors:** Stanley Bram, Diego J. Orea, Graeme Lindsey, Soyu Zi, Jordan Quenneville, Hui Xu, Sarah Leach, Jenna J. Guthmiller, Angad P. Mehta

**Author notes:** Contributed equally to this work (Stanley Bram, Diego J. Orea). Contributed equally to this work (Graeme Lindsey, Soyu Zi).

## Abstract

The B cells of the human immune system have evolved somatic hypermutation (SHM) mechanisms that introduce mutations at the immunoglobulin genomic loci at a significantly higher frequency than the rest of the genome thereby allowing them to evolve antibody sequences without compromising fitness due to genome-wide mutations. Inspired by these observations, here we developed a continuous directed evolution platform in human B cell lines (CODE-HB) that repurposes the SHM mechanisms to a stable, non-immunoglobulin genomic locus of the human B cell lines to continuously evolve cytosolic and surface displayed proteins with a broad mutational spectrum comprising of substitutions, deletions and insertions. We developed a human B cell surface display platform and used CODE-HB to evolve neutralizing antibodies targeting avian influenza and escape variants of influenza. Given the modularity and simplicity of CODE-HB, we anticipate that this platform can be used for rapidly evolving biotechnologically relevant biomolecules directly in human cells.

## Introduction

Random mutations followed by selection is one of the key mechanisms that results in evolution of new functions in nature. Typically, mutational rates are low so as not to compromise the fitness of the organism as a result of genome-wide mutations. However, this protection against error also results in a relatively slow diversification of biomolecules and the subsequent evolution of new functions. For example, in a typical bacterial or human cell the mutational frequency is around 10^-10^ to 10^-9^ substitution per base.^1^ Therefore, for a mutation to occur within a single gene of around 1000 bases, a cell would have to undergo more than a million rounds of cell division. The rate of evolution is further constrained by the number of mutations a genome can tolerate before the organism is not viable. This limit is especially important in the context of directed evolution experiments that rely on biomolecular diversity to evolve biomolecules (proteins, RNAs) with desired functions. Directed evolution has traditionally relied on *in vitro* techniques, such as utilizing randomized oligonucleotide sequences or error-prone polymerase chain reactions (PCR), to diversify genes encoding target biomolecules rather than relying on low mutation frequencies *in vivo*. After *in vitro* diversification, subsequent steps in directed evolution involve screening or selection methods to enrich and identify variants that exhibit a desired phenotype, followed by detailed characterization to establish the link between genotype and phenotype. Given the process’s iterative requirements, each round of a directed evolution campaign requires generating biomolecular diversity in vitro before proceeding with screening and selection, rendering it a labor-intensive and time-consuming approach to profile the evolutionary landscape and identify biomolecules with desired properties.^2–7^

To overcome some of these limitations, various continuous directed evolution platforms have been developed^8^. Orthogonal replicons with high mutagenic rates have been established in model bacteria like *E. coli*^9^ or model eukaryotes like yeasts^10,11^. Genes encoding proteins of interests are encoded in orthogonal replicons, and are subjected to continuous mutagenesis by expressing an error prone DNA polymerase that specifically replicates the orthogonal replication system and not the genomic DNA. This approach results in the continuous generation of a library of biomolecular variants within the cells without compromising their viability due to genomic mutations. The cells expressing a library of variants are screened under selection conditions to identify cells expressing variants resulting in a desired phenotype (e.g., antibiotic resistance, protein/protein binding). This step is typically followed by sequencing experiments to identify the mutant gene encoding the selected protein variant. Similar to orthogonal replicons, transcription coupled DNA modification enzymes (e.g., T7 RNA polymerases that have been fused to deaminases) have been engineered to create DNA mismatches at C:G base pairs, which upon repair result in the generation of biomolecular diversity.^12–14^ In addition to these cell-based approaches, virus-based continuous evolution approaches have been developed.^15–18^ In these platforms either the viral genome or viral helper plasmids encode genes corresponding to the protein of interest. Biomolecular diversity is generated by either engineering mutagenic polymerases or by expressing mutagenic factors. These approaches typically link evolutionary outcomes (binding, enzyme activity) to viral propagation. Each of these approaches requires extensive engineering efforts and several of the continuous directed evolution efforts in mammalian cells are typically virus mediated approaches.^16–18^

The studies described here were inspired by the mammalian adaptive immune system, which has evolved mechanisms to overcome the evolutionary bottleneck of low error frequencies and mutational tolerance to rapidly evolve proteins in response to antigen exposure. B cells use DNA rearrangement, recombination and somatic hypermutation (SHM) mechanisms to rapidly evolve antibodies.^19–22^ The SHM machinery allows B cells to introduce mutations specifically at immunoglobulin genomic loci at a significantly higher frequency than the rest of the genome.^20^ This selective diversification allows B cells to rapidly evolve new antibody sequences without compromising genomic fitness due to mutational burden, thus providing an attractive feature for biomolecular evolution. Importantly, SHM mediated mutagenesis also results in relatively broad mutational profile comprising of substitutions, insertions and deletions.^23,24^ Indeed, SHM mechanisms have been leveraged by using animal immunization followed by B cell maturation process to identify new antibodies targeting defined epitopes^25^ as well as for evolving catalytic antibodies^26^. Moreover, studies have attempted evolution of optimized B cell receptor sequences and IgMs by incorporating the corresponding genetic elements at the immunoglobulin loci of B cells.^27–31^ Similar to these studies, chicken lymphoma B cell line has also been explored to transiently evolve proteins from immunoglobulin loci.^32–35^ B cells have also been engineered to express custom antibodies targeting defined epitopes as potential therapeutics.^28,29,36^ Of particular interest to this work, a pioneering study by Tsien and coworkers demonstrated that low frequency-virus based genomic integration methods resulted in transient expression and evolution of fluorescent proteins when the corresponding gene was integrated at the immunoglobulin locus.^37^ However, low integration frequency along with genetically instability and sustained expression from such loci are undesirable traits for continuous directed evolution.^24,38^ In spite of all the these efforts, B cell-based laboratory evolution experiments have not found widespread utility. These observations led us to investigate if could overcome the bottlenecks in unlocking the remarkable potential of human B cells for continuous directed evolution of proteins directly in human cells.

Here, we describe a novel virus-free <u>Co</u>ntinuous <u>D</u>irected <u>E</u>volution platform in <u>H</u>uman <u>B</u> cell lines (CODE-HB) that recruits and repurposes the inherent B cell SHM mechanisms to rapidly, and orthogonally, evolve proteins of interest. We first developed precise genome editing approaches to incorporate defined genetic elements at a stable, non-immunoglobulin locus in human B cell line genomes. This allowed us to achieve stable high levels of protein expression in more than 90% of the cell population. To continuously evolve reporter proteins expressed from this locus, we identified and added a series of defined DNA elements from an immunoglobulin locus upstream of the gene encoding a reporter protein, facilitating the recruitment of the B cell SHM machinery to this non-immunoglobulin stable locus. We demonstrate that these engineered human B cell lines can rapidly evolve reporter proteins (e.g., enhanced Green Fluorescent Protein, eGFP). To adapt this platform to rapidly evolve membrane displayed proteins, we developed a human B cell surface display platform and combine it with CODE-HB. Using this approach, we display Fragment antigen binding domains (F_ab_) of antibodies on surface of human B cells and use CODE-HB to evolve these F_ab_ sequences for targeting avian subtypes of influenza hemagglutinin. This resulted in the identification of H5 targeting antibodies. Such antibodies are important in the context of recent zoonotic spread of H5N1 strains of avian influenza.^39^ We also show that our platform can be used to evolve antibodies to escape variants of hemagglutinin. Further, we demonstrate that our evolved antibodies are more potent that the parent antibodies in viral neutralization assays. We comprehensively characterize the mutational profile and breadth of this approach by using single-molecule sequencing experiments and we observe a broad mutational profile comprising of substitutions, deletions and insertions which is unique to our approach as compared to most other most other continuous directed evolution approaches that are commonly used. Given the modularity and simplicity of CODE-HB to rapidly evolve both cytoplasmic proteins as well as surface displayed proteins, we anticipate that this platform can be used for developing biologics as well as for proteins and enzymes directly in human cell lines. Furthermore, this engineered stable system can be used to provide new insights into the SHM mechanisms.

## Results

### Precise and stable genome editing of human B cell lines for reporter protein expression

Our first goal was to develop an approach that would allow incorporation of defined genetic elements at a defined stable locus in B cell lines. Most commonly, lentivirus-based approaches have been used to randomly incorporate genetic elements in the genomes of B cell lines.^37^ Recently, CRISPR/Cas9 plasmids along with adeno-associated virus (AAV) containing recombination cassettes have been used in B cell lines to replace endogenously encoded antibodies with heterologous ones targeting specific antigens^36^. For our studies, we investigated if we could develop a relatively simple, virus-free, plasmid DNA-based approach to edit genomes of B cell lines with a goal to recombinantly express reporter proteins (Figure 1A-C) from a defined and stable genomic locus. We envisioned using a two-plasmid system composed of (i) a CRISPR/Cas9 plasmid DNA containing genetic elements to express Cas9 and guide RNA targeting a locus of interest, and (ii) a separate donor plasmid containing homology regions to a specific locus on the B cell genome as well as DNA sequence encoding a defined integration cassette. For these studies, we used the RA 1 human B cell line for which the SHM mechanisms have been extensively studied.^24,24,40^ We used donor plasmid DNA, p274, and CRISPR/Cas9 plasmid DNA, p276,^41^ that were compatible for integration at the safe harbor H11 locus at chromosome 22 (Chr.22: 31437323 - 31439351) in the human genome. Importantly, this locus is a non-immunoglobulin locus in B cells with no reported instability or recombination^19,42^. We then optimized DNA formulations and electroporation conditions to transfect RA 1 cell line with plasmid DNA (see methods section for details). Next, we generated a homology-directed recombination construct with ∼700 bp homology arms to the H11 integration site. Between these homology arms, we placed an EF1a promoter to drive eGFP expression, and a PGK promoter to drive puromycin resistance cassette (puroR) expression. Finally, we developed recovery and puromycin selection conditions for RA 1 cell lines, where selection was tracked phenotypically by eGFP expression (Supplementary Figure 1 and methods section). The cells were then analyzed by microscopy and fluorescence activated cell sorting (FACS), to determine the levels of eGFP expression. As shown in Figure 1D-E, at least 90% of the total cells demonstrated GFP signals. We also confirmed the integration of the desired recombinant DNA at the targeted integration site in chromosome 22 by isolating the total genomic DNA, followed by PCR analysis (Supplementary Figure 2).

**Figure 1:**
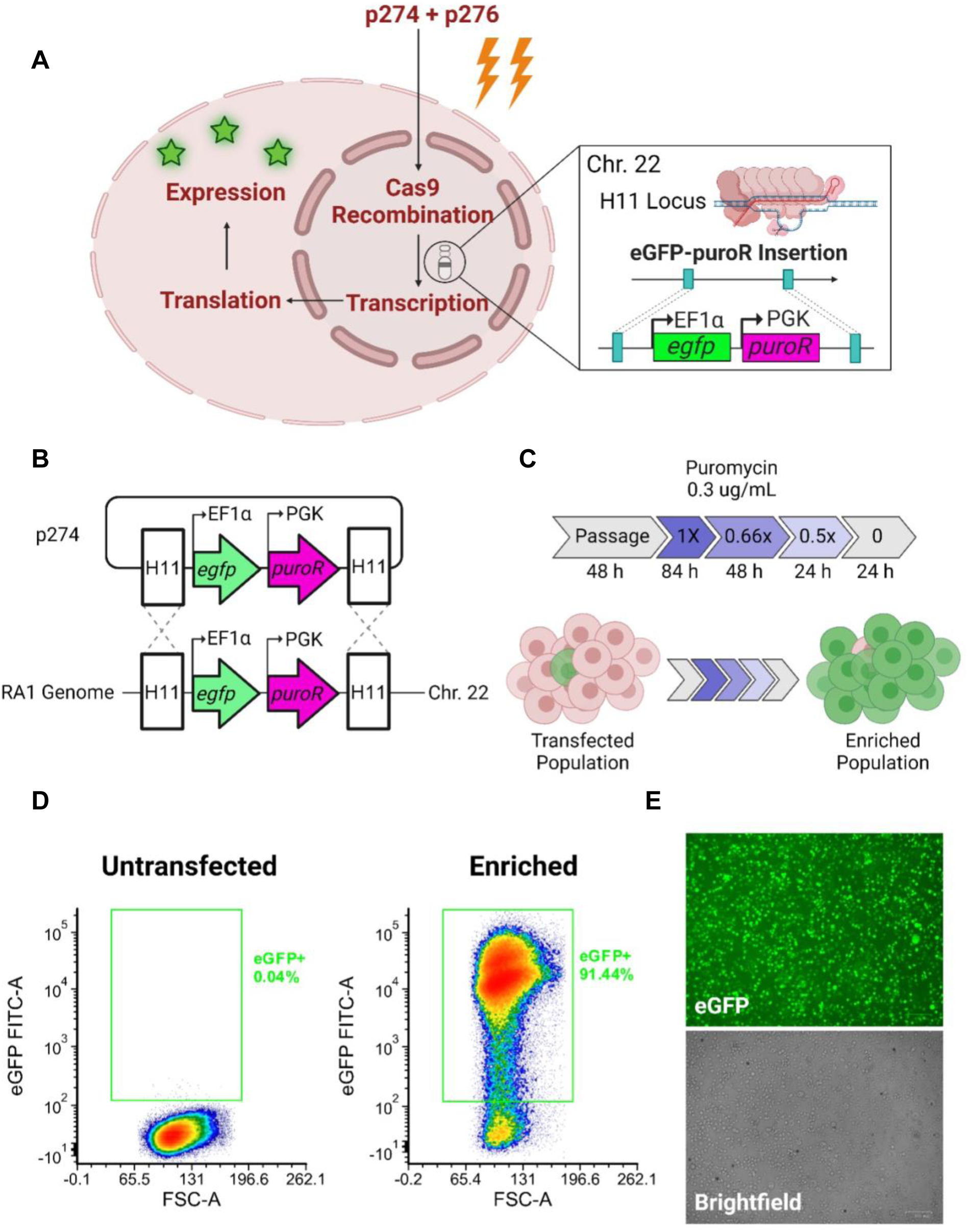
Streamlining RA 1 transfection and enrichment workflow. (**A**) Transfection scheme for RA 1 cell line using a two-plasmid system. (**B**) Integration scheme of homology cassette, following DSB formation by RNA guided Cas9. (**C**) Selection scheme for transfected RA 1 cell line containing the puromycin resistance cassette, puroR. (**D**) FACS density plot of gated eGFP+ cells in untransfected RA 1 cell line and those enriched through puromycin selection; FL1 represents the FITC fluorescence channel. (**E**) Imaging performed on enriched RA 1-274-eGFP cells following puromycin selection. Scale bar indicates 100 µM.

### Engineering human B cell lines to continuously evolve a reporter protein

Having established robust and precise genome editing in B cell lines, our next efforts were directed towards repurposing human B cell lines for continuous directed evolution. As mentioned earlier, viral based integration of fluorescent proteins within the IgH locus facilitated their evolution.^37^ This prior study provided insights into the genomic locus that preferentially underwent SHM due to its ability to attract transcription factors conducive to this process. Tsien and coworkers observed that a series of variants of fluorescent proteins evolved when the corresponding gene was recombined into the genomic Ig heavy-chain locus in chromosome 14 between gene IgHV7-34-1 and IgHV4-34 genes. We hypothesized that the DNA sequence upstream of this integration site at the immunoglobulin locus could play an important role in the evolution experiments performed by Tsien and coworkers. We analyzed the sequences upstream of this integration site and observed the presence of a DNA sequence just upstream (of where the integration site observed by Tsien and coworkers) that had putative recognition site for the Oct transcription factor.^43,44^ Upon comparing sequences that were upstream of IgHV4 locus, we observed the presence of relatively conserved sequences that had the oct motifs (Supplementary Figure 2) which are suggested to be important for transcription from immunoglobulin loci.^44^ Transcription initiation has at the immunoglobulin loci has been linked to SHM.^45,46^ Based on these observations, we investigated if addition of one of these sequences (shown in Supplementary Figure 2) upstream of the EF1a promoter driving eGFP expression could result in SHM at this safe harbor stable H11 locus. Particularly, we chose a 213 base pair fragment from IgH V4-55 (henceforth termed proXIV-1) to test this hypothesis. We also used an eGFP variant, T66I, that lacked detectable fluorescence as a reporter protein (eGFP*). To convert eGFP to eGFP*, we modified the codon from ACC to ATC. We placed proXIV-1 DNA sequence upstream of the gene encoding eGFP* and the EF1a promoter (plasmid DNA, pSB1). If proXIV-1 was able to successfully recruit B cell SHM machinery to this non-immunoglobulin locus where the gene encoding eGFP* was integrated, we would expect mutations to accumulate in this gene, with some of the mutations resulting in the reversion of the non-fluorescent eGFP* to fluorescent eGFP, or variants thereof (Figure 2A-C). This is because the SHM machinery including activity of activation-induced deaminase (AID), a key enzyme in SHM that deaminates cytidine residues to uridine^20^ resulting in a mismatch followed by DNA repair including error-prone base excision repair (BER), which could lead to reversion and other mutations that recover eGFP fluorescence. One of the common outcomes AID mediated deamination followed by repair is the transition of T to C.^47^ Therefore, if this transition mutation occurs in our experiment, it would revert eGFP* back to its original eGFP form (Figure 2A-C). We also made another control DNA construct that had eGFP* driven by the EF1a promoter but that did not contain proXIV-1 (plasmid DNA, pSB2).

**Figure 2:**
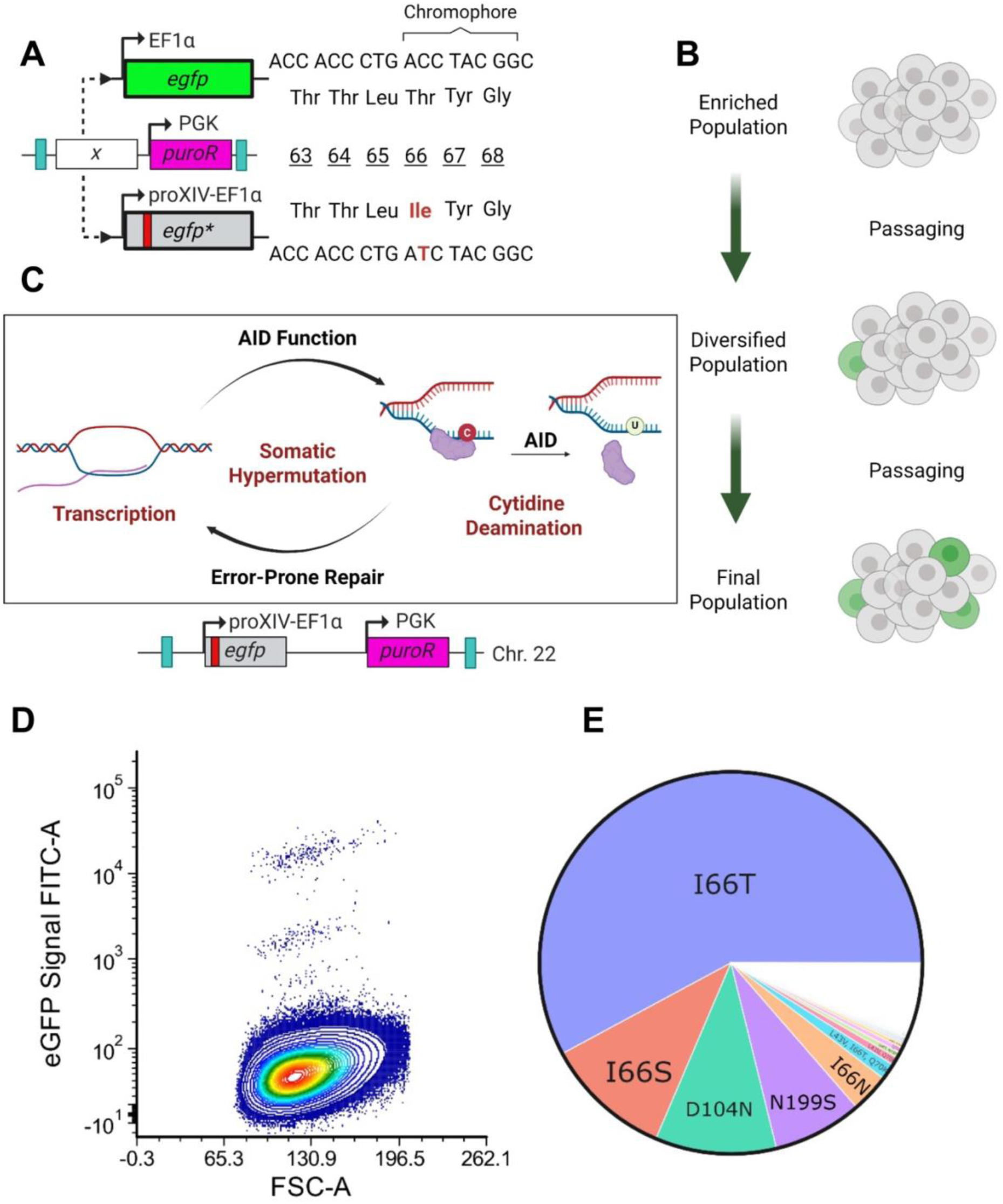
Creation of eGFP knockout, eGFP*, and subsequent restoration of fluorescence through SHM. (**A**) Scheme for introduction of a C to T mutation resulting in change of the Thr residue in the GFP chromophore responsible for fluorescence generation to Ile. (**B**) Scheme for serial passaging and observation of emerging eGFP+ fluorescence. (**C**) Scheme for SHM function in RA 1-proXIV-eGFP* cells. (**D**) FACS density plot showing the appearance of eGFP+ events when RA 1-SB1 cell lines are cultured and passaged for multiple rounds. See Supplementary Figure 4A for microscopy images corresponding to the instances of eGFP+ cells after CODE-HB evolution experiments on eGFP*. See Supplementary Figure 4B for FACS density plots corresponding to RA 1-SB2 cell lines that are cultured and passaged for multiple rounds. See Supplementary Figure 5 for FACS density plots for multiple time points during 30 day evolution experiments. (**E**) Pie chart of NGS analysis showing the emergence of high frequency variants. See Supplementary Figure 6 for more details of mutational profile in sorted eGFP positive samples.

We transfected RA 1 cell line with plasmids p276 and pSB1. We then recovered and selected RA 1 cell line where the DNA segment containing proXIV, EF1a promoter, eGFP, and the puromycin cassette was integrated into chromosome 22, creating the stable RA 1-proXIV-1-eGFP* cell line (RA 1-SB1). For control experiments, we transfected RA 1 cell line with plasmid p274-eGFP*, generating the stable control RA 1-eGFP* cell line, which lacked the proXIV-1 sequence upstream of the eGFP* sequence (RA 1-SB2). For both cell lines we confirmed the integration of the DNA segment at the chromosome 22 locus using PCR analysis (Supplementary Figure 3). Both cell lines were propagated for multiple passages and at each passage, the cell lines were monitored for the appearance of fluorescence signal corresponding to a reverted eGFP variant. After five passages, we performed flow cytometry analysis and observed the appearance of a GFP positive fluorescent events in RA 1-SB1 cell lines that had proXIV-1 sequence upstream of EF1a promoter (Figure 2D, see Supplementary Figure 4 for microscopy images). In contrast, we observed minimal (if any) GFP positive fluorescent events in RA 1-SB2 cell lines that expressed eGFP* but lacked proXIV-1 sequence upstream of EF1a promoter (Supplementary Figure 4). We repeated these evolution experiments in replicates and observed similar results (Supplementary Figure 5). To comprehensively characterize the mutational profiles of these evolution experiments, we performed deep sequencing experiments discussed below.

### Sequencing experiments to elucidate the mutational profile of eGFP*

In order to understand the mutational profile of the evolved eGFP* variants that recovered their fluorescence, we sorted eGFP positive cells using fluorescence-activated cell sorting (FACS), isolated their RNA, and generated the cDNA corresponding to the eGFP* transcript using reverse transcriptase-PCR (RT-PCR) methods. Because we wanted to obtain mutational profiles within a single eGFP* transcript, we performed single molecule long read PacBio sequencing experiments. The sequencing reads (Supplementary Figure 6 and Supplementary Table 5) demonstrated that the introduced mutation in *eGFP* (C197T, nucleotide change) to produce *eGFP*\*, was reverted to its original fluorogenic sequence (T197C, nucleotide change). This reversion accounted for 58.52% of the reads (Supplementary Table 5). Additionally, the second most prevalent mutation in the eGFP positive sorted samples was T197G which converted Ile66 to Ser, effectively recreating the chromophore tripeptide sequence of native GFP, Ser-Tyr-Gly, accounting for 10.19% of mutational reads (Supplementary Table 5).

Other notable mutations included G310A (9.98%) and A596G (7.71%). The G310A substitution changed Asp104 to Asn (Supplementary Figure 6). The A596G substitution changed Asn199 to Ser. Additional substitutions, although less frequent, also contributed to the diversity of amino acid changes. For instance, C208A changed Gln70 to Lys, introducing a positive charge; C127G changed Leu43 to Val; and T50A changed Val17 to Gly, potentially introducing flexibility. It is worth noting that the T197C substitution frequently occurred both independently and in conjunction with other mutations. The co-occurring mutations that were most enriched in the population alongside T197C include G310A, A596G, C208A, and C127G. Additionally, we also observe T197A (Ile66Asn) mutation that does not co-occur with T197C, as they are mutually exclusive due to both being substitutions at the same site. Interestingly, in some cases, T197C was accompanied by as many as three other single point mutations—C127G, C208A, A305G. We also detected A196G/T197C double mutations, which is a previously known GFP variant.^48,49^ A wide range of other mutations (resulting in amino acid changes) were observed in these sequencing experiments (see Supplementary Table 5 and Supplementary Data File 1), demonstrating the remarkable scope of mutations that can be accrued within the gene using this approach.

To characterize the mutational profile across the entire population of unselected cells, we performed sequencing experiments on the unsorted naïve pool of cells that had undergone multiple rounds of cell culturing and passaging which would result in the accumulation of mutations. Particularly, we propagated the RA 1-SB1 cell lines for multiple passages. Then we isolated genomic DNA from an early passage (passage 3 after generating the cell line) and a late passage (passage 8 after generating the cell line), PCR amplified gene encoding eGFP* and sequenced the PCR products using Illumina sequencing to obtain higher sequencing depth. We performed comprehensive mutational analysis as shown in Figure 3 and Supplementary Figures 7-11. A heat map showing genomic DNA mutations in the at passage 3 is shown in the Supplementary Figure 7 and another heat map showing new mutations at passage 8 is shown in the Figure 3D. As can be seen from these mutational heat maps, we observe a broad mutational spectrum in both earlier and later passages with a wide range of substitution pattern. A bar chart depicting mutational bias in this experiment is shown in the Figure 3D. Additionally, since we had sequencing data for multiple timepoints of evolution in replicates, we were able to plot representative examples of some of the interesting evolutionary trajectories (Figure 3A). As expected, one of the key mutants was T197C mutation, which reverted the mutated eGFP*. In addition to this, we observed unique mutations in passage 8 that were not observed in earlier passage 3 (Figure 3A). We also observed a series of stacked mutations—several of the representative phylogenetic trees are shown in Figure 3A. Finally, in addition to substitution mutations, we also observed the presence of deletions and insertions (Supplementary Figure 8), albeit at a relatively lower frequency, which is also consistent with previous studies on SHM in B cell lines.^24^ We also performed eGFP* evolution experiment in replicate, and the mutational profiles of the replicate experiment are shown in Supplementary Figure 5. Further, we also isolated total RNA from passage 3 and passage 8 cells, generated cDNA, PCR amplified cDNA corresponding to the evolved eGFP* transcripts and sequenced the PCR products using Illumina sequencing. The mutational profiles of the replicate experiment are shown in Supplementary Figures 9-10. In each of the cases we observed similar mutational profiles and breadth (Supplementary Figures 9-10). All these observations are consistent with those previously reported for SHM in B cells and B cell lines.^47,50^ We call this continuous directed evolution platform as CODE-HB.

**Figure 3.**
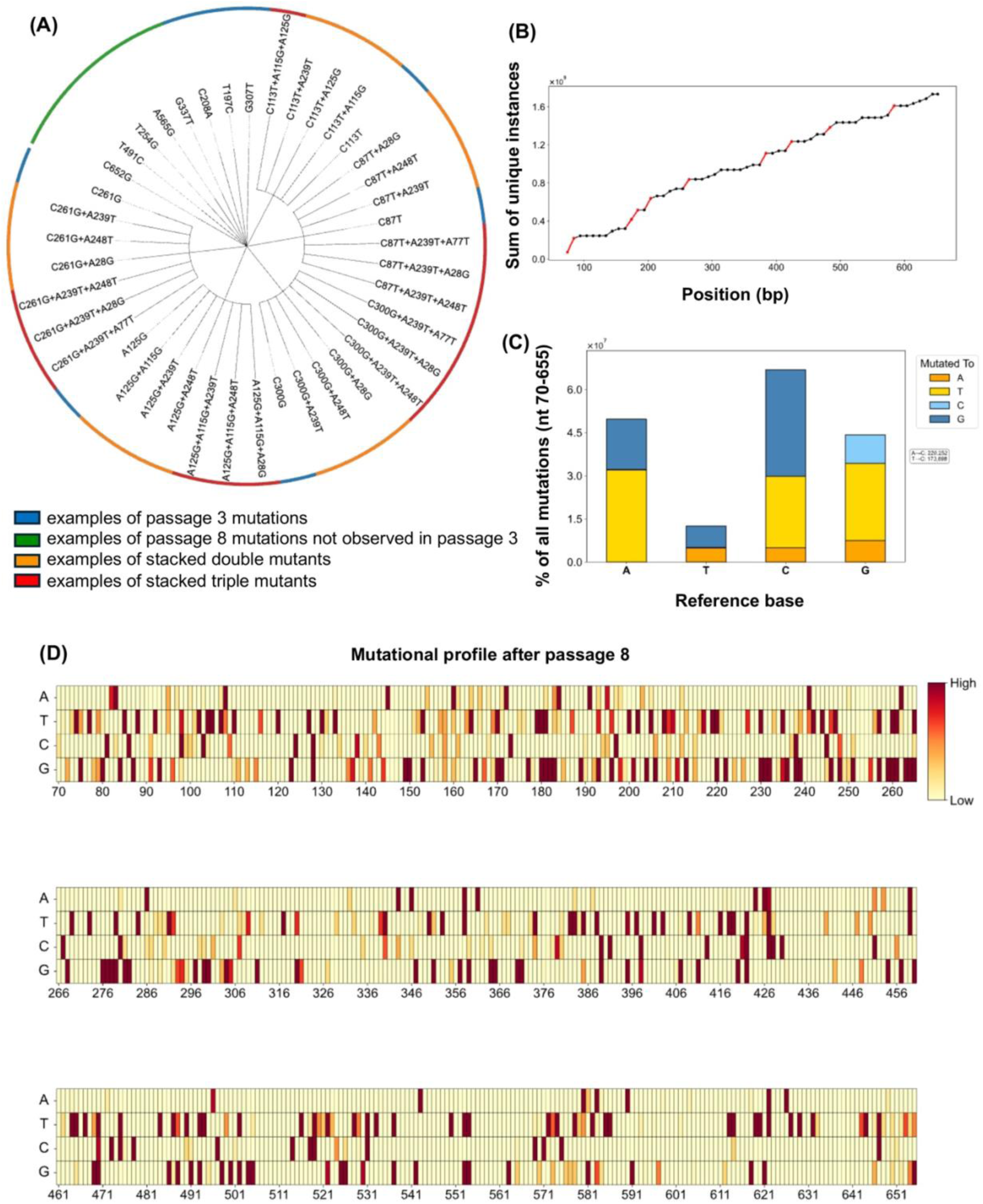
Mutational analysis of unsorted, naïve pool RA1 cells engineered to express eGFP* after iterative rounds of passaging. **(A)** Phylogenetic tree showing representative nucleotide level mutations in the eGFP* locus. Mutations highlighted in blue initially occur during an early passage and then accumulate associated mutations in late passages (orange and red labels). Mutations labeled green are exclusive to the late passage and do not occur in the early passage. For examples of more than 2 stacked mutations, insertions and deletions see Supplementary Figure 6 and Supplementary Figure 8. **(B)** Integral plot of occurrences over the sequence length of eGFP*. **(C)** Bar plot comparison of nucleotide mutations in the eGFP* sequence at a late passage. Mutations are plotted as the percentage of total reads/sum of all mutations. **(D)** Heatmap showing density of mutations across the eGFP* sequence after 8 passages.

### Mutational bias during eGFP* evolution experiments using CODE-HB

In order to characterize the mutational profile and biases, we next analyzed the mutations observed during the eGFP* evolution experiments. First, we analyzed the sequencing data from naïve pool of unsorted cells that were passaged for multiple passages thereby resulting in the accumulation of mutations. Based on these sequencing profiles, we characterized the mutational biases for each of the bases. We observed a broad substitution pattern (Figure 3C) that is consistent with SHM mediated mutations.^24,47^ As expected, we observed that cytosine (C) bases were mutated to thymine (T), guanine (G) and adenine (A) at relatively high frequency. This can be attributed to AID mediated cytidine deamination followed by repair. Similarly, we observed that the G bases were substituted with A, T and C bases with considerable frequency. We observed that A bases were substituted with G and T bases at moderate to high frequency and C bases at a lower frequency. And T bases were substituted at a relatively lesser frequency with either A or G bases. A plot of the cumulative occurrences of mutations is shown in Figure 3B. This trend line exhibited sections with increases in slope, corresponding to regions with higher number of mutations. As mentioned earlier, in addition to substitution mutations, we also observed the presence of deletions and insertions (Supplementary Figure 8). In the eGFP positive sorted samples, we similarly observed a broad mutational profile (Supplementary Figure 6). A plot of the cumulative occurrences of mutations for these sorted samples is shown in the Supplementary Figure 6). Also, in the eGFP positive sorted samples, the observed mutational profiles for each of the nucleotide bases is shown in the Supplementary Figure 6C. When the corresponding amino acid changes were mapped onto the protein structure, they localized to a lateral portion of the eGFP barrel, proximal to and including the chromophore (Supplementary Figure 6E). Importantly, these mutational profiles, preferences and spread were similar to those previously characterized SHM in B cells and B cell lines^47,50^, indicating that the mutations in CODE-HB are predominantly due to B cell SHM mechanisms. It is also important to note here that SHM mediated mutational profiles are highly sequence dependent and could vary based on the hot spots or cold spots for SHM which remains an active area of investigation.^47,51^

### Mechanistic characterization to elucidate the DNA sequences that can be used to recruit SHM machinery

The findings described above, led us to investigate if we could identify other sequences like proXIV-1 that are capable of driving mutagenesis in CODE-HB. As described above, when we analyzed the sequences from upstream of IgHV4 locus, we observed the presence several sequences that were similar to proXIV-1 (Supplementary Figure 2A-C). We investigated if the addition of one of these sequences upstream of the eGFP* sequence could result in mutagenesis in our CODE-HB system. Particularly, we chose a 213 base pair fragment from IgH V4-34 (henceforth termed proXIV-2) and replace proXIV-1 in pSB1 with proXIV-2 (resulting plasmid was named pSB5). We transfected pSB5 along with p276 in RA 1 cell line to generate RA 1-SB5 cell line which expresses non-fluorescent eGFP*. Next, we continued to culture and passage this cell line and analyzed them by flow cytometry analysis to determine if GFP positive signals appear over time. Remarkably, we detected the presence of significant GFP positive events (Figure 4A, 4C, 4D and Supplementary Figure 5A-B) in RA 1-SB5 cell line and minimal GFP positive events (if any) in RA 1-SB2 cell lines that lack proXIV sequences. This clearly suggested that similar to proXIV-1 sequence, the proXIV-2 sequence was also able to drive mutagenesis in CODE-HB. A sequence comparison highlighting the conserved residues in proXIV-1 and proXIV-2 is shown in Supplementary Figure 2D.

**Figure 4:**
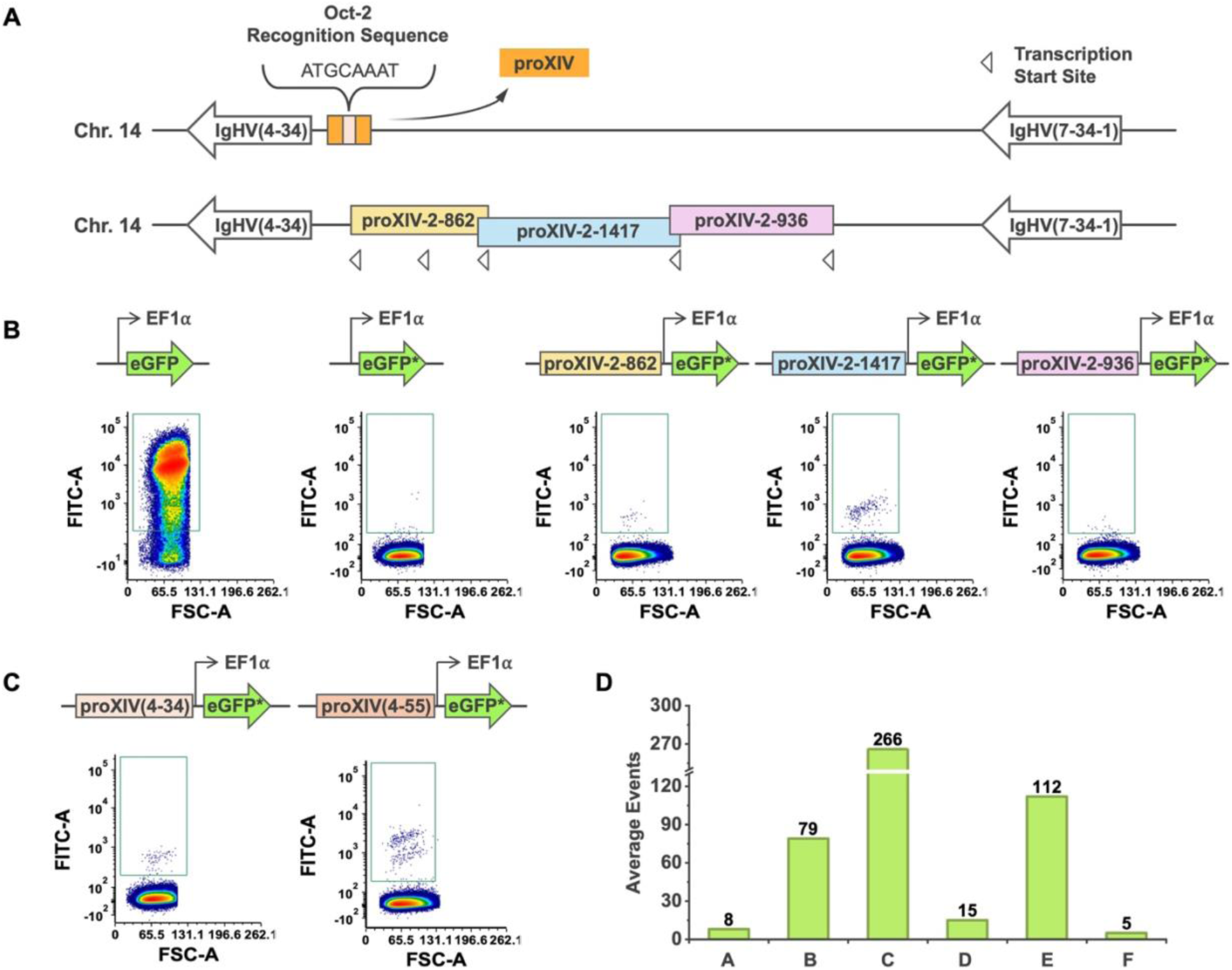
Mechanistic studies that resulted in the identification of additional SHM recruiting sequences. (**A**) Genomic locus corresponding to IgHV(4-34) from where proXIV-2, proXIV-2-862, proXIV-2-1417 and proXIV-2-936. (**B**) Flow cytometry analysis density plots of the transfected RA1 cells. (**C**) FACS density plots of evolved cells that have either proXIV-2 obtained from IgHV(4-34) or proXIV-1 obtained from IgHV(4-55) upstream of eGFP* sequence. (**D**) Bar chart representing the average event counts per 500,000 events for each sequence at day 13. A: EF1a-eGFP*, B: proXIV(4-34), which is proXIV-2, C: proXIV(4-55), which is proXIV-1, D: proXIV-2-862-EF1a-eGFP*, E: proXIV-2-1417-EF1a-eGFP*, F: proXIV-2-936-EF1a-eGFP*

To understand if other DNA fragments further upstream of proXIV-2 sequence could drive mutagenesis in CODE-HB, we took around 2500 base pair sequence upstream of proXIV-2 and generated three distinct DNA fragments from this sequence—proXIV-2-862, proXIV-2-1417 and proXIV-2-936 (Figure 4A-C); note that each of these fragments had a potential transcription start site and proXIV-2-862 sequence contains proXIV-2 sequence within it. We replaced proXIV-2 in pSB5 with either of the three DNA fragments to generate plasmids pSZ1, pSZ2 and pSZ3 (Supplementary Table 3). We individually transfected either of these plasmids along with p276 in RA 1 cell lines to generate RA 1-SZ1, RA 1-SZ2 and RA 1-SZ3 cell lines. In each case we continued to culture and passage these cell lines and analyzed them by flow cytometry analysis to determine if GFP positive signals appear over time. We observed the presence of GFP positive signals in RA 1-SZ1 and RA 1-SZ2 cell lines and minimal GFP positive signals (if any) in RA 1-SZ3 cell lines (Figure 4C-D). This suggested that both proXIV-2-862 and proXIV-2-1417 were able to drive mutagenesis in CODE-HB but proXIV-2-936 was not able to drive mutagenesis in CODE-HB. Taken together, these experiments allowed us to identify four different DNA sequences that were able to drive mutagenesis in our CODE-HB system. A representative sequence comparison of these sequences is shown in Supplementary Figure 2. Among all sequences we tested, proXIV-1 sequence from IgH V4-55 consistently yielded the highest frequency of eGFP-positive events and was therefore selected for further evolution experiments.

### Developing facile F_ab_ surface display platform in human cells

Surface display technologies like phage display, yeast display and mRNA display have proved to be crucial for performing selections in directed evolution campaigns.^4,52–57^ These approaches have been especially applied to develop highly successful biologics like therapeutic antibodies. Therefore, in order to expand the scope of CODE-HB to develop biologics, we developed a human B cell surface display platform. Particularly, we envisioned combining our human B cell surface display platform with CODE-HB to evolve potent antibodies targeting emerging strains of avian influenza that are a threat to human health and world economy (e.g., H5N1, 2024). To this end, we began by selecting a fragment antigen binding region (F_ab_) of a well characterized broadly neutralizing antibody, CR9114.^58^ Following a method inspired by Moffett et al.^36^, a single-chain antibody binding fragment (F_ab_) was engineered by fusing the light and heavy chains with a 60 amino acid glycine-serine repeat (GlySer) linker. This fusion construct was further modified for surface display by adding a membrane localization sequence at the N-terminus and a major histocompatibility complex (MHC) I transmembrane helix at the C-terminus, facilitating its stable membrane tethering for cell surface presentation.^59^ Furthermore, we inserted a FLAG-tag between the constant region of the heavy chain and the transmembrane helix to facilitate detection of construct expression independently from antigen binding (Supplementary Figure 12). The DNA encoding this chimeric protein sequence (F_ab_-surface display protein) was subcloned into p274 where the expression of the F_ab_-surface display protein was driven by the EF1a promoter. The resulting plasmid was called pSB3. To evaluate whether this surface display platform was functional in human cell lines, we transfected the Expi 293 cell line with plasmid pSB3. The expression of the F_ab_ was evaluated by binding of APC-conjugated Anti-FLAG antibodies to these Expi 293 cells. The binding of the surface displayed F_ab_ to the reporter hemagglutinin A, H1, was evaluated by using the fluorescent signal of a C-terminally fused mCherry. Analysis of expression and binding was performed using flow cytometry. To ensure that the emissions of the two fluorophores were distinctly separated, mCherry was excited with a 561 nm laser and detected through a 613/18 nm bandpass filter, while APC was excited with a 640 nm laser and detected using a 660/10 nm bandpass filter. Using this approach, we were able to distinctly delineate expression and binding due to non-overlapping fluorescence signals corresponding to binding (mCherry) and expression (APC) (Supplementary Figure 13). This distinction facilitated accurate and independent evaluations of both expression levels and binding efficacy at the single cell level. Furthermore, the data revealed that the population of CR9114 F_ab_, when displayed on the cell surface, maintained its functional integrity, demonstrating effective binding across a spectrum of hemagglutinin subtypes, including H3 and H7 (Supplementary Figure 13C).

Next, we validated that this surface display platform was functional in B cell lines as well. We transfected plasmid pSB3 and p276 in RA 1 B cell lines, recovered them and selected them as before to generate RA-SB3 cell lines. These RA-SB3 cell lines were analyzed for expression of F_ab_-surface display protein using APC-conjugated Anti-FLAG antibody and the binding of the surface displayed F_ab_ to H1 was evaluated using an H1-mCherry fusion protein (Figure 5A-B). We observed robust expression and H1 binding for these RA 1 mutant cell lines displaying CR9114 F_ab_ (Figure 5C).

**Figure 5.**
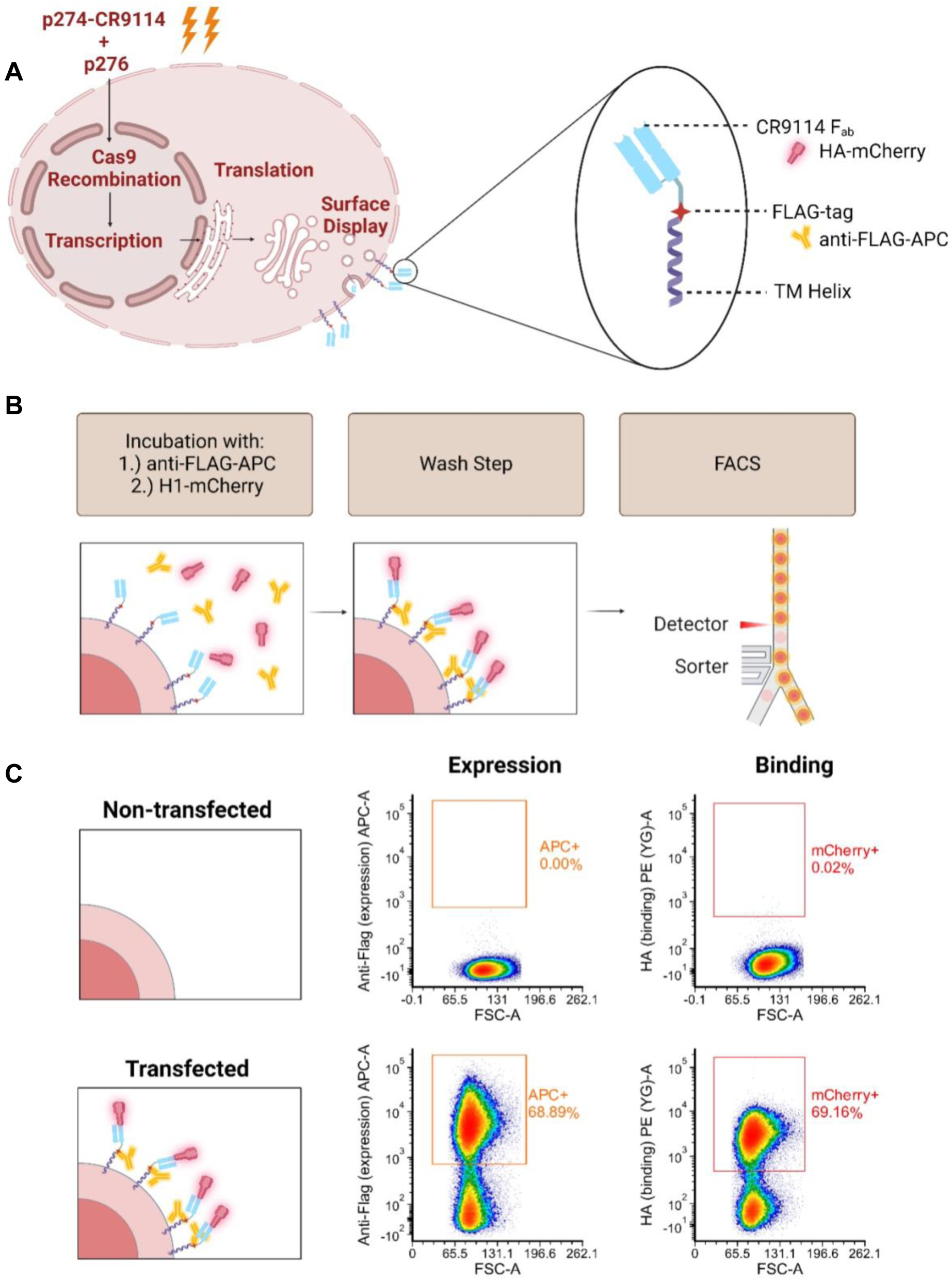
Development of a surface display platform for CODE-HB. (**A**) Scheme of F_ab_ display on the surface of RA 1 cell line, utilizing fluorescent HA-mCherry and fluorescent anti-FLAG-APC mAbs to parameterize the chimeric protein into separate binding and expression variables. (**B**) Scheme of FACS experiment. (**C**) Density FACS plots of enriched RA 1-CR9114 cells stained with anti-FLAG-APC and H1-mCherry.

### Evolving antibody fragments by combining F_ab_ surface display with CODE-HB

Next, we investigated if the surface displayed F_ab_ sequences could be evolved using CODE-HB. Particularly, we investigated if the CR9114 F_ab_ can be evolved to exhibit a greater degree of binding towards hemagglutinin A (HA) from emerging strains of avian influenza (H5); identifying H5 selective antibodies is especially important in the context of recent zoonotic spread of H5N1 strains of avian influenza.^39^ We started by inserting the CR9114 F_ab_ surface display protein into a B cell integration vector containing the proXIV enhancer sequence to generate plasmid pSB4 (Figure 6A). We transfected plasmid pSB4 and p276 into RA 1, recovered cells and selected them as before to generate RA 1-SB4. We also utilized the control plasmid pSB3 that was identical to pSB4 but lacked proXIV. Using this integration plasmid, we generated a control cell line RA 1-SB3. We then performed iterative rounds of cell passaging, analysis and selections to determine if we can evolve F_ab_ sequences with increased binding towards H5 (Figure 6B). To refine the selection process for cells exhibiting increased affinity towards H5, we performed flow cytometry analysis to evaluate the ratios of binding to expression (Figure 6C-D). We used a control cell line, RA 1-SB3 expressing CR9114 F_ab_ that lacked proXIV sequence upstream and performed H5 binding analysis using flow cytometry (Supplementary Figure 13D). This allowed us to determine basal level of binding to expression ratio which allowed us to establish a “High Binding” gate for events surpassing expected binding levels based on expression alone (Figure 5C). Cells that exceeded this benchmark within the proXIV-inclusive sample, yet not present in the controls lacking proXIV sequence, were posited as having undergone favorable somatic hypermutations within their recombinant CR9114 construct sequences.

**Figure 6.**
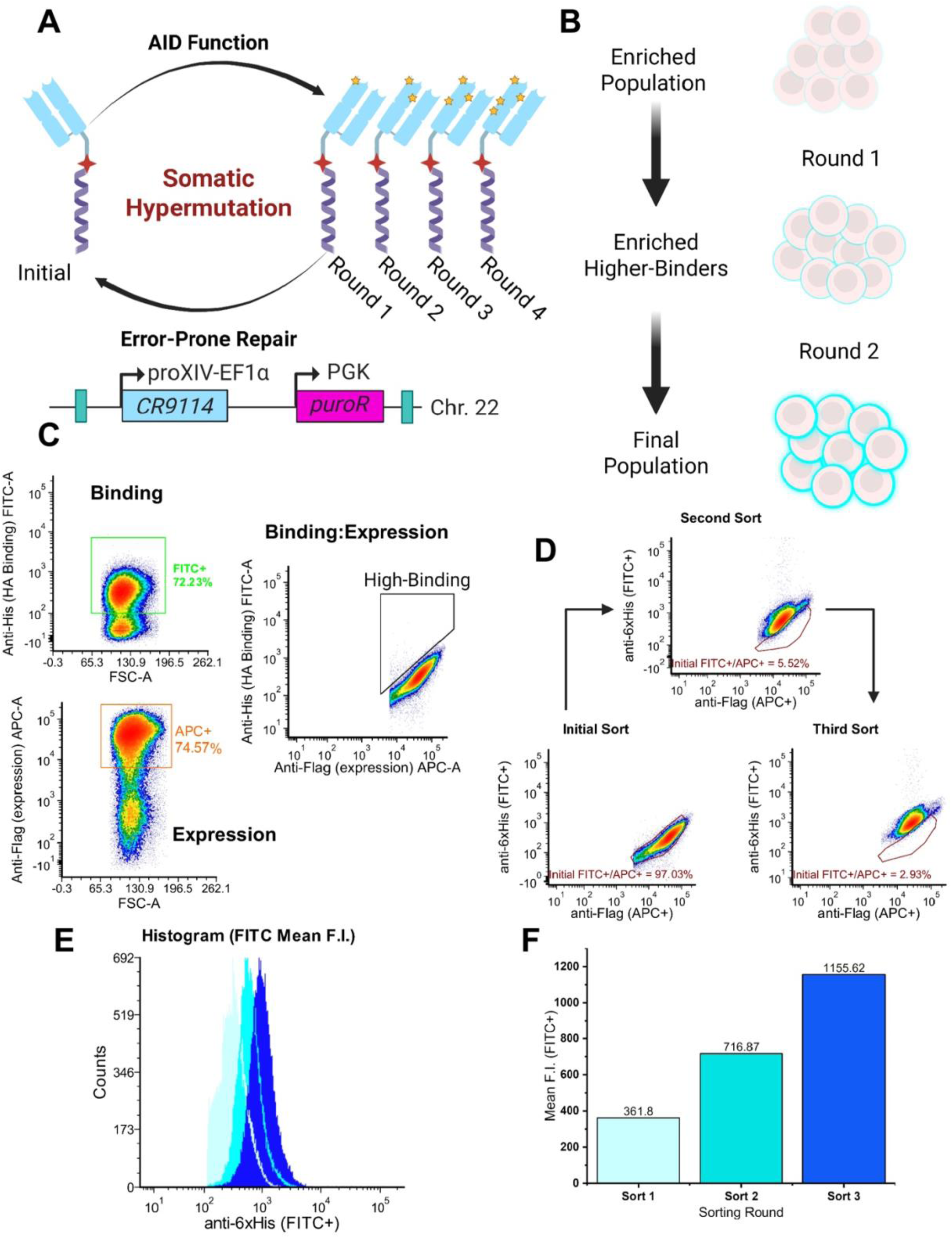
Workflow for the continuous directed evolution of higher-binding CR9114 variants. (**A**) Schematic for the incorporation of mutations into the gene encoding CR9114 F_ab_ by iterative cycles of SHM. (**B**) Scheme for the iterative cycle of enriching cells binding to H5-mCherry, followed by diversification of the CR9114 gene through SHM that occurs during transcription, followed by selection of higher-binding variants and subsequent enrichment. (**C**) FACS density plots of the RA 1-SB3 gating strategy: expression (APC+) and binding to H5 (FITC+) channels, When binding versus expression is plotted, this yields a linear relationship (binding:expression plot) from which higher-binding variants may be gated and sorted for growth. (**D**) FACS density plots for iterative sorting and passaging (expression vs binding) for continuously evolving RA 1-SB3 cell lines. For this verification experiment His-tagged H5 was used and FITC-conjugated anti-His antibody was used to detect binding of cells to His-tagged H5. (**E**) Histogram representing each iterative sort in terms of H5-binding. (**F**) Bar chart representing mean F.I. of the histogram.

Upon utilizing the “High Binding” gate for selection, an enrichment of approximately 40,000 cells was achieved that were cultivated to saturation. Subsequent iterations of FACS sorting and analysis of this population highlighted an intriguing divergence from the control group’s fluorescence profile, characterized by a decreasing mean fluorescence intensity in the expression channel, contrasted with a consistent intensity in the binding channel (Supplementary Figure 14A). This unexpected trend prompted a deeper molecular investigation, leading to the isolation of RNA from the sorted population and Sanger sequencing of the CR9114 amplicon. The sequencing results revealed a prevalent mutation that specifically resulted in the conversion of Tyr532 to Phe532 (TAC to TTC), adjacent to the FLAG-tag region, suggesting that a targeted selection pressure in this area resulted in a phenotype that could be detected (Supplementary Figure 14B). These observations demonstrated that a phenotype can be linked to the genotype and can be evolved continuously by using CODE-HB.

In addition to this population, we also observed a dynamic population of high binders and low binders as compared to the control group (Figure 6D, Supplementary Figure 13D, Supplementary Figure 14C). The enrichment process showed a marked improvement in target population isolation across successive sorting steps. Initially, the percentage of events that escaped the control wild-type binding gating was 2.97%. This number increased substantially to 94.48% after the second sort, and further to 97.07% following the third sort, demonstrating progressive refinement of the target cell population (Supplementary Figure 15). Correspondingly, the histogram data illustrated a significant rise in the mean fluorescence intensity (F.I.) of the H5-binding channel (FITC+), with values of 361.8, 716.87, and 1155.62 in the initial, second, and third sorts, respectively (Figure 6E-F). For this verification experiment His-tagged H5 was used and FITC-conjugated anti-His antibody was used to detect binding of cells to His-tagged H5. Post-sorting, the cells were expanded, and their RNA was isolated for subsequent conversion to cDNA.

### Single-molecule long read sequencing to elucidate the mutational profiles of evolved F_ab_

Once the cDNA of the transcript corresponding to the F_ab_-surface display protein was generated, long read single molecule sequencing was performed (Figure 7A). Third generation sequencing generated a total of 642,858 reads, of which 471,562 met the expression threshold based on published PacBio error rates (15 observations).^60^

**Figure 7.**
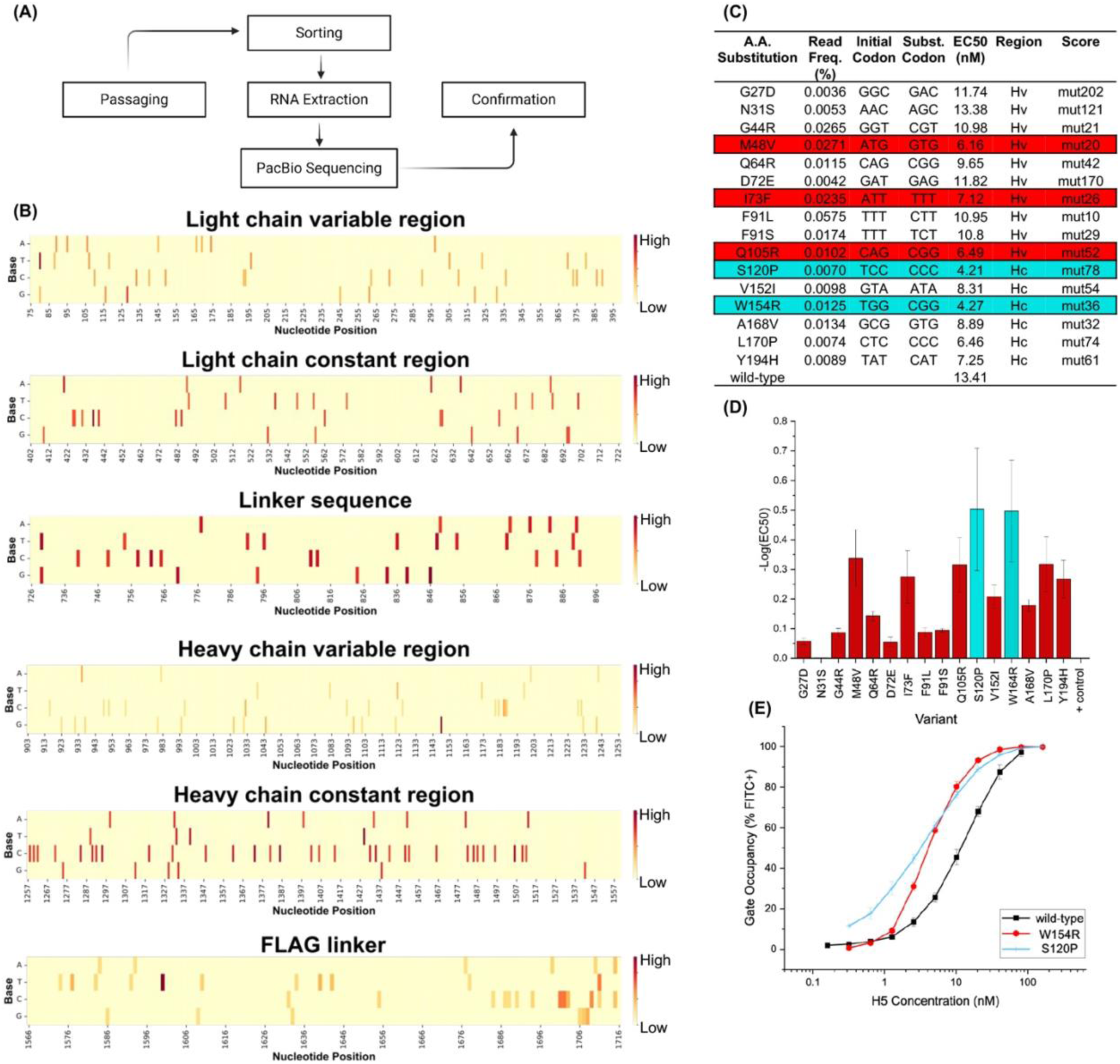
Isolation of enriched CR9114 variants obtained by SHM and their experimental validation. (**A**) Experimental scheme for identification and characterizing higher-binding variants. (**B**) Heatmap showing density of mutations across the surface displayed CR9114 F_ab_ after 3 rounds of sorting and enrichment. See Supplementary Figure 19 for mutational profile after 1 round of sorting and enrichment. See supplementary Figure 20 for examples of insertion deletion mutations in evolved CR9114 sequences. (**C**) Table of read frequencies for the tested variants alongside their substituted codons, encoded amino acids, and EC50 values; variants were named based on the amino acid substitution found in the heavy chain of CR9114 and numbered based on the read frequency of their unique nucleotide substitution in the sequencing data - lower numbers indicate a higher degree enrichment. Point mutations listed for the F_ab_ fragment also include at least one additional mutation, specifically A1601T and/or A1149G. (**D**) Bar chart representing the EC50 values of the 16 CR9114 heavy chain variants (red: variants in heavy variable region, cyan: variants in heavy constant region). (**E**) Titration curve for the two high-affinity CR9114 variants, W461R and S427P, relative to the CR9114 wild-type control.

Further analysis focused on heavy chain variants encompassing both the variable (Hv) and constant (Hc) regions, due to the heavy chain-only binding mode of CR9114 to HA as indicated by the crystal structure (PDB: 4CQI).^61^ Unique reads from the top 200 variants were shortlisted, resulting in a final set of 16 variants that included both variable and constant regions of the heavy chain. Notably, these variants represented only 0.25% of the total reads. Analysis of these variants revealed a variety of nucleotide substitutions, including transitions, transversions, insertions, and deletions (Figure 7B, Figure 7C, Supplementary Figures 16-20 and Supplementary Data File 2). Approximately 22% of the unique read sequences introduced insertion or deletion mutations when aligned to the reference sequence. Of the remaining sequences, 82% corresponded to substitution mutations comprising of at least two mutations: A1149G and A1601T. The first mutation, A1149G, was a silent mutation that altered the codon of a glutamic acid residue without changing the amino acid sequence. The second mutation, A1601T, resulted in an amino acid change from tyrosine to phenylalanine within the FLAG tag sequence (tAc→tTc, Tyr→Phe). This mutation results in the change of a tyrosine residue within the FLAG tag to phenylalanine which potentially reduces antibody recognition there by increasing the binding:expression ratio. The subsequent multiple sequence alignment included 208 unique sequences. In addition to these mutations, we see a series of stacked mutations on top of A1601T mutant, for example we observe A1149G/A1601T mutant, A1601T/C107A/A1149G mutant, C107A/G119T/A1149G/A1601T mutant. Several representative examples of stacked mutations are shown in Supplementary Figure 17-18). Beyond these variants, a variety of other mutations were present across the F_ab_ sequence with varying frequency (Supplementary Data File 2). A heat map of CR9114 mutational profile after the first round of sorting and enrichment is shown in Supplementatry Figure 19 and another heat map of CR9114 mutational profile after the third round of sorting and enrichment is shown in Figure 7B. As can be seen from these data, there is a significant diversification of CR9114 sequences after multiple rounds of evolution and enrichment. Following the nucleotide-level analysis, remaining reads were translated, and protein level variations were analyzed (Supplementary Figure 18). In addition to this, examples of insertion and deletion mutations in the evolved variants of CR9114 is shown in the Supplementary Figure 24. Based on these analyses, several variants were shortlisted for validation and further characterization as described in the sections below.

### Validating CR9114-F_ab_ variants for increased binding towards hemagglutinin H5

Next, we identified and validated F_ab_ sequences that demonstrated enhanced binding. We only selected a subset of variants from the sequencing data based on the on mutational frequency and/or mutational positioning (e.g., in the CDR loops). Functional characterization of these variants involved determining their EC50 values through hemagglutinin H5 titration against the surface-displayed form (Figure 7 and Supplementary Figures 21-22). Overall, the variants exhibited lower EC50 values compared to the CR9114 wild-type control, reflecting enhanced binding affinities in all the cases we tested. Among the variants we tested, two variants, VH S120P and VH W154R, demonstrated the lowest EC50 values of around 4.2 nM (Figure 7C-D). These values were approximately three times lower than that of the CR9114 wild-type control, highlighting the superior, yet modestly improved, binding affinity of these selected variants. Interestingly, the two variants with the lowest EC50 values, VH S120P and VH W154R, both contained mutations in the heavy chain constant region. This finding is particularly noteworthy as it suggested that alterations in the constant region, typically associated with structural and stability roles rather than direct antigen binding, can significantly influence the binding affinity of the antibody by optimizing an already-evolved binding interface. This observation highlights the potential for targeted mutations in this constant region to improve therapeutic antibody efficacy. The mutant VH W154R was tested for retention of binding towards other hemagglutinin variants (H1, H3, H5) and was found to not only retain but also exhibit more favorable EC50 values than the wild-type CR9114 (Supplementary Figure 22).

In addition to these observations, variants in the heavy variable region that demonstrated enhanced binding relative to the controls were also detected (Figure 7C-D, Supplementary Figure 21). The VH I73F variant of CR9114 introduced a point mutation within a critical complementarity-determining region (CDR) loop on the heavy chain, which interfaces directly with the hemagglutinin H5 antigen (crystal structure PDB: 4CQI). The VH M48V variant of the CR9114 antibody is another point mutation within the beta sheet of the heavy chain variable region, a critical area for maintaining the antibody’s structural integrity. The VH Q65R variant of the CR9114 antibody involved a point mutation within a flexible loop of the heavy chain variable region, situated near a hydrogen bond network in CR9114 (PDB: 4CQI, Supplementary Figure 23).

### Testing the evolved variants of CR9114 using viral neutralization assays

To determine if the evolved CR9114 variants still retained viral neutralization potential, we first recombinantly produced CR9114 variants (Supplementary Figure 24). We then performed ELISA assays and verified binding to H5 (Supplementary Figure 25). Next, to determine if the antibody variants are able to neutralize the influenza virus, we performed viral neutralization assays with biosafety level two compatible influenza virus, influenza A/Puerto Rico/1934 (H1N1). To quantify the capacity of our CR9114 variants to block receptor engagement, we performed hemagglutination-inhibition (HAI) assays using turkey erythrocytes and influenza A/Puerto Rico/1934 (H1N1) (Figure 8A). Wild-type CR9114 exhibited an HAI₅₀ of 50 µg/mL, consistent with its broadly neutralizing profile. Each of the heavy-chain variable-region mutants—designed to probe fine adjustments in the antigen-contact interface—yielded HAI₅₀ values statistically indistinguishable from the parental antibody (mean HAI₅₀ ≈ 50-100 µg/mL; n = 3). In stark contrast, the constant-region substitution W154R conferred a significant four-fold enhancement in inhibitory potency: the W154R variant achieved 50% inhibition of hemagglutination at just 12.5 µg/mL (Figure 8B). This improvement implies that modifications outside of the antigen-binding site can influence the functional avidity the ability of CR9114.

**Figure 8:**
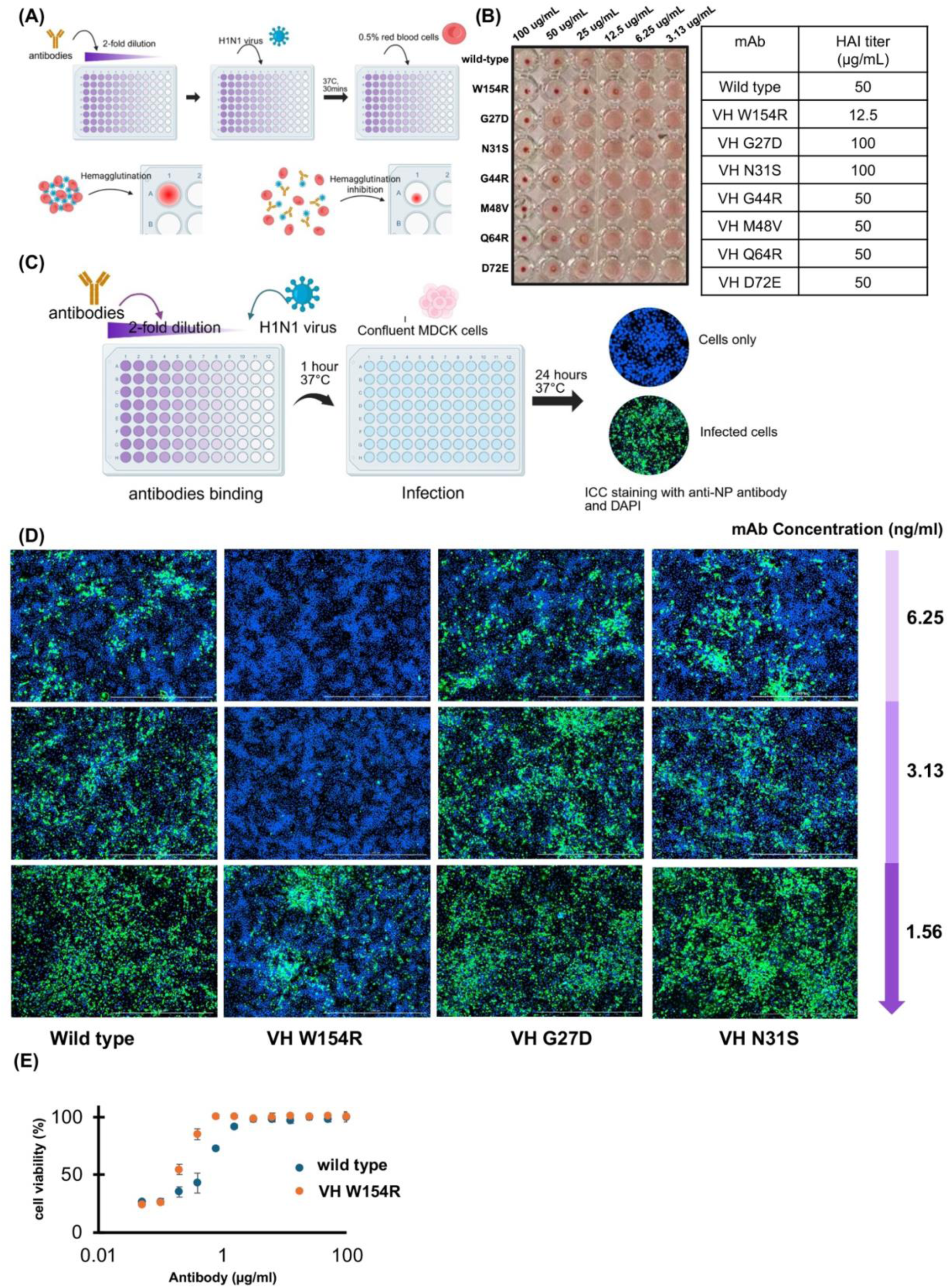
Viral neutralization validation of CR9114 mutants of interest generated using CODE-HB. (**A**) Schematic of the hemagglutination-inhibition (HAI) assay workflow. (**B**) Imaging data from the hemagglutination assay for CR9114 variants that exhibited ELISA binding, with corresponding HAI titers required for inhibition. (**C**) Workflow schematic for influenza neutralization assays. (**D**) Imaging data showing neutralization of A/Puerto Rico/1934 (H1N1) following incubation with purified variants CR9114 mAbs. (**E**) Antibody dose response experiments for viral neutralization assays demonstrating improved efficacy of W154R variant of CR9114 as compared to wild type CR9114.

To determine whether the W154R variant’s increased HAI activity translates to stronger blockade of viral entry, we next assessed viral neutralization in a cell-based infection assay. Influenza A/Puerto Rico/1934 virions were pre-incubated with serial dilutions of each mAb before inoculation onto MDCK monolayers. Wild-type CR9114 required concentrations above 100 nM to achieve >90% reduction in cytopathic effect, whereas the W461R variant maintained equivalent neutralizing efficacy at 10 nM—well within physiologically attainable antibody titers (Figure 8C-E). Thus, in addition to achieving high binding for H5, this variant was also able to neutralize H1N1 in our assays. To understand how they key variant W154R alters the structure of CR9114 antibody, we computationally substituted the W154 with R in the crystal structure of the CR9114 Fab in complex with the H5 hemagglutinin and performed energy minimization (Supplementary Figure 26). While CR9114 exclusively binds to H5 HA using its CDRH loops, previous studies suggest that the binding contributions made by the light chain also play an important role in antigen binding.^62^ Our analysis suggested a significant bond shortening for the preserved hydrogen bond between heavy chain H164 and light chain Q168; this light chain residue is connected to a broader hydrogen bond network in the L_c_ region that interfaces with the L_v_ region and potentially transfers this stability to the light chain residue R31, which stabilizes the configuration of CDRH residues.

Together, these data reveal that mutations in the Fc-proximal domain can markedly improve functional potency of CR9114 against H1N1. These experiments demonstrate the utility of CODE-HB for rapid evolution and engineering of high-affinity antibodies.

### Leveraging broad mutational profile of CODE-HB for evolution experiments

To showcase how the broad mutational profile of CODE-HB can we used for antibody engineering, we present another antibody evolution campaign. This evolution campaign starts off with previously identified 047-09_1A02 antibody that shows high binding to influenza H1/California/2009 but lower binding to influenza H1/Michigan/2015 H1 hemagglutinins.^63^ To enhance the binding 047-09_1A02 to influenza H1/Michigan/2015, we employed our CODE-HB system (Figure 9A). Particularly, we engineered RA 1 cells to display 047-09_1A02 F_ab_ on the surface (RA 1-DJO1 cell line) and performed evolution experiments to identify potent binders of H1/Michigan/2015 (Figure 9B). After sequential rounds of evolution and enrichment, we sequenced the 047-09_1A02 F_ab_ transcripts to determine key mutations that result in enhanced binding to H1/Michigan/2015. As expected, we observed a wide range of substitution mutations (Figure 9C, D). In addition to this, we also observed a seven–amino-acid deletion within the sc60 linker. Additionally, we also observed several substitution mutations on top of the 7 amino acid deletion (Figure 9D), including mutations in the region corresponding to CDR loops. By constructing and assaying variants bearing each alteration alone or in combination, we discovered that excising those seven residues from the 047-09_1A02 F_ab_ plays a crucial role in improving the binding of the 047-09_1A02 F_ab_ to H1/Michigan/2015 (Figure 9E). The presence of these stacked variants that include deletions and substitutions in a single construct clearly demonstrates the unique ability of CODE-HB to evolve variants with a broad mutational profile comprising of substitutions, deletions and insertions; such mutational profiles typically inaccessible using most other continuous directed evolution approaches.

**Figure 9:**
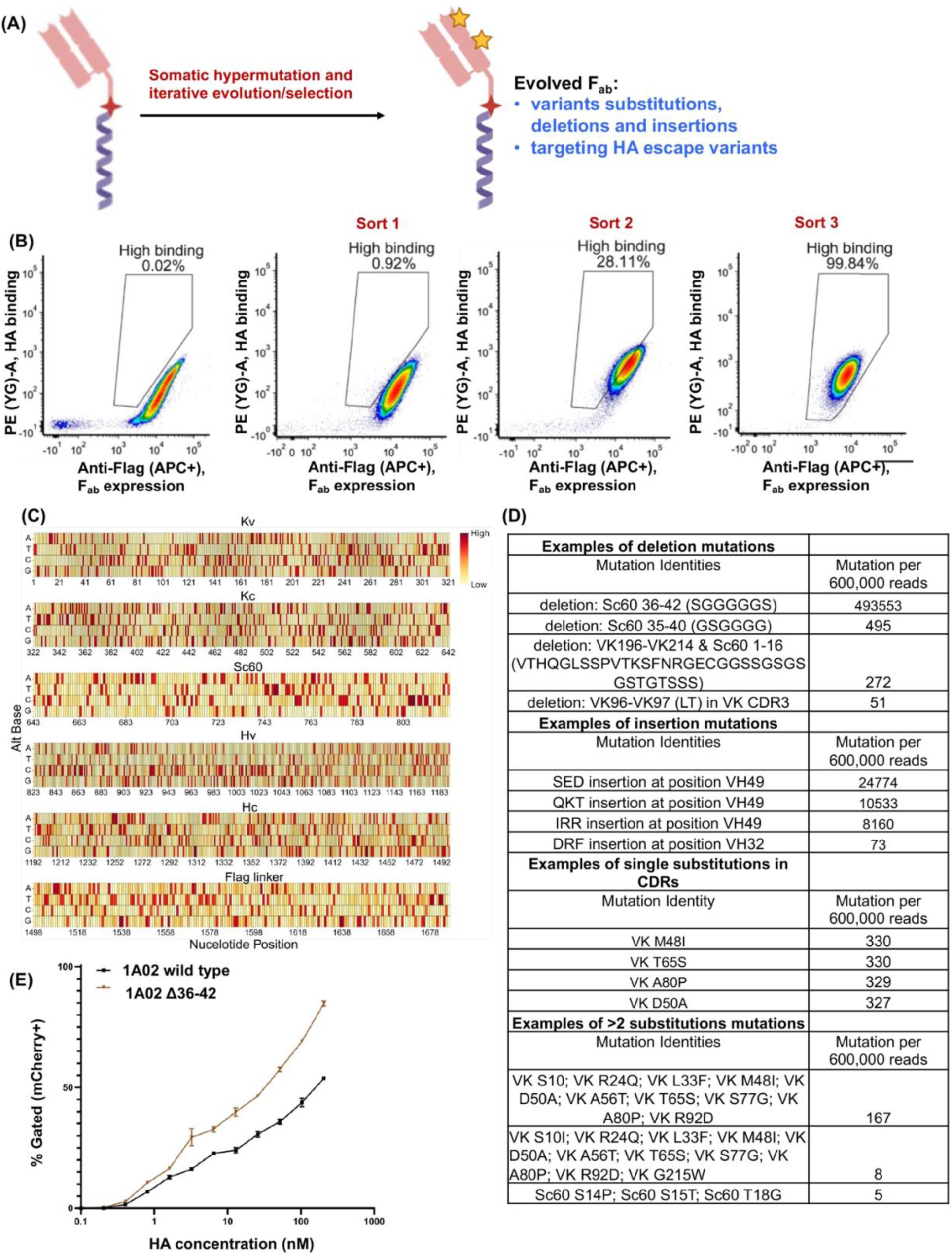
Directed evolution of the lateral-patch–binding F_ab_ 047-09_1A02 to improve binding against contemporary variants of H1N1 influenza. (**A**) Overview of 047-09_1A02 evolution using CODE-HB. (**B**) Evolutionary trajectory after iterative FACS sorting: binding to H1/Michigan/2015 plotted against surface expression of the F_ab_, illustrating an approximately linear relationship. (**C**) Heat map showing mutation density across the 047-09_1A02 F_ab_ sequence. (**D**) Recurrent in-frame deletions (span shown) and insertions (anchor nucleotide indicated) displayed alongside frequent single-nucleotide substitutions; reads with more than two amino-acid substitutions are also highlighted. Counts are normalized to 600,000 reads. (**E**) FACS-based H1/Michigan/2015 titration comparing the wild-type 047-09_1A02 F_ab_ with a variant containing a 7-amino-acid deletion (positions 36–42) in the sc60 linker between the light and heavy chains.

## Discussion

Directed evolution platforms have significantly advanced the ability to rapidly evolve biomolecules with defined functions. Advances in molecular biology and synthetic biology further resulted in the development of continuous directed evolution platforms where cell or virus-based systems are used (see introduction section). One of the key advantages of these platforms is the ability of engineered cells to introduce biomolecular diversity *in situ* without requiring *in vitro* library generation. Another attractive feature of some of these systems is the design where the evolutionary outcomes are linked to the propagation of cells or viruses with selective growth advantage. Several other efforts are ongoing to build, optimize and utilize continuous directed evolution platforms in model bacteria, yeasts, mammalian cells and viruses. Our efforts were inspired by the SHM mediated rapid antibody evolution in B cell lines. Several studies have extensively characterized the biochemistry and intrinsic mutation rates of SHM, and the timing of SHM in relation to phenotype evolution in B cells and B cell lines, including RA 1 cell lines.^21,24,64^

We focused our efforts on investigating if the SHM mechanisms in B cells can be repurposed for engineered continuous directed evolution. Despite a few previous efforts to test evolution experiments in B cells, none of these approaches have found any widespread utility. Our approach creates a roadmap to overcome several of the bottlenecks in using B cell lines for continuous directed evolution. Particularly, we developed a virus free, precise and efficient genome editing in human B cell lines. This is in direct contrast to all previous studies that use low efficiency viral transduction with random genomic integration. We further achieved stable high levels of protein expression in more than 90% of the cell population. Further, in our platform to continuously evolve variants, we developed a strategy that recruits and repurposes the inherent B cell SHM mechanisms, to rapidly and orthogonally evolve proteins of interest from a stable, safe harbor locus within human B cells. We also developed of a new B cell surface display platform for selection. This does not limit the type of antibodies or antibody fragments that can be displayed on the surface of the B cell lines. Perhaps most importantly, using our approach we achieve stable, continuous diversification and evolution of genes from a stable, safe harbor locus that expresses the desired reporter protein in more than 90% of cells. In previous B cell approaches evolve genes from a very low fraction of cells as evolution only occurs on a small fraction of cells which have a random integration event at the immunoglobulin locus. Additionally, these previous approaches are further hindered by the fact that the immunoglobulin loci can undergo loss of expression or gene deletions.

Using CODE-HB, we demonstrated that this platform can be used to rapidly evolve eGFP variants. We also developed a B cell surface display platform for displaying reporter proteins on the surface of the B cells. Using this approach, we evolved F_ab_ sequences targeting avian subtypes of influenza hemagglutinin (e.g., H5). We also demonstrated that this approach can used to rapidly and continuously evolve antibody sequences in human B cell lines. Particularly, our evolution experiment with matured antibody sequence CR9114 resulted in isolation of CR9114 variants that were able to bind H5 with higher avidity. Importantly, several of these evolved antibody variants retained their viral neutralization potential at similar or higher level to the starting CR9114. Our evolution experiments with 047-09_1A02 antibody resulted in the evolution of a F_ab_ variant that had significantly more potent binding that the starting 047-09_1A02 F_ab_ sequence. Additionally, unlike other continuous directed evolution approaches in bacteria, yeast and viruses, our approach makes it possible to attain a broad mutational spectrum comprising of substitutions, deletions and insertions, and at least in one of the evolutionary campaigns this mutational profile provides plays an important role in evolving a desired phenotype. This clearly highlights the utility of CODE-HB for continuous directed evolution.

Based on our observations, the estimated mutation rate achieved by the system was around 5 × 10⁻⁵ to 9 × 10⁻⁵ substitutions per base per generation, translating to 5 × 10⁻² to 9 × 10⁻² substitutions per kilobase per generation (0.005% to 0.009%, based on both eGFP* and F_ab_ evolution experiments, Supplementary Table 6). This rate can be contextualized by comparing it with other mutagenesis systems. Phage-Assisted Continuous Evolution (PACE), demonstrates a higher mutation rate of 7.2 × 10⁻³ substitutions per base per generation, which equates to 2.3 substitutions per kilobase (0.23%)^65^. Mutazyme, an error-prone DNA polymerase from Agilent Technologies, operates within an even higher mutation rate range of 0.3% to 1.6%. In contrast, the OrthoRep system reports a mutation rate of 1 × 10⁻⁵ per base per generation, translating to 1 × 10⁻² substitutions per kilobase per generation (0.001%).^10^ These comparisons illustrate that this system’s mutation rate is intermediate among commonly used mutagenesis platforms - it is significantly lower than that of PACE and Mutazyme, yet higher than that of OrthoRep, potentially allowing for more rapid generation of genetic diversity. We believe that it might be possible to further modulate the mutation rates of CODE-HB by incorporating various SHM recruiting sequences that have been previously identified.^33^ Using CODE-HB approach, we observe diversification of codon optimized eGFP sequences, different F_ab_ sequences, and even non-Fab sequences like unnatural linker sequences and transmembrane domains. This suggest that in principle this approach should work for diversification and possible evolution any gene of interest encoding protein or RNA. We anticipate that CODE-HB can be used for developing biologics as well as for peptides, proteins and enzymes with desired properties. This platform may allow the exploration of human cells for continuous directed evolution of medicinally relevant proteins like antibodies, antibody fragments, fluorescent proteins, metabolic enzymes, reporters and membrane proteins that often fail to function properly *in vivo* when evolved in bacterial or yeast cells,^14^ receptor/ligand pairs, transcription factors, decoy receptors amongst others. We anticipate that such synthetic systems also have the potential to be repurposed to elucidate finer molecular details of the SHM mechanisms.

### Limitations of the Study

The current mutation rates for CODE-HB are constrained by the SHM mediated mutation rates in the B cell lines. This study has not explored methods to alter these mutational rates. Similarly, this study does not explore methods to alter mutational profiles during the evolutionary campaigns. At this stage, this method requires sequential rounds of B cell passaging followed by selection and enrichment of cells with desired phenotypes. Currently, no negative selection methods are incorporated into CODE-HB, and we anticipate that incorporating negative selection could further reduce rounds of selection needed to attain a desired phenotype.

## Materials and Methods

### Bacterial strains, mammalian cell lines, growth media, DNA constructs

*E. coli DH5α* was used for all transformations and bacterial cultures and grown in either LB or 2xYT medium. Expi 293 were used for transient transfection and initial construct testing and were grown in Expi 293 Expression Medium. RA 1 (ATCC CRL-1596) were used for SHM experiments and grown in RPMI 1640 Medium supplemented with 10% FBS; both cell lines were validated by STR analysis. The oligonucleotides used in this study are listed in Supplementary Table 1, the synthesized DNA fragments are listed in Supplementary Table 2, the plasmid DNAs used in this study are listed in Supplementary Table 3 and the B cell lines generated in this study are listed in Supplementary Table 4.

### Preparation of plasmid DNA samples for transfection of RA 1 cell line

Using the QIAprep® Spin Miniprep Kit (Qaigen: 27104), plasmids for transfection were prepared from E. coli cultures. Initially, E. coli containing the desired plasmids were grown in 2 mL of 2x YT media, inoculated from glycerol stock. The next day, this culture was diluted 1% into 30 mL of LB media in a 125 mL Erlenmeyer flask and incubated at 37°C with 220 rpm shaking. After 24 hours, the culture was collected in 50 mL conical tube and centrifuged at 10,000 xg for 5 minutes, the supernatant was decanted, and the pellet was resuspended in 800 µL of P1 buffer with RNase. Following vortexing, 1 mL of P2 buffer was added and the mixture was lysed for 5-7 minutes with shaking at 150 rpm. Then, 1.4 mL of N3 buffer was added, the tube was inverted for mixing, and left with shaking for 3 minutes to form a precipitate. This was centrifuged at 12,000 xg for 5 minutes, and the clear supernatant was distributed among two spin columns for purification, following the kit’s protocol. The DNA was then serially eluted from the two columns using 45 uL of Dnase-free water, using the pass-through of the first column as the elution media for the next column.

### Transfection of RA 1 cell line

RA 1 cell line was cultured to a density of 1.5-2.0 million cells/mL using growth media RPMI 1640 + 10% FBS. For transfection, the Mirus Ingenio electroporation solution (Mirus Bio: MIR 50114) was pre-warmed by shaking at 37°C for 10 minutes. Meanwhile, 7 million cells were centrifuged at 100xg for 10 minutes at room temperature, resuspended in 100 µL Mirus electroporation buffer, and combined with 20 µL DNA solution, containing 10 µg integration vector and 10 µg Cas9/sgRNA plasmid. A final volume of 110 µL was then transferred to a 0.2 cm electroporation cuvette.

Electroporation was performed using the BioRad XCell Total Electroporation system (BioRad: 1652660) at 133V, 950 µF, infinite resistance, in a 0.2 cm cuvette, at room temperature. Post-electroporation, cells rested at room temperature for 5-7 minutes, then were resuspended in 1 mL RPMI + 10% FBS and transferred to a 6-well plate in a total volume of 3 mL RPMI + 10% FBS. Cells were then grown in this vessel for 24 hours, washed, and resuspended in 10 mL RPMI + 10% FBS media in a T025 flask, and grown for 48 hours. For determining the concentration of puromycin required for the selection of cell lines with genomic integrant, we titrated puromycin levels in the growth medium. Cell viability was evaluated using trypan blue staining and microscopic examination, as well as assessing the impact on growth rates via cell counts, all aiming to find an optimal puromycin concentration that allowed for selection without irreversibly compromising cell health or inducing irreversible morphological alterations. Using this approach, we determined that the ideal starting concentration of puromycin for effective selection of our RA 1 mutant cell lines was 0.3 µg/mL. The RA 1 cells that were transfected with p274 and p276 were recovered and selected in growth medium containing 0.3 µg/ml of puromycin for 84 hours. The volume of growth medium was increased in a stepwise manner from 10 mL to 30 mL over the course of one week.

### Selection of mutant RA 1 cell lines

Transfected RA 1 cell lines, post-cultivation in T025 flasks for 48 hours, were passaged into fresh T025 flasks at a density of 500,000 cells/mL in 10 mL volume of growth media, supplemented with 0.3 µg/mL puromycin (Sigma Aldrich: P8833). This selection medium was utilized to grow the cells for 84 hours. Subsequently, cell cultures were diluted to 15 mL by adding 5 mL of growth media and incubated for an additional 48 hours. Following this, cells were further diluted to 20 mL with 5 mL of growth media and again grown for 24 hours. The dilution process continued, increasing the volume to 30 mL with an addition of 10 mL growth media for another 24 hours. At the conclusion of the selection regimen, cells exhibiting puromycin resistance were deemed “enriched” for the integration event. These enriched cells were cryopreserved at a density of 10,000,000 cells/mL. The cryopreservation protocol involved resuspending cells in growth media amended with 5% DMSO, followed by a 30-minute chill on dry ice before storage in the vapor phase of liquid nitrogen. Our genomic DNA analysis confirms the integration of the recombinant cassette at the desired locus and suggests the generation of heterozygous mutants (Supplementary Figure 3). Typical transfection efficiencies, as detected by eGFP fluorescence 48 hours post-transfection, range from 2% to 7%. This suggests that, theoretically, less than 10% of the variants in any produced library are incorporated into the final selected population of integrant (based on eGFP integration). The low incorporation rate may certainly limit the diversity of the starting mutant library, but an average transfection would produce up to 350,000 variants as novel starting points from the wild-type sequence to subsequently diversify.

### RNA isolation and cDNA preparation

Total RNA from sorted cells was isolated using the PureLink RNA Mini Kit (ThermoFisher: 12183018A), following the manufacturer’s instructions. Cells (5,000,000) were resuspended in 600 µL Lysis Buffer with 1% BME, vortexed for cell lysis, and homogenized via a 21-gauge needle. After adding 1.5 volumes of ethanol (96-100%) and vortexing, the mixture was column-purified and eluted in 18 µL RNase-free water. The isolated RNA underwent reverse transcription with the Superscript III First Strand Synthesis Kit (ThermoFisher: 18080051), per the manufacturers’ protocol. For this, 16 µL of the isolated RNA solution was combined with the provided set of 2 µL poly-dT primer and 2 µL dNTPs, heated at 65°C for 5 minutes, then cooled on ice for 1 minute. A mix of 2 µL reverse transcriptase, 8 µL MgCl_2_, 4 µL DTT, 4 µL 10X RT buffer, and 2 µL RNase Out (total 20 µL) was prepared. Both mixes were combined, vortexed, and incubated for 50 minutes at 50°C, then denatured at 85°C for 5 minutes. Adding 2 µL RNase H and incubating for 20 minutes at 37°C finalized the cDNA mixture that was ready for use in PCR.

### PCR amplification from cDNA

PCR optimization for individual cDNA samples involved initial dilution of the cDNA to 10%, which then served as a template in PCR mixtures. PCRs were performed using PrimeStar MAX (Takara: R047A) to maximize replication fidelity. In these reactions, each primer was serially diluted 2-fold from a starting concentration of 1.6 µM. For the amplification of CR9114-based amplicons, the optimization process utilized primers SB 274A and SB 212B. Similarly, eGFP-based amplicons were obtained through optimization with primers SB 193A and SB 193B.

### Microscopy

Images were produced by the Bio-Rad ZOE Fluorescent Cell Imager.

### FACS Analysis and Sorting

To identify FLAG-tagged F_ab_ displays, APC-conjugated Anti-FLAG antibodies (Biolegend 637308) and FITC-conjugated Anti-His antibodies (ICLLab CHIS-45A-Z) were utilized, targeting the 6xHis tag on hemagglutinins H3 and H5 (SinoBiological 40859-V08H and 11700-V08H, respectively). H1-mCherry was produced by chimeric protein fusion of mCherryXL to the C-terminus of the hemagglutinin H1 ectodomain. FACS analysis and sorting were performed on a BD FACSMelody or BD Accuri C6 system, using a combination of FCS Express 6 and FlowJo11 for FACS plot generation and data analysis. Binding versus expression was quantified by establishing a linear regression between the FITC and APC signals. Cells displaying APC fluorescence two logarithms higher than untransfected controls were classified as “Expressing”. High-binding cells were isolated using triangular gating, adjusted so the gate’s hypotenuse was parallel to the regression line and elevated to include no more than 2% of parent gate events. For sorting, 6-8 million cells were processed at the maximum flow rate and collected into a collection tube with 3 mL of growth media, all maintained at 5°C.

### Third generation sequencing data analysis

Multi-FASTA files corresponding to PacBio sequencing data were analyzed using Python (3.10.12). Briefly, multi-FASTA files were imported using BioPython (1.59) and collapsed to unique reads and their occurrence. Unique reads with an occurrence threshold below 15 were discarded. Surviving unique reads were then aligned to the reference sequence using BioPython with the following alignment parameters: match_score = 2, mismatch_score = -3, open_gap_score = -5, extend_gap_score = -2, query_right_open_gap_score = 0, query_left_open_gap_score = 0, query_right_extend_gap_score = -1, query_left_extend_gap_score = -1. Alignments which introduced gaps in the interior of the alignment were discarded. Surviving unique reads were then used as a basis for mutational profile calculations. Data handling was done using Pandas (2.2.2). Histogram generation was done using matplotlib (3.9.0). Sunburst plots and pie charts were generated using Plotly (5.22.0)

### Structural Analysis and Modeling

Crystal structures of hemagglutinin H5 and CR9114 (PDB: 4FQI) were visualized using PyMOL. The predicted crystal structure of the chimeric CR9114 surface-displayed protein was calculated using AlphaFold3 and visualized using PyMOL.

### Calculation of EC50

The EC50 of surface-displayed CR9114 and its variants was determined by fitting the sigmoidal data to a dose-response curve using the Hill equation protocol in OriginPro 2024, and subsequently extracting the EC50 parameter.

### Construction of plasmid DNAs

All cloning related PCRs were performed using PrimeStar HS polymerase (Takara: R040A). The integration plasmid, p274, was sourced from Addgene (ID 164851), and the Cas9/sgRNA plasmid, p276, was also sourced from Addgene (ID 164850). Cloning on p274 was performed by extracting the proXIV sequence from RA 1 genomic DNA using primers SB 168A and SB 168B, which was then cloned directly upstream of the EF1a promoter via Gibson Assembly. Single-point mutagenesis to create an eGFP* variant was conducted using the KLD method with primers SB 171A and SB 171B. To substitute the eGFP component of the native p274 sequence with novel genes, p274 was linearized using primers SB 209A and SB 209B. F_ab_ constructs, synthesized as gBlocks by IDT, were amplified with primers SB 274A and SB 212B and cloned into p274 using Gibson Assembly. For the purpose of F_ab_ evolution, p274 was linearized with SB 209A and SB 275B, introducing an optional BsiWI cut site for restriction/ligation cloning. PCR bands were purified using a combination of the QIAprep® Spin Miniprep Kit (Qaigen: 27104) and GeneJET Gel Extraction Kit (ThermoFisher: K0691). PCR products, in 30 µL reactions, were electrophoresed on a 1% agarose gel at 130V for 14 minutes. The fluorescent bands were then excised and dissolved in 400 µL of GeneJET binding buffer, incubated at 55°C for 10 minutes, and vortexed. To this, 300 µL of 100% ethanol was added and the mixture was vortexed again before being transferred to a spin column. The purification process involved sequential washing with 500 µL of PB Buffer and 750 µL of PE Buffer from the QIAprep® kit, and then 750 µL of Wash Buffer from the GeneJET kit. After centrifugation to dry the spin column, DNA was eluted in DNase-free water.

**pSB1:** pSB2 was linearized by PCR using the oligonucleotides SB 169 A/B. The proXIV element was amplified by PCR from the genome of RA 1 using the oligonucleotides SB 168 A/B. The amplified proXIV fragment was inserted into linearized pSB2 by Gibson assembly to afford pSB1.

**pSB2:** p274 was linearized by PCR using the oligonucleotides SB 171 A/B, which encode the eGFP* knockout mutation. The linearized p274 was subject to blunt-end ligation by Kinase-Ligase-DpnI (KLD), to afford the plasmid pSB2.

**pSB3:** p274 was linearized by PCR using the oligonucleotides SB 209A / SB 275B. A gBlock of the light/heavy fusion protein F_ab_-CR9114 (PDB: 4FQI) was codon optimized for H. sapiens and amplified in three steps to incorporate the secretion signal, using the oligonucleotides SB 246A / SB 274B, followed by GL 7B / SB 274B, lastly by SB 274 A/B. A gBlock of the linker-FLAG-MHCI helix domain of the chimera was codon optimized for H. sapiens and amplified using the oligonucleotides SB 275A / SB 252B. The two fragments were inserted into the linearized p274 by Gibson assembly to afford the plasmid pSB3.

**pSB4:** pSB1 was linearized using the oligonucleotides SB 209A / SB 275B. The complete chimeric F_ab_-CR9114-Linker-FLAG-MHCI helix gene was amplified by PCR of pSB3 using oligonucleotides SB 274A / SB 252B. The gene was inserted into linearized pSB1 by Gibson assembly to afford the plasmid pSB4.

**pSZ1:** pSB1 was linearized using the oligonucleotides DJO 726/ DJO 727. The pro862 element was amplified by PCR from the genome of RA 1 using the oligonucleotides DJO 725/ DJO 728. The gene was inserted into linearized pSB1 by Gibson assembly to afford the plasmid pSZ1.

**pSZ2:** pSB1 was linearized using the oligonucleotides DJO 730/ DJO 731. The pro936 element was amplified by PCR from the genome of RA 1 using the oligonucleotides DJO 729/ DJO 732. The gene was inserted into linearized pSB1 by Gibson assembly to afford the plasmid pSZ2.

**pSZ3:** pSB1 was linearized using the oligonucleotides DJO 734/ DJO 735. The pro1417 element was amplified by PCR from the genome of RA 1 using the oligonucleotides DJO 733/ DJO 736. The gene was inserted into linearized pSB1 by Gibson assembly to afford the plasmid pSZ3.

**pDJO1:** pSB4 was linearized using the oligonucleotides DJO 124/ DJO 408. A gBlock of the light/heavy fusion protein F_ab_-047-09_1A02 was codon optimized for H. sapiens and amplified using oligonucleotides DJO 475/ DJO 476. The gene was inserted into linearized pSB4 by Gibson assembly to afford the plasmid pDJO1.

### Influenza virus propagation

Madin-Darby canine kidney (MDCK) cells (American Type Culture Collection, CCL-34) and Human embryonic kidney (HEK) 293T were grown and maintained in Minimum Essential Medium Egale supplemented with 10% fetal bovine serum (Gibco), 100 U/mL penicillin and 100 μg/mL streptomycin (Gibco), and 1× GlutaMAX (Gibco) at 37◦C, 5% CO2, and 95% humidity. To generate a seed stock, eight plasmids encoding each of the segments of influenza A/Puerto Rico/8/1934 (H1N1) virus were transfected into HEK 293T using jetOPTIMUS (Polyplus) following the manufacturer’s protocol. 48 hours post-transfection, media were collected as seed stock and TCID50 was tittered with MDCK cells. A T75 of full confluence MDCK cells were infected with MOI 0.0001 of virus and media was collected after 24 hours. The virus was aliquoted, kept in -80C and tittered for TCID50.

### The Hemagglutination Inhibition Assay

50 µl of 2-fold serial dilutions of virus in Dulbecco’s PBS (DPBS) was mixed with 50 μL 0.5% turkey RBCs (Fisher, 50-203-4867) in a round-bottom 96-well plate (Greiner Bio-One) and incubated at 4°C for 1 h. HA titer was determined to be 128 as reciprocal of highest dilution providing full hemagglutination.

Prepare 25 µL of 2-fold dilutions of antibodies in DPBS with the highest and lowest final concentrations at 100 µg/mL and 0.19 µg/ml, respectively. Mix with 25 µL of H1N1 virus with 4 HA units in round-bottom 96-well plate and incubate for 30 mins at room temperature. Then add 0.5% turkey RBCs and incubated at 4°C for 1h. All antibodies were tested in triplicate.

### Microneutralization assay

MDCK cells were seeded in 96-well plates at 8k cells per well and incubated overnight. Antibodies were serially diluted at 2-fold, starting from 100 µg/mL to 0.78 µg/mL, and then mixed with 100 TCID_50_ of influenza A H1NA virus in 100 µL of infection medium (Minimum Essential Medium supplemented with 10 mM HEPES, 0.125% BSA, 1× GlutaMAX (Gibco) and 1 µg/mL TPCK-treated trypsin). The virus-antibody mixture was incubated at 37°C, 5% CO2 for 1h. MDCK cells were washed with PBS once and infected with 100 ul virus-antibody mixture for 1h at 37°C, 5% CO_2_. Then the mixture was removed and replaced with 100 µL of infection medium. The plates were kept at 37°C, 5% CO_2_ for 24 h and fluorescence-focused assay was performed. The cells were washed with PBS, replaced with 100 µL methanol and kept at -20C for 30 mins. Then the cells were washed twice with PBS and added 3% BSA in PBS and incubate at RT for 1 h. The cells were stained with influenza A NP antibody (D67J) conjugated with FITC (Invitrogen, cat# MA1-7322) at 1µg/mL for 2 h at RT. Then the cells were washed twice with PBST and 1 time with PBS and stained with DAPI for 10 mins. The plates were then washed with PBS twice and imaged with Biotek Cytation 5 for analysis.

### qPCR experiments

Genomic DNA (gDNA) was isolated from enriched RA1-eGFP cells using a PureLink™ Genomic DNA Mini Kit, and quantified by NanoDrop. The stock gDNA (10 ng/µL) was used to generate an 8-point, 1:2 serial dilution series for the sample reactions. Primers specific for 100-200 bp regions of both the eGFP coding sequence and the chromosome 22 homology cassette were designed using the IDT PrimerQuest™ Tool. Primer pairs were synthesized at 100 µM each and combined into a final primer mixture at 0.83 μM each. Reactions were performed in 0.2 mL PCR tubes in 10 µL total volume, using PowerUp™ SYBR™ Green Master Mix. For each reaction strip (8 wells), 10 µL of primer mix was prepared and aliquoted into seven of eight tubes in a strip. 20 µL of primer mix containing 10 ng/µL gDNA was added to the first tube of a PCR tube strip and a two-fold serial dilution was performed by transferring 10 µL from the first tube into 10 µL water in the second tube, and so on through the series. Using a P20 multichannel pipette, 5 µL of each diluted gDNA/primer mix was transferred into separate PCR tubes already containing 5 µL of using PowerUp™ SYBR™ Green Master Mix, yielding a final reaction volume of 10 µL per well with 10 ng to 0.078 ng input gDNA. Amplicons corresponding to the eGFP target and the chromosome 22 homology cassette were PCR-amplified, gel-purified, and quantified. Each amplicon was diluted in an identical 1:2 series starting at 1 ng/µL and processed in parallel with sample reactions to generate standard curves. All qPCR reactions were performed on a QuantStudio 7 Pro Real-Time PCR System using the manufacturer’s recommended cycling conditions. Each run began with a 2-minute UNG incubation at 50 °C, followed by a 2-minute polymerase activation step at 95 °C. Amplification consisted of 40 cycles of denaturation at 95 °C for 15 seconds and combined annealing/extension at 60 °C for 1 minute. Quantification cycle (Cq) values were exported and plotted against log₁₀[Standard (ng/μL)] to generate standard curves. Sample copy numbers were calculated by interpolation from the linear regression of the standards, and normalized to input gDNA.

### Antibody overexpression and purification

Antibody genes encoding CR9114 and its eight variants were cloned into the pVitro vector—each construct harboring a hygromycin-resistance cassette—and transformed into *E. coli* DH5α (pV-1 through pV-8). To maximize plasmid yield without impairing growth, bacterial cultures were maintained in 37.5–75 µg/mL hygromycin, after which plasmids were purified and prepared for mammalian expression. Expi293 cells were seeded at 0.5 × 10^6^ cells/mL in 30 mL Expi293 medium and expanded to ∼6–7 × 10^6^ cells/mL, then diluted to 3 × 10^6^ cells/mL 24 hours before transfection. For each milliliter of cells, 4 µL ExpiFectamine™ 293 reagent was mixed with 46 µL Opti-MEM and incubated for 5 minutes, while separately 1 µg of plasmid DNA was diluted in 50 µL Opti-MEM; the two mixtures were then combined and incubated for an additional 20 minutes. During this interval, cells were counted and adjusted to 2–3 × 10^6^ cells/mL in 125 mL flasks, then returned to the incubator. The transfection mixtures were then added to the cultures, which were incubated for 18 hours in the incubator set to 37 °C with 8% CO₂. Finally, Enhancer 1 (6 µL/mL) and Enhancer 2 (60 µL/mL) were added sequentially, and cells were cultured for a further 3–5 days to allow robust antibody expression. Antibodies were purified from the clarified culture supernatant by Protein A affinity chromatography (MabSelect, Cytiva). First, expression medium was harvested from 125 mL flasks and clarified by two successive spins at 4,200 × g for 10 minutes, with the supernatant transferred to fresh tubes after each centrifugation. The clarified supernatant was then filtered through a 40 µm mesh into a chilled tube. Meanwhile, 750–1,000 µL of mAbselect resin beads were pelleted at 1,500 × g for 5 minutes to remove the 20% ethanol storage buffer, then washed twice with 12 mL binding buffer (20 mM sodium phosphate, 150 mM NaCl, pH 7). The washed resin was resuspended in 1–3 mL binding buffer and combined with the chilled supernatant, then gently rocked at 4 °C for 2 hours to bind antibody. The resin–antibody slurry was applied to a gravity-flow column, washed with 20 mL of binding buffer (delivered as two 10 mL aliquots), and eluted with 5 mL of elution buffer (50 mM sodium phosphate, pH 3.0). Finally, the pooled eluate was concentrated using a 100 kDa MWCO, 4 mL Amicon concentrator and the antibody concentration was determined by Qubit fluorometry.

### Antibody binding assays using ELISA

ELISA assays were carried out in Nunc Maxisorp 96-well plates. Lyophilized H5 hemagglutinin (HA) was reconstituted to 0.2 mg/mL according to the manufacturer’s instructions (Sino Biological, A/Vietnam/1194/2004), then diluted 1:200 in phosphate-buffered saline (PBS). One hundred microliters of the diluted HA solution were added to each well and the plate was incubated overnight at 4 °C. The following day, wells were emptied by pipetting, washed once with 410 µL PBS, and blocked with 410 µL of 1% bovine serum albumin (BSA) in PBS for 1 hour at room temperature. After blocking, the solution was removed and wells were washed once with 410 µL PBS. Purified monoclonal antibodies (mAbs) were prepared in a separate dilution plate by first bringing each to 400 nM in 150 µL of 1% BSA/PBS, then performing two-fold serial dilutions across the plate, leaving one well blank as a no-antibody negative control. Seventy microliters of each mAb dilution were transferred to the antigen-coated plate and incubated for 2 hours at 37 °C. Wells were then emptied and washed four times with 410 µL PBS before addition of goat anti-human IgG1-HRP secondary antibody at a 1:3,000 dilution in 1% BSA/PBS. After a 2-hour incubation at 37 °C, wells were washed five times with 410 µL PBS. One hundred microliters of TMB substrate (Invitrogen) were added to each well and allowed to develop for 30 minutes at room temperature, then the reaction was stopped with 50 µL of 2 M H₂SO₄. Absorbance at 450 nm was measured using a Cytation5 Plate Reader.

### Sequencing analysis for 047-09_1A02 evolution experiments

PacBio sequencing data for the 047-09_1A02 lineage were processed in Python (v3.10) leveraging Biopython (v1.81), pandas, and matplotlib. Initially, raw reads were oriented by pairwise alignment against annotated reference sequences using Biopython’s PairwiseAligner, then subjected to high-throughput mapping with Minimap2 (v2.26). Resulting SAM files were converted to sorted, indexed BAMs via SAMtools (v1.17). For mutation profiling, unique read sequences and their abundances were imported from pre-parsed CSV tables. Reads failing to meet an occurrence threshold (≥15) or deviating by more than ±10% from the expected amplicon length were discarded. Remaining high-confidence reads were globally aligned to the reference, and base-level variants—including substitutions, insertions, and deletions—were catalogued. Coding-region nucleotide changes were translated to amino acid substitutions; synonymous mutations were excluded, and nonsynonymous events were tabulated to generate position-wise mutation frequencies and comprehensive amino acid–level mutation landscapes. To interrogate mutational linkage, nonsynonymous substitutions within each read were used to construct pairwise co-occurrence matrices. The most prevalent co-occurring mutation pairs were visualized as heatmaps, and hierarchical clustering of the co-occurrence data produced Newick-format trees delineating mutational lineages. Indel events were extracted from the aligned BAM files using pysam, with each insertion or deletion annotated by genomic position, size, and local sequence context. Aggregate indel counts and length distributions were displayed in bar-plot form to summarize insertion and deletion patterns across the dataset.

### Genomic DNA sequencing analysis

Genomic DNA was harvested from RA1 EGFP* cells at early (passage 3) and late (passage 10) time points using the PureLink Genomic DNA Kit, following the manufacturer’s protocol. On-target editing at the EGFP* locus was assayed by PCR amplification with primers SB1/SB2, yielding a ∼700 bp fragment; the ARID1A locus on chromosome 1 served as an off-target control and was amplified in parallel. Amplicons were purified and subjected to paired-end Illumina MiSeq sequencing (2×250 bp reads). Raw reads were mapped to either the EGFP* or ARID1A reference sequences via Minimap2 (v2.26) using short-read alignment presets. Resulting SAM files were converted into sorted, duplicate-marked BAMs with SAMtools (v1.17). Only properly paired, uniquely mapped reads were retained for variant calling. Substitutions, insertions, and deletions were extracted using custom Python routines built on the pysam and Biopython libraries; high-confidence single-nucleotide variants were filtered by base quality (Q ≥ 30). Mutation counts were tallied by nucleotide position and aggregated into 10-nt bins to reveal hotspot regions. Per-base coverage depth was likewise computed and binned to evaluate uniformity and to flag any under-sequenced intervals within each amplicon.

For naïve pool mutational analysis, amplicon library preparation and paired-end MiSeq sequencing followed the protocols established for on- and off-target mutation profiling, with sequencing depth increased to 5 million reads per sample to maximize coverage and sensitivity. MiSeq reads were quality-filtered (Phred Q≥30) and aligned to the EGFP (on-target) or ARID1A (off-target) reference. Alignments were coordinate-sorted and only mapped reads with usable sequence were retained, correcting reverse-strand reads by reverse complementing based on the SAM flag. Identical sequences were then collapsed to a single entry and assigned a read support count (number of raw reads supporting that unique sequence). Within predefined analysis windows (e.g., EGFP nt 70–655; ARID1A nt 250–860), single-nucleotide substitutions were called by globally aligning each unique sequence to the reference using Biopython’s PairwiseAligner (match 2.0, mismatch −1.0, gap-open −5.0, gap-extend −0.5). Mutation frequency was summarized weighted by read support and very low-support events were excluded (occurrences_weighted < 2000) to account for PCR and sequencing error rate. In order to compare between naïve pool libraries, we normalized total signal to 5×10⁶ effective reads. All analyses were performed in Python (pandas, numpy, matplotlib, pysam, Biopython). After quality filtering, filtering occurrences less than 2000, followed by comparing to reference sequences, we observed no new mutations in ARID1A sequences when sequences from passage 0 (prior to evolution) cells were compared to passage 8 cells (Supplementary Figure 11). In contrast, we observed a whole series of mutations for our evolution experiments for on-target genes of interest.

## Acknowledgments

The research reported in this publication was supported by the HHMI EPI grant (UI Award #111112). This funding was received by A.P.M. The authors thank Alvaro Hernandez, Chris Wright and Christopher Fields (all at Roy J. Carver Biotechnology Center, UIUC) for helpful discussions on sequencing studies. A.P.M. thanks Prof. Wilfred van der Donk, Prof. Nicholas Wu and Prof. Beth Stadtmueller for helpful discussions on biochemical experiments. A.P.M. thanks Prof. Wilfred van der Donk for reading the manuscript and providing critical feedback.

## Author contributions

A.P.M. and S.B. conceived the project. A.P.M., S.B., D.J.O., G.L., S.Z., and H.X. designed experiments. S.B. S.B., D.J.O., G.L., S.Z., and H.X. performed experiments and obtained all data reported in this manuscript. Sequencing was performed at the Roy J. Carver Biotechnology Center at University of Illinois Urbana-Champaign. S.B., D.J.O., J.Q. analyzed the sequencing data. S.L. and J.J.G. provided recombinant hemagglutinin. A.P.M., S.B., D.J.O., G.L., S.Z., J.Q., and H.X. analyzed the data. All authors helped with manuscript preparation.

## Data Availability

All the data except the raw sequencing files is listed in the manuscript and the supporting information. The raw sequencing datasets generated and analyzed during this study are available from the corresponding author upon reasonable request.

