## Supporting Information for "Virus-free continuous directed evolution in human cells using somatic hypermutation"

[#]Contributed equally to this work

[$]Contributed equally to this work

### Supporting Information

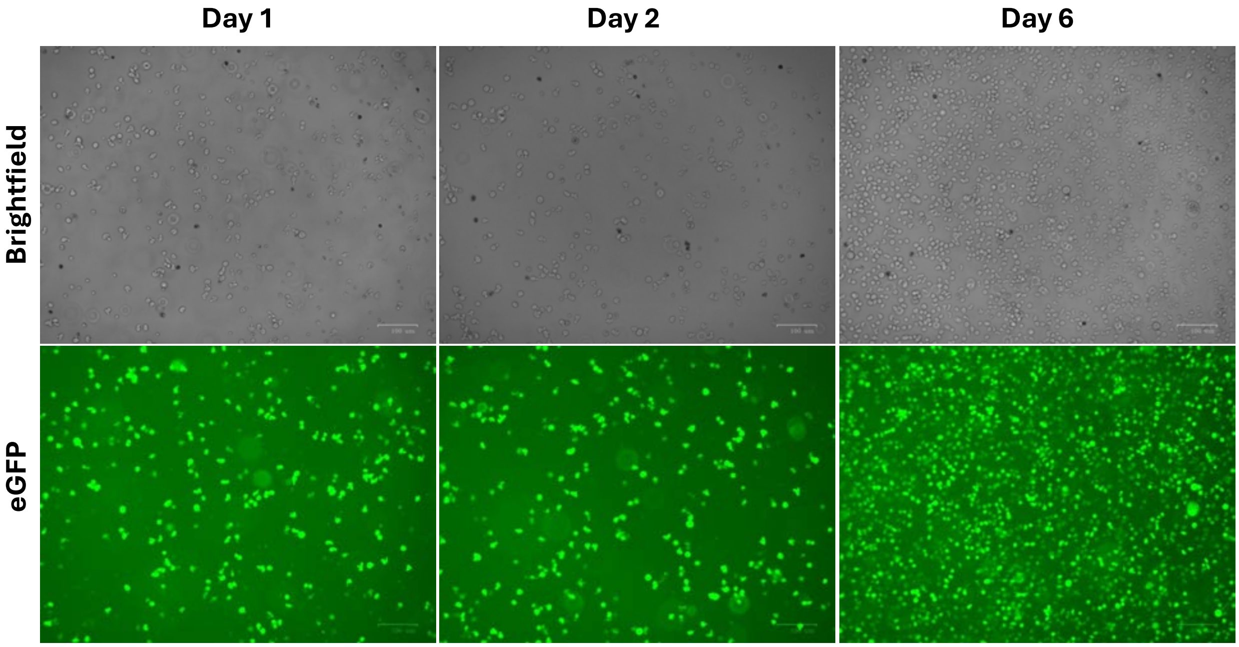


**Supplementary Figure 1: Microscopy images:** Time course of selection for puromycin-resistant mutant cell lines expressing eGFP.

**A**

| IgHV Gene | Locus | Percent Identity | Alignment Length |
|---|---|---|---|
| V4-4 | NC_000014.9: 106012569 - 106012357 | 91.549 | 213 |
| V4-20 | NC_000014.9: 106324900 - 106324689 | 89.671 | 213 |
| V4-30-2 | NC_000014.9: 106349932 - 106349721 | 91.08 | 213 |
| V4-34 | NC_000014.9: 106374306 - 106374094 | 100 | 213 |
| V4-55 | NC_000014.9: 106606764 - 106606552 | 89.671 | 213 |
| V4-59 | NC_000014.9: 106627893 - 106627681 | 91.08 | 213 |
| V4-61 | NC_000014.9: 106639769 - 106639557 | 90.141 | 213 |
| V4-80 | NC_000014.9: 106873400 - 106873187 | 88.785 | 214 |

**B**

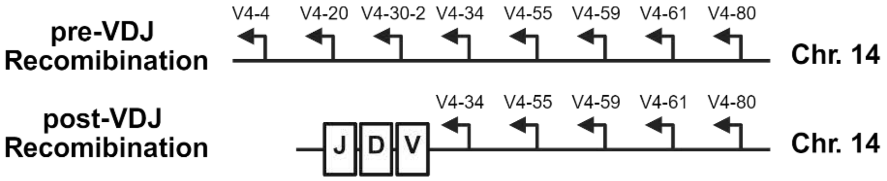


**C**

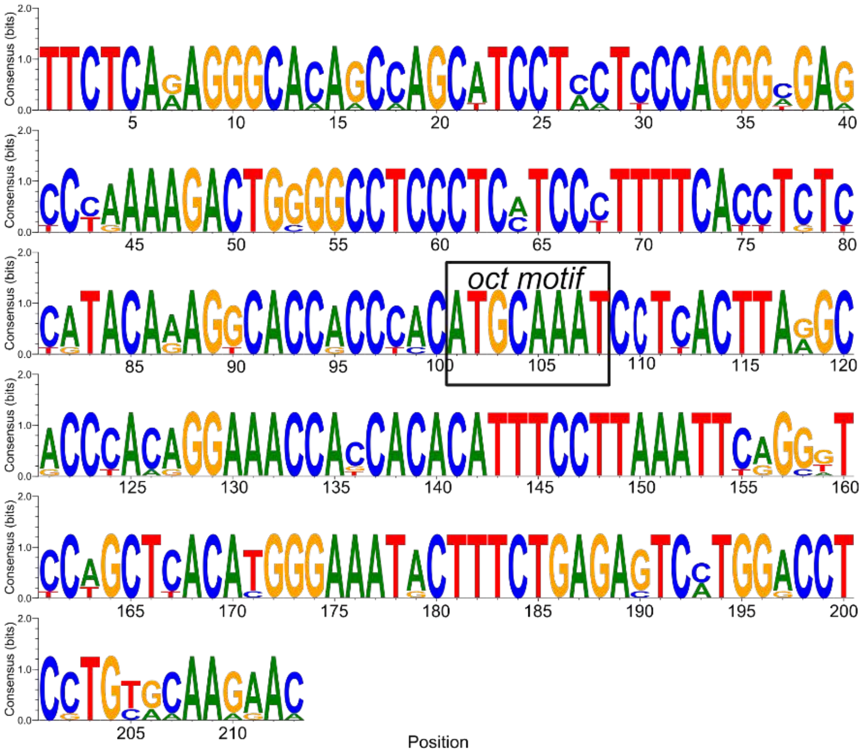

**D**

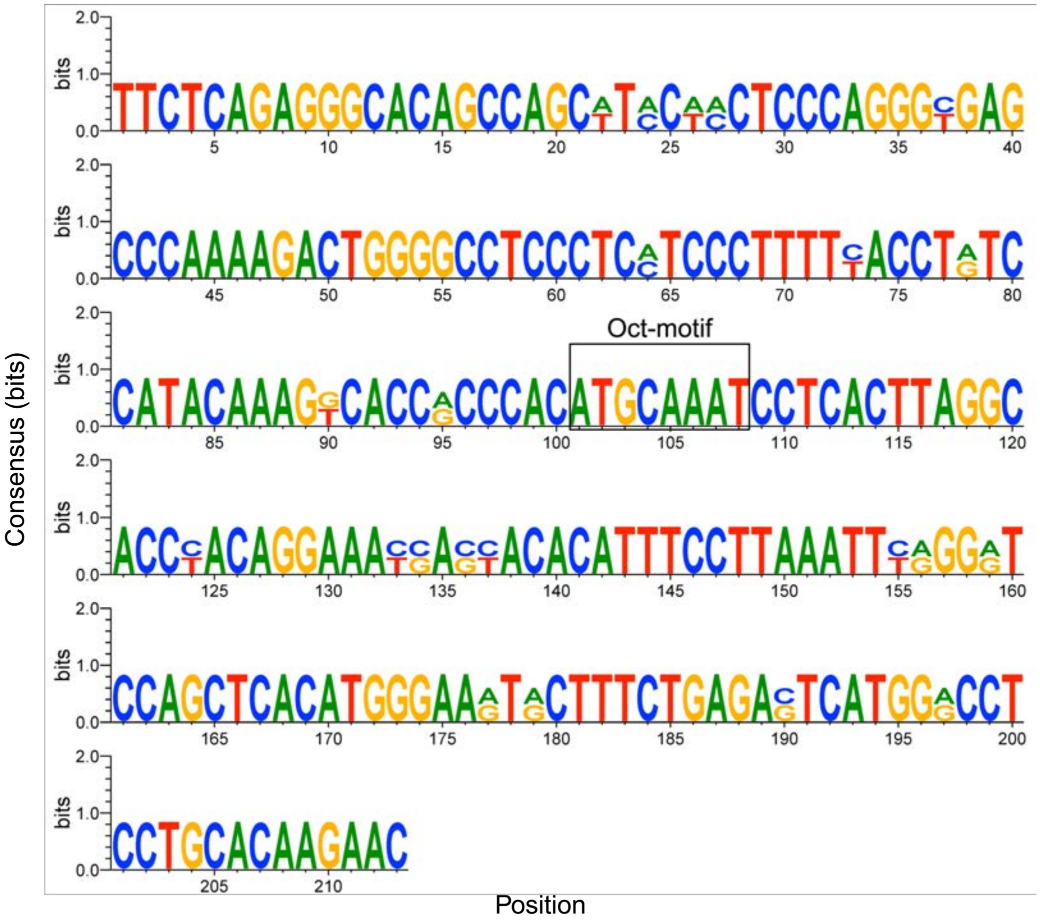


**Supplementary Figure 2: Alignment of various proXIV sequences from the V4 family. (A**) Table of aligned sequence to proXIV(4-34). (**B**) Schematic layout of the IGHV4 family dispersed among the IgH locus, pre- and post- recombination. (**C**) Sequence logo analysis of the eight proXIV-like sequences obtained from genomic alignment (boxed: *oct*-binding motif). (**D**) Sequence logo analysis of proXIV-1 and proXIV-2 sequences obtained from genomic alignment. A 213 base pair region in the vicinity of Oct-motif was chosen for alignment.

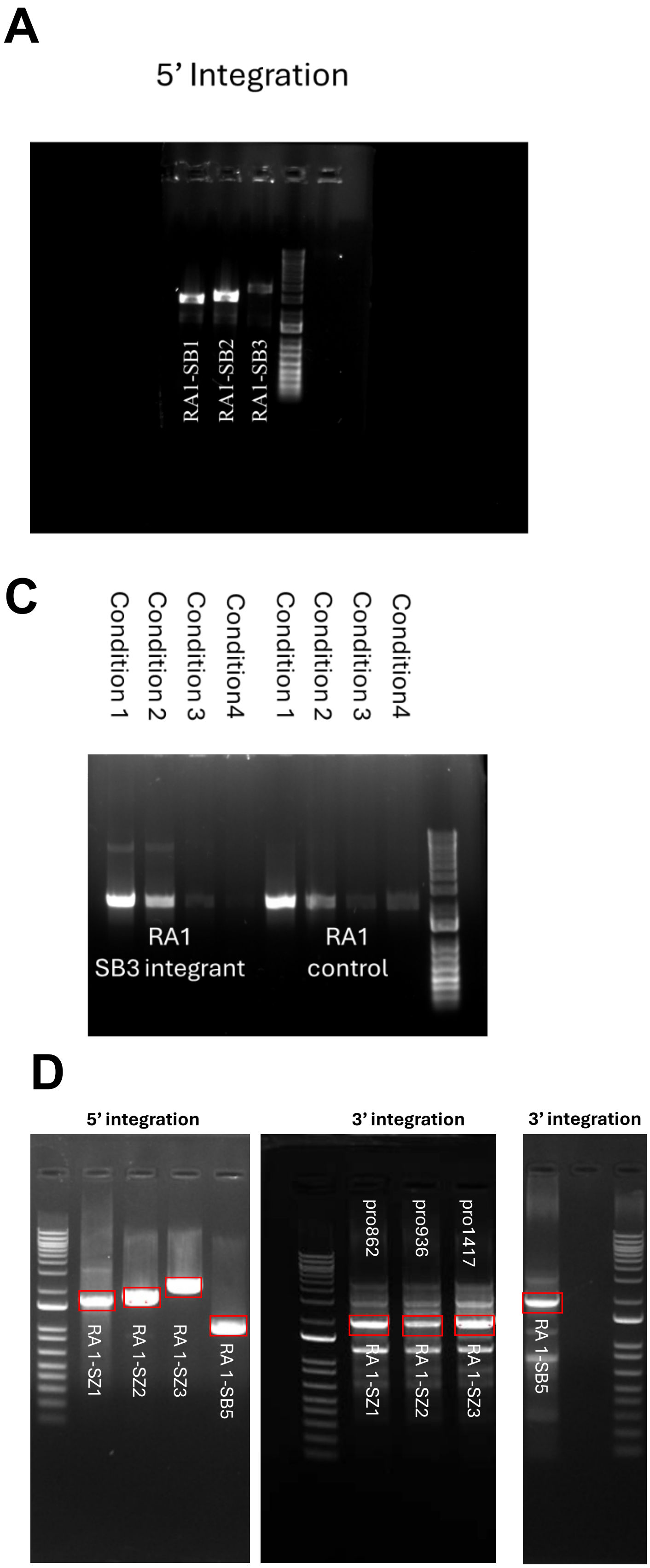
A
5’ Integration
RA1-SB1
RA1-SB2
RA1-SB3
B
3’ Integration
RA1-SB1
RA1-SB2
RA1-SB3
C
Condition 1
Condition 2
Condition 3
Condition4
Condition 1
Condition 2
Condition 3
Condition4
RA1
SB3 integrant
RA1
control
D
5’ integration
RA 1-SZ1
RA 1-SZ2
RA 1-SZ3
RA 1-SB5
3’ integration
pro862
pro936
pro1417
RA 1-SZ1
RA 1-SZ2
RA 1-SZ3
3’ integration
RA 1-SB5

## E

| | puro | eGFP | 3' Homology |
|---|---|---|---|
| **qPCR Concentration (ng/uL)** | **14.83** | **17.08** | **17.23** |
| **Relative Zygosity** | **0.86** | **0.99** | **1.00** |

**Supplementary Figure 3: Genomic DNA analysis of the mutant cell lines**. (**A**) PCR analysis to confirm the integration locus at the 5'-end of integration site (expected size ~2650 bp for RA1-SB1, 2860 bp for RA1-SB2, and ~3900 bp for RA1-SB4). (**B**) PCR analysis to confirm the integration locus at the 3' end of the integration site (expected size ~3550 bp for RA1-SB1 and RA1-SB2, and ~4500 bp for RA1-SB3). (**C**) PCR to confirm heterozygosity of the RA1-SB3 population. (**D**) Genomic DNA PCR analysis to confirm cell lines. (**E**) qPCR results to determine the zygosity of eGFP integration into the genomic DNA.

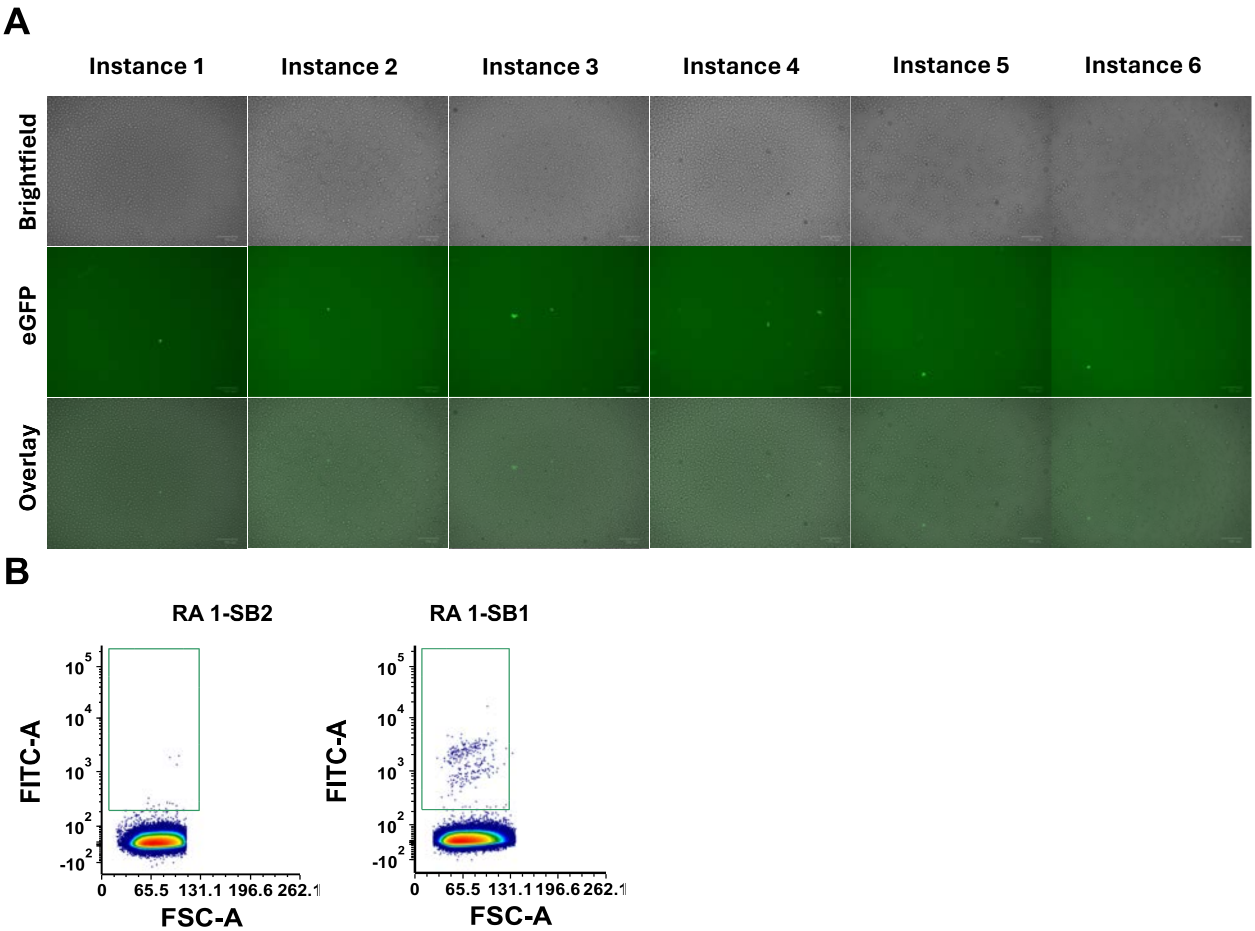


**Supplementary Figure 4: Microscopy images for the eGFP* evolution experiments**. (**A**) Instances of eGFP* reversion. (**B**) Flow cytometry density plots showing the appearance of significant levels of eGFP positive signals in RA 1-SB1 cell lines after multiple rounds of evolution and lack of eGFP positive signals in the control cell lines, RA 1-SB that express eGFP* but lack proXIV-1 sequence.

**(A)**

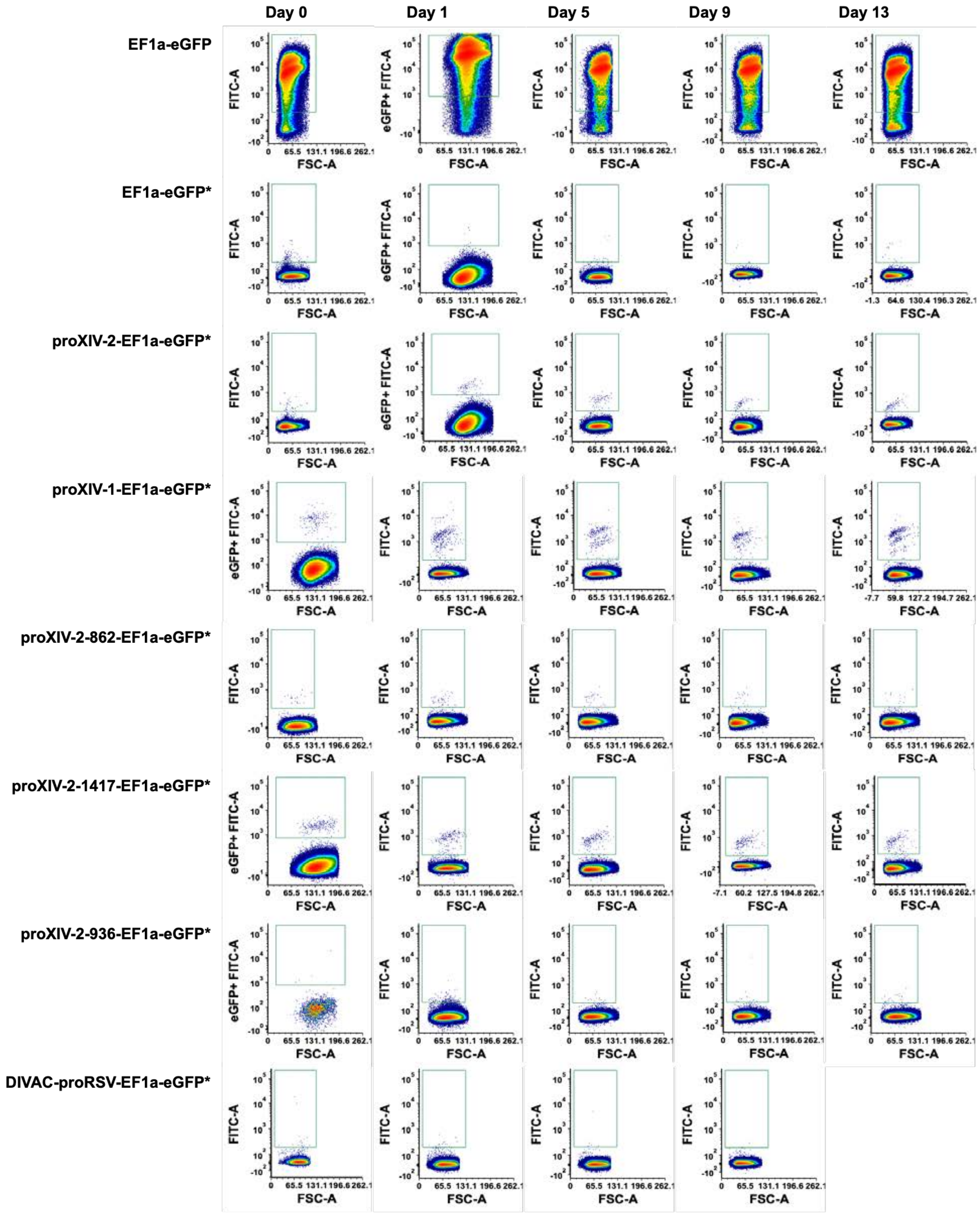

Day 0
Day 1
Day 5
Day 9
Day 13
EF1a-eGFP
EF1a-eGFP*
proXIV-2-EF1a-eGFP*
proXIV-1-EF1a-eGFP*
proXIV-2-862-EF1a-eGFP*
proXIV-2-1417-EF1a-eGFP*
proXIV-2-936-EF1a-eGFP*
DIVAC-proRSV-EF1a-eGFP*
FITC-A
eGFP+ FITC-A
FSC-A

(B)

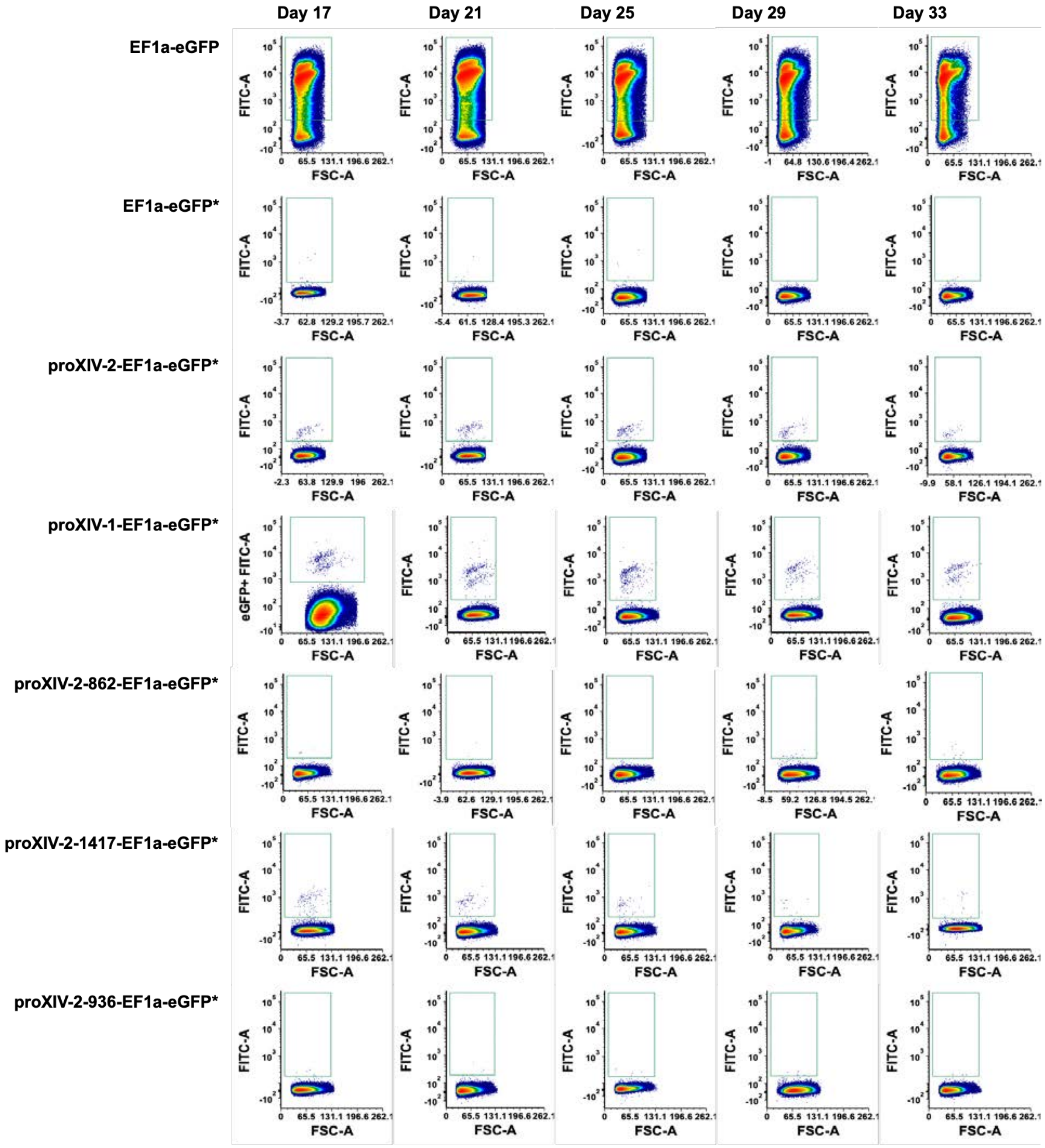


**Supplementary Figure 5: Flow cytometry density plots for mechanistic characterization of SHM recruiting sequences.** FACS density plots of evolved cells that have either proXIV-1-EF1a, proXIV-2-EF1a, proXIV-2-862-EF1a, proXIV-2-936-EF1a, DIVAC-EF1a upstream of eGFP*. In addition to this there are FACS density plots corresponding to positive control (eGFP expression) and a negative control, i.e., cells expressing eGFP* but lacking any of the SHM recruiting sequences. This figure shows evolution experiments from day 17 to day 33 after generation of cell lines. Note that in addition to proXIV sequences, we also tested previously identified Diversification Activator sequence (DIVAC) sequence. However, in our hands, addition of this sequence resulted in loss of cell viability after four passages.

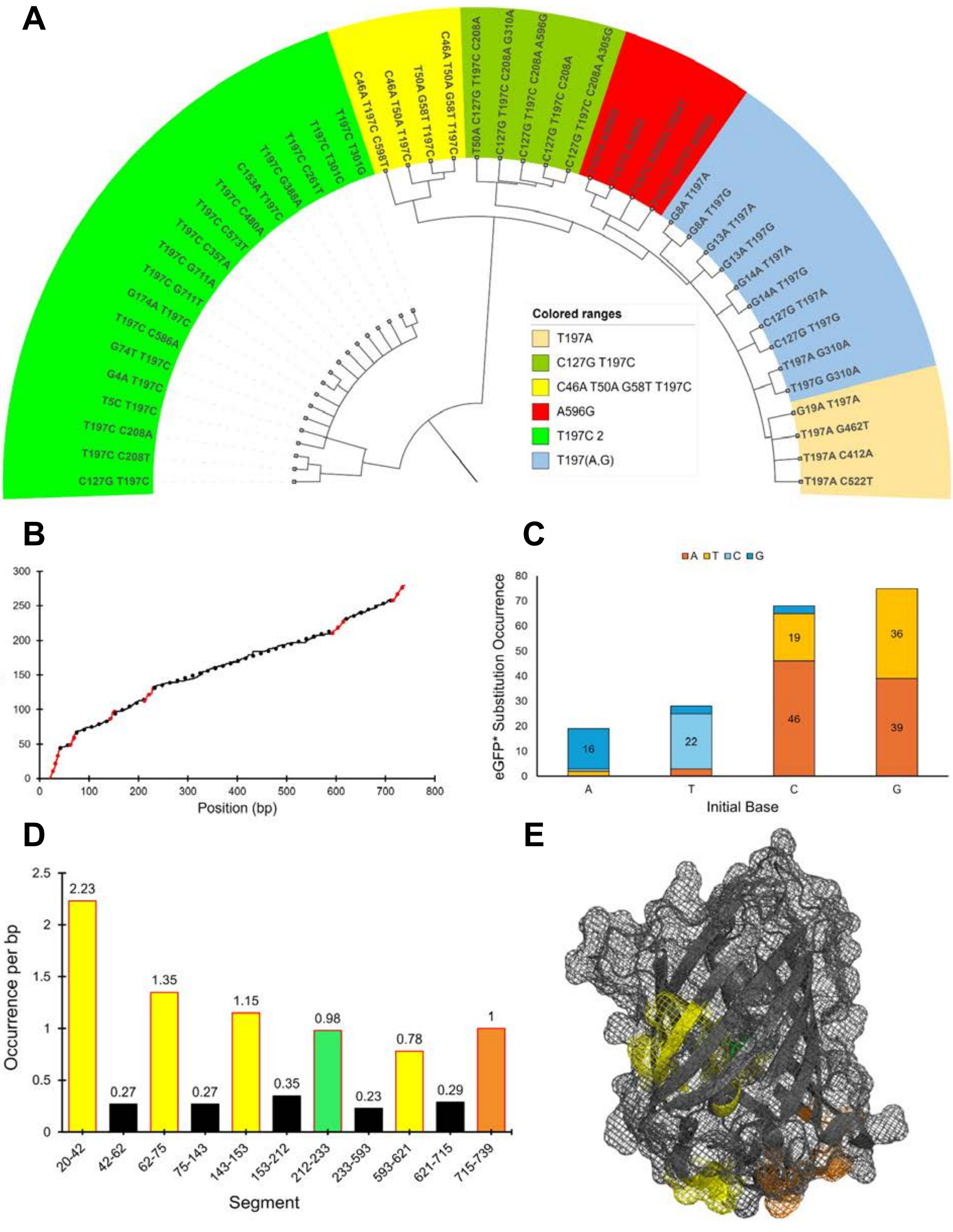
A
C46A T197C C598T
C46A T50A T197C
T50A G58T T197C
C46A T50A G58T T197C
T50A C127G T197C C208A
C127G T197C C208A G310A
C127G T197C C208A A596G
C127G T197C C208A
C127G T197C C208A A305G
T197C T301G
T197C T301C
T197C C261T
T197C G388A
C153A T197C
T197C C480A
T197C C573T
T197C C357A
T197C G711A
T197C G711T
G174A T197C
T197C C586A
G74T T197C
G4A T197C
T5C T197C
T197C C208A
T197C C208T
C127G T197C
G8A T197A
G8A T197G
G13A T197A
G13A T197G
G14A T197A
G14A T197G
C127G T197A
C127G T197G
T197A G310A
T197G G310A
G19A T197A
T197A G462T
T197A C412A
T197A C522T
Colored ranges
T197A
C127G T197C
C46A T50A G58T T197C
A596G
T197C 2
T197(A,G)
B
Sum of Unique Instances
Position (bp)
C
A T C G
eGFP* Substitution Occurrence
Initial Base
16
22
46
19
39
36
D
Occurrence per bp
Segment
2.23
0.27
1.35
0.27
1.15
0.35
0.98
0.23
0.78
0.29
1
20-42
42-62
62-75
75-143
143-153
153-212
212-233
233-593
593-621
621-715
715-739
E

**Supplementary Figure 6: The mutational landscape of the eGFP* sequences from the GFP positive sorted cells.** (**A**) A representation to indicate the occurance of single point mutations, and multiple stacked mutations in single eGFP* transcript. (**B**) Integral plot of occurrences over the sequence length of eGFP*. (**C**) Bar chart depicting the mutational bias within eGFP*. (**D**) Bar chart of slopes retrieved from each segment of the integral plot, representing mutational hotspots within eGFP*. (**E**) Crystal structure of eGFP (PDB: 2Y0G), overlayed with color codes corresponding to the hotspot bar chart.

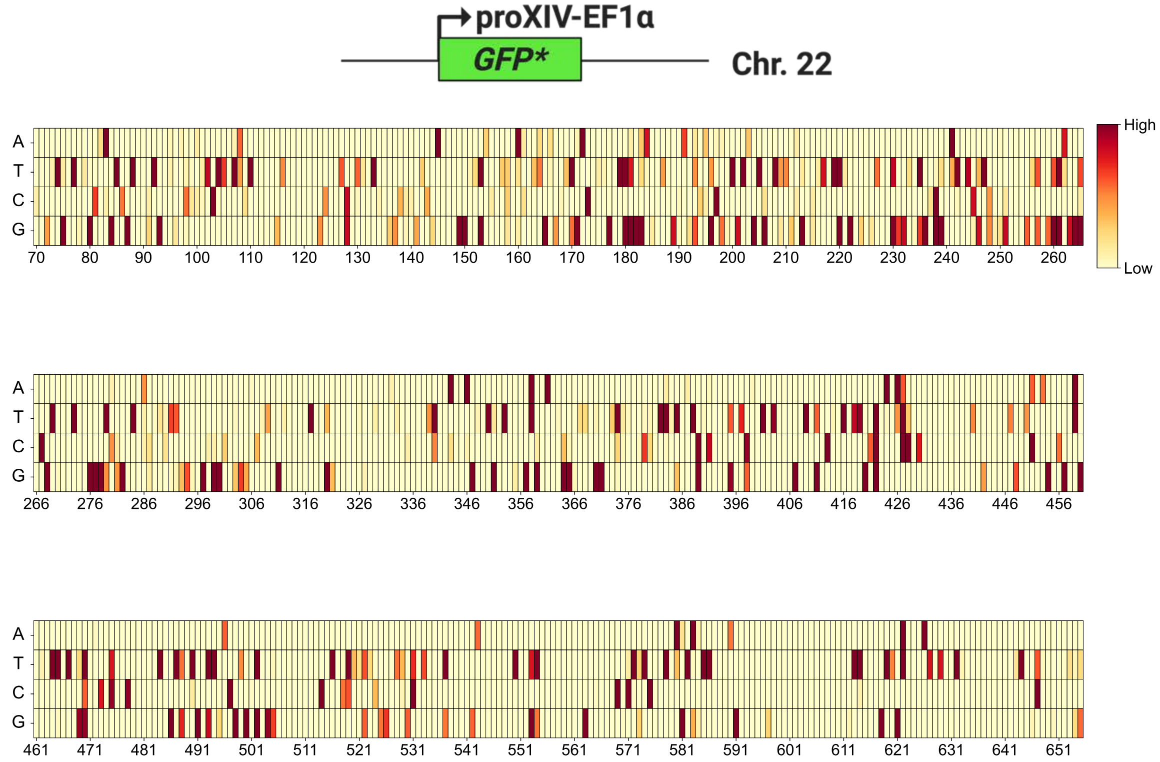


**Supplementary Figure 7: Additional results from sequencing of eGFP* genomic locus from unsorted samples at passage 3**. Heatmap showing density of mutations across the eGFP* sequence at passage 3.

| Examples of deletion mutations | |
|---|---|
| **Mutation Identities** | **Mutation per 5000000 reads** |
| deletion 94-99 (GGCGAG) | 703 |
| deletion 589-600 (CCCGACAACCAC) | 2695 |
| deletion 259-264 (TCCGCC) | 998 |

| Examples of insertion mutations | |
|---|---|
| **Mutation Identities** | **Mutation per 5000000 reads** |
| TAA insertion at position 285 | 3810 |
| TAC insertion at position 285 | 1072 |

| Examples of substitution mutations | |
|---|---|
| **Mutation Identity** | **Mutation per 5000000 reads** |
| T197C | 42335 |
| T254G | 41943 |
| A460G | 37258 |
| G350C | 8825 |
| C191G | 7689 |

**Supplementary figure 8: Examples of nucleotide level mutations across eGFP* locus.** Recurrent in-frame deletions (span shown) and insertions (anchor nucleotide indicated) are displayed alongside frequent single-nucleotide substitutions. Counts are normalized to 12 million.

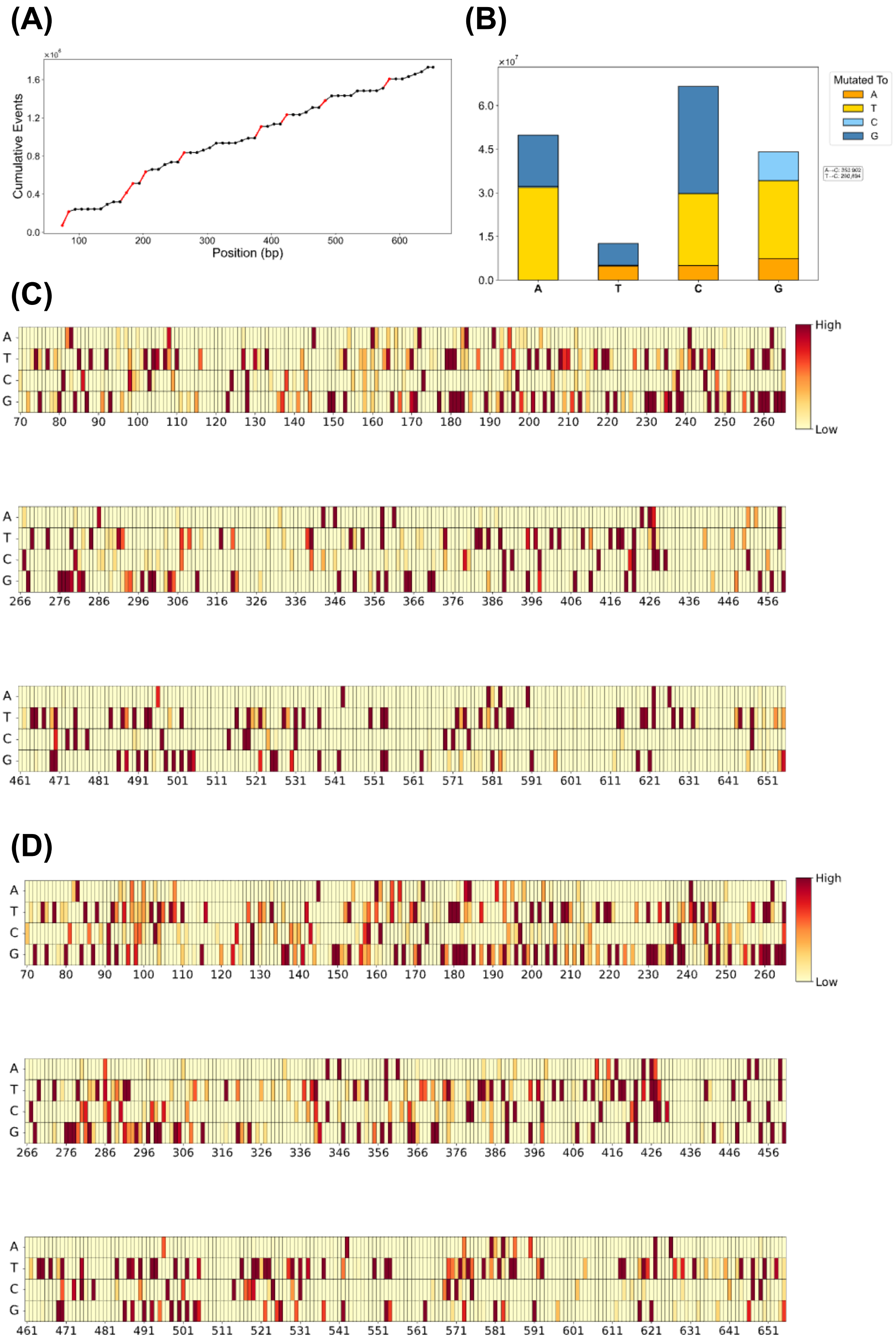
(A)
Cumulative Events
Position (bp)
(B)
Mutated To
A
T
C
G
(C)
High
Low
(D)
High
Low

**Supplementary Figure 9: eGFP* evolution experiments were performed in replicates, and these are the results from sequencing of cDNA corresponding to eGFP* transcripts in a naïve pool of unsorted cells. (A)** Integral plot of occurrences over the sequence length. Red highlight indicates mutational hotspot. **(B)** Bar plot comparison of nucleotide mutations in the eGFP*. Mutations are plotted as the percentage of total reads/sum of all mutations. **(C)** Heatmap showing density of mutations across the eGFP* sequence at passage 3. (**D**) Heatmap showing density of mutations across the eGFP* sequence at passage 8.

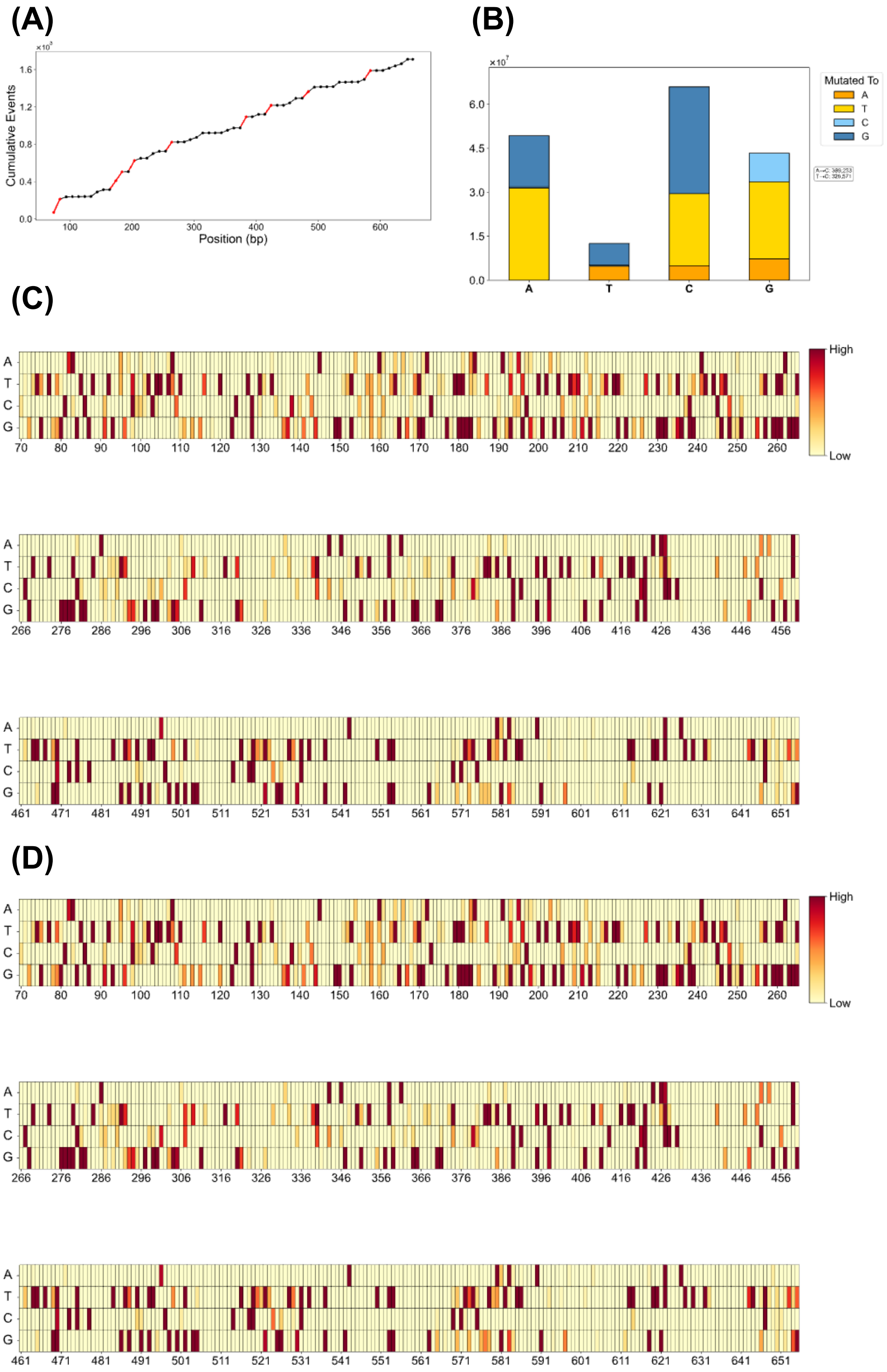
(A)
(B)
(C)
(D)
Cumulative Events
Position (bp)
Mutated To
A
T
C
G
High
Low

**Supplementary Figure 10: eGFP* evolution experiments were performed in replicates, and these are the results from sequencing of genomic locus corresponding to eGFP* in a naïve pool of unsorted cells. (A)** Integral plot of occurrences over the sequence length. Red highlight indicates mutational hotspot. **(B)** Bar plot comparison of nucleotide mutations in the eGFP*. Mutations are plotted as the percentage of total reads/sum of all mutations. **(C)** Heatmap showing density of mutations across the eGFP* sequence at passage 3. (**D**) Heatmap showing density of mutations across the eGFP* sequence at passage 8.

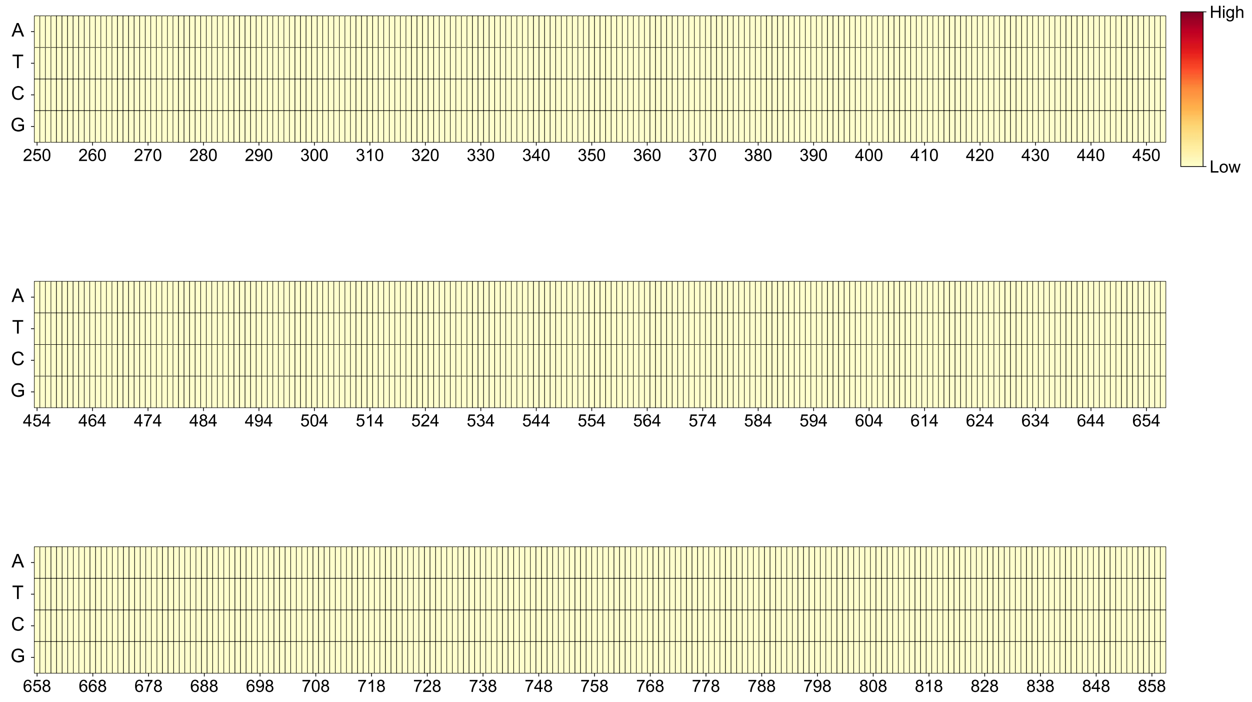


**Supplementary Figure 11: Sequencing experiments to determine off-target mutations on the genome.** To understand the background genomic mutational rate, we performed sequencing analysis ARID1A locus of a representative population of cells was sequenced before and after evolution experiments. The heat map demonstrates the lack of detection of any high abundance and high confidence exclusive mutations. Under the same analysis criterion, we observe a wide range of mutations for our on-target reporter genes (e.g., eGFP*, antibodies) after evolution experiments

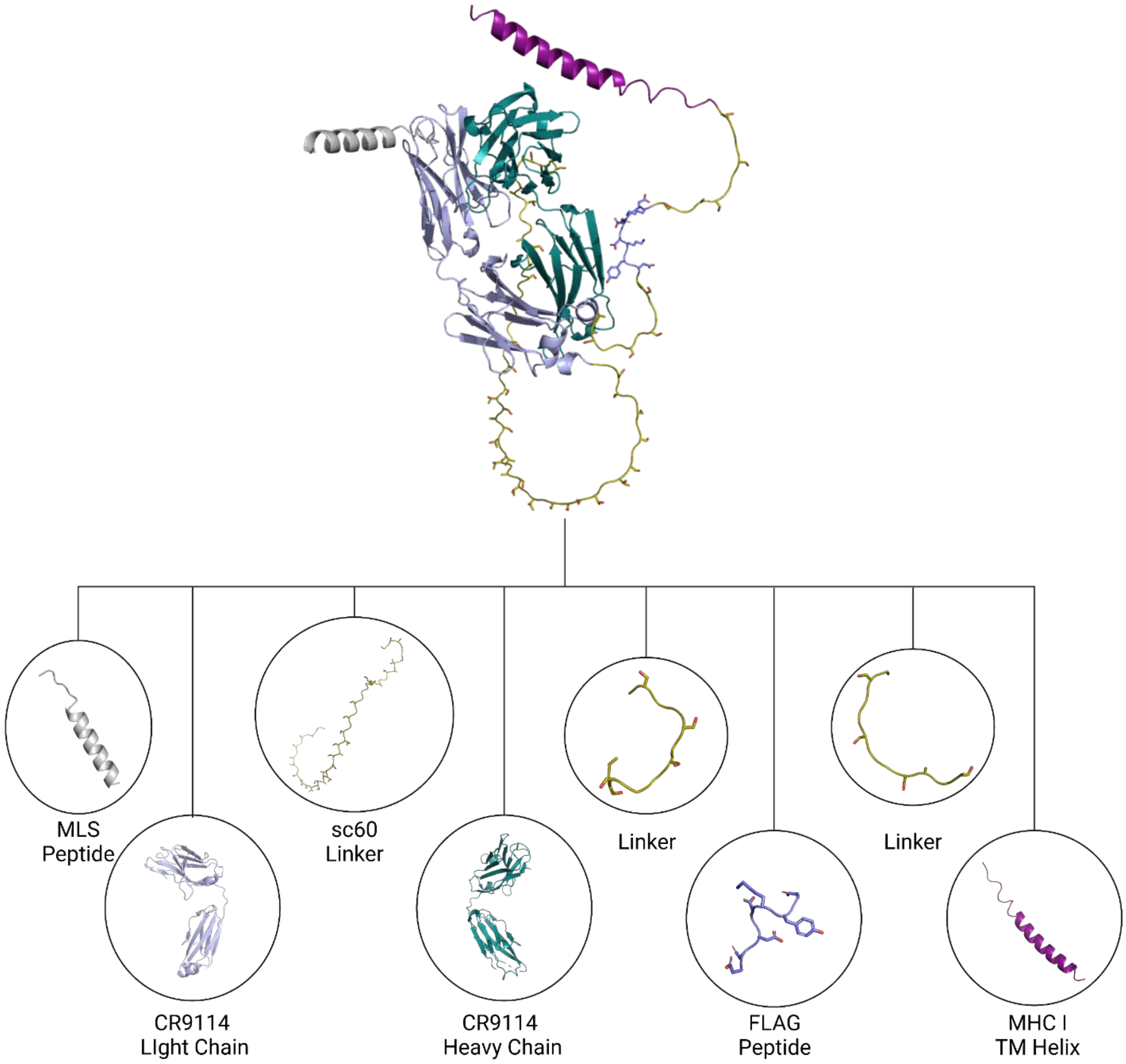


**Supplementary Figure 12:** AlphaFold3 predicted structure of the fusion protein and visualized representations of the various components.

A

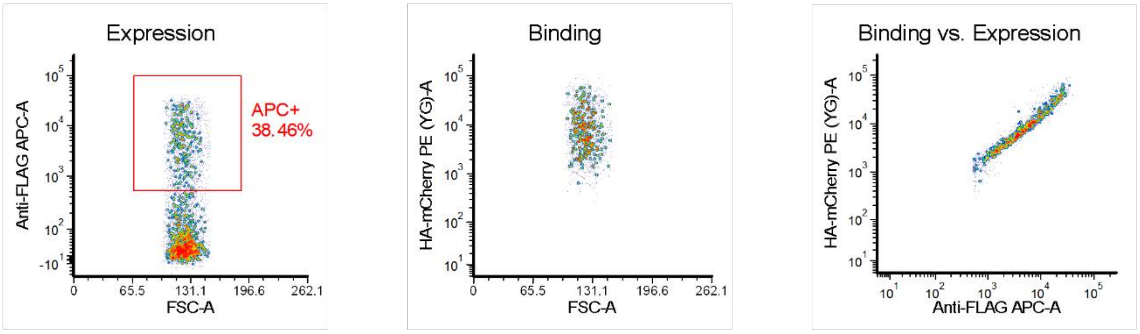
Expression
Anti-FLAG APC-A
FSC-A
APC+
38.46%
Binding
HA-mCherry PE (YG)-A
FSC-A
Binding vs. Expression
HA-mCherry PE (YG)-A
Anti-FLAG APC-A

B

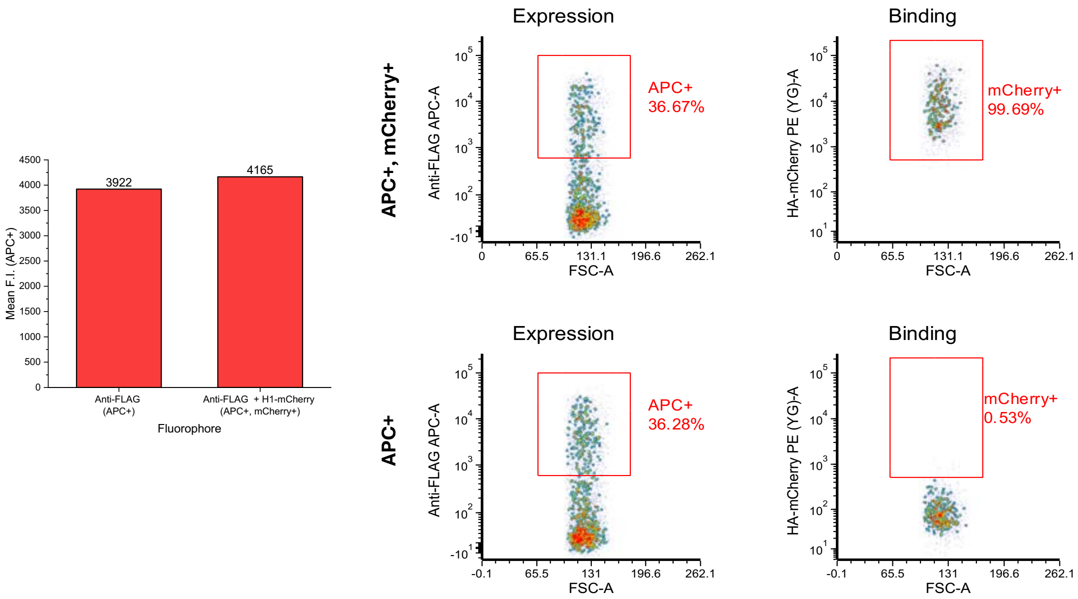
Mean F.I. (APC+)
3922
4165
Anti-FLAG
(APC+)
Anti-FLAG + H1-mCherry
(APC+, mCherry+)
Fluorophore
APC+, mCherry+
Expression
Anti-FLAG APC-A
FSC-A
APC+
36.67%
Binding
HA-mCherry PE (YG)-A
FSC-A
mCherry+
99.69%
APC+
Expression
APC+
36.28%
Binding
mCherry+
0.53%

**C**

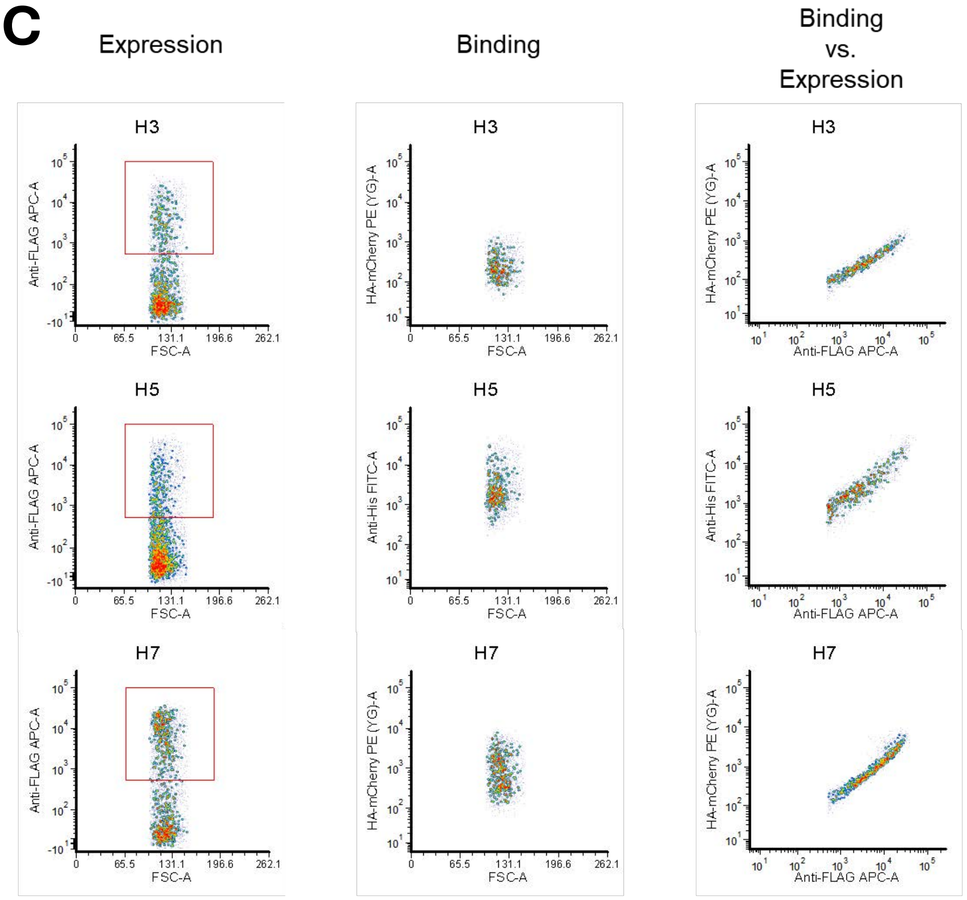
Expression
Binding
Binding vs. Expression
H3
H5
H7
Anti-FLAG APC-A
FSC-A
HA-mCherry PE (YG)-A
Anti-His FITC-A
Anti-FLAG APC-A

**D**

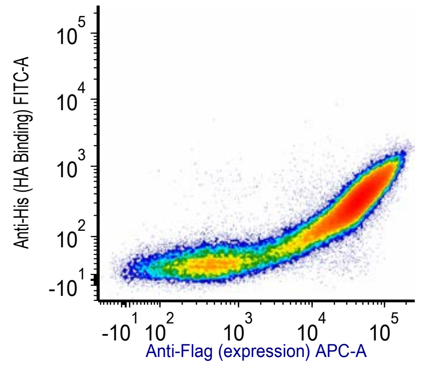
Anti-His (HA Binding) FITC-A
Anti-Flag (expression) APC-A

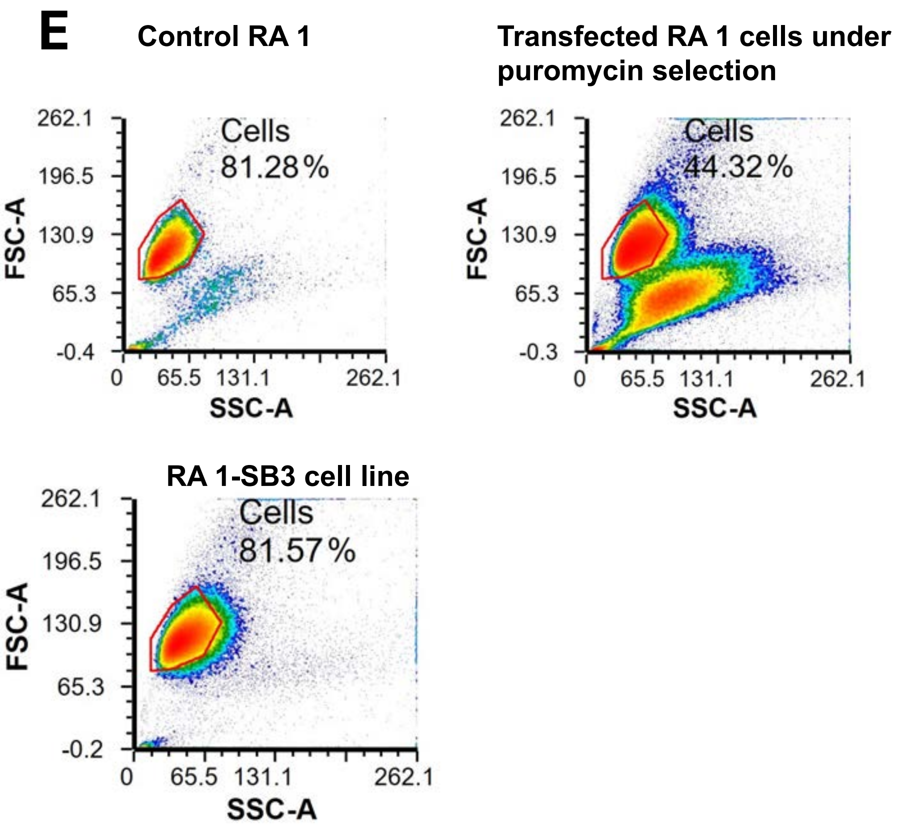


**Supplementary Figure 13: FACS of wild-type CR9114 binding variants of hemagglutinin.** (**A**) Density-corrected FACS dot plots for Expi 293 cells transfected with pSB3 and individual expression, binding, and binding vs. expression plots when incubated with H1-mCherry. (**B**) Data showing that the under the gating and sample preparation conditions we developed, the mCherry signal does not significantly contribute to the mean fluorescent intensity of the APC signal. (**C**) Denisty-corrected FACS dot plots for Expi 293 cells transfected with pSB3 and individual expression, binding, and binding vs. expression plots when incubated with H3, H5, and H7. (**D**) FACS dot plot for RA 1-SB3 cell line showing binding to H5. (**E**) FSC-A vs SSC-A plots for RA 1 cell lines, transfected RA 1 cell lines at the stage of puromycin selection and RA 1-SB3 cell lines post selection and cell line generation.

**A**

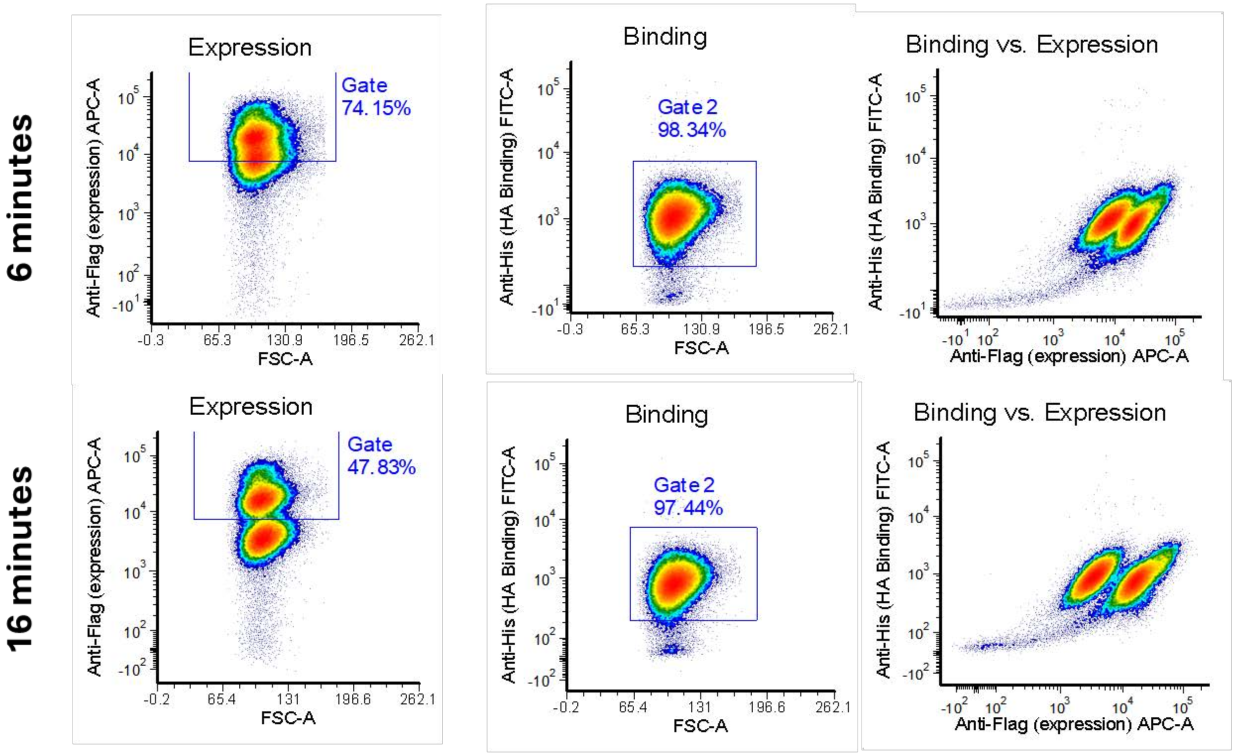

6 minutes
16 minutes
Expression
Gate
74.15%
47.83%
Anti-Flag (expression) APC-A
FSC-A
Binding
Gate 2
98.34%
97.44%
Anti-His (HA Binding) FITC-A
Binding vs. Expression


**B**

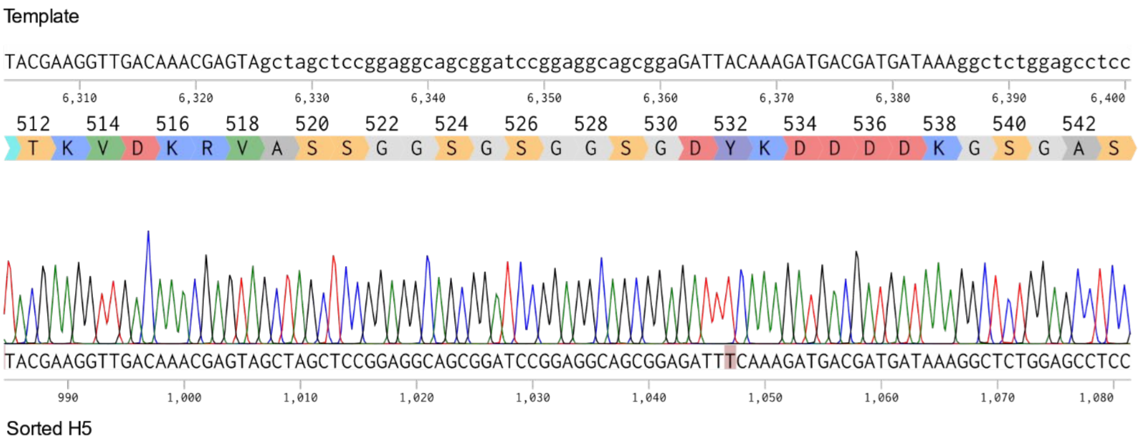

Template
TACGAAGGTTGACAAACGAGTAgctagctccggaggcagcggatccggaggcagcggaGATTACAAAGATGACGATGATAAAggctctggagcctcc
512 514 516 518 520 522 524 526 528 530 532 534 536 538 540 542
T K V D K R V A S S G G S G S G G S G D Y K D D D D K G S G A S
TACGAAGGTTGACAAACGAGTAGCTAGCTCCGGAGGCAGCGGATCCGGAGGCAGCGGAGATTTCAAAGATGACGATGATAAAGGCTCTGGAGCCTCC
Sorted H5

C

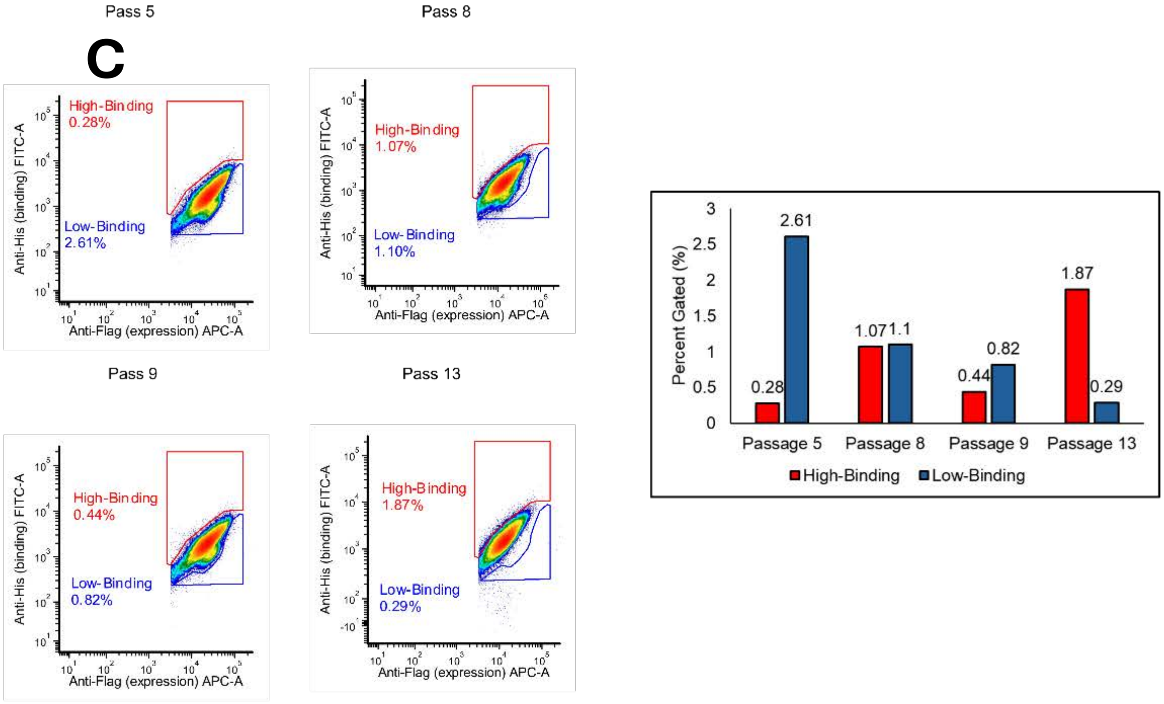


**Supplementary Figure 14: FACS sorting and detection of variant Y532F**. (**A**) Density-corrected FACS dot plots for enriched RA1-SB4 and their corresponding changes in the expression (anti-FLAG binding) during the course of the sort: Binding channel did not change magnitude, but expression channel shows population that decreased magnitude during the sort. (**B**) Chromatogram revealing the nucleotide substitution detected through its phenotype by FACS. (**C**) Bar charts of high-binding and low-binding populations as they change over the course of passaging RA1-SB4. Bar chart of high- (red) and low- binding (blue) populations.

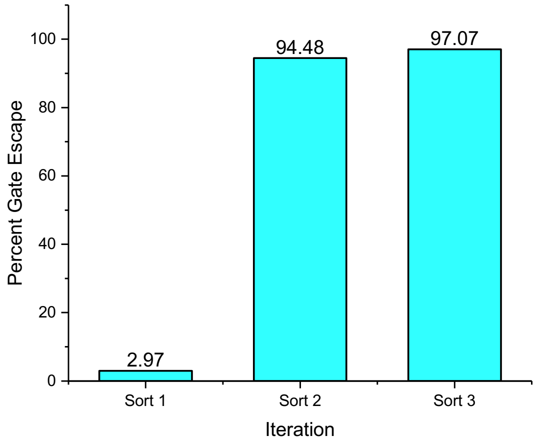


**Supplementary Figure 15: Iterative FACS sorting of high-binding population in RA1-SB4.** Bar chart representing the Gate Escape frequency of each iterative sort.

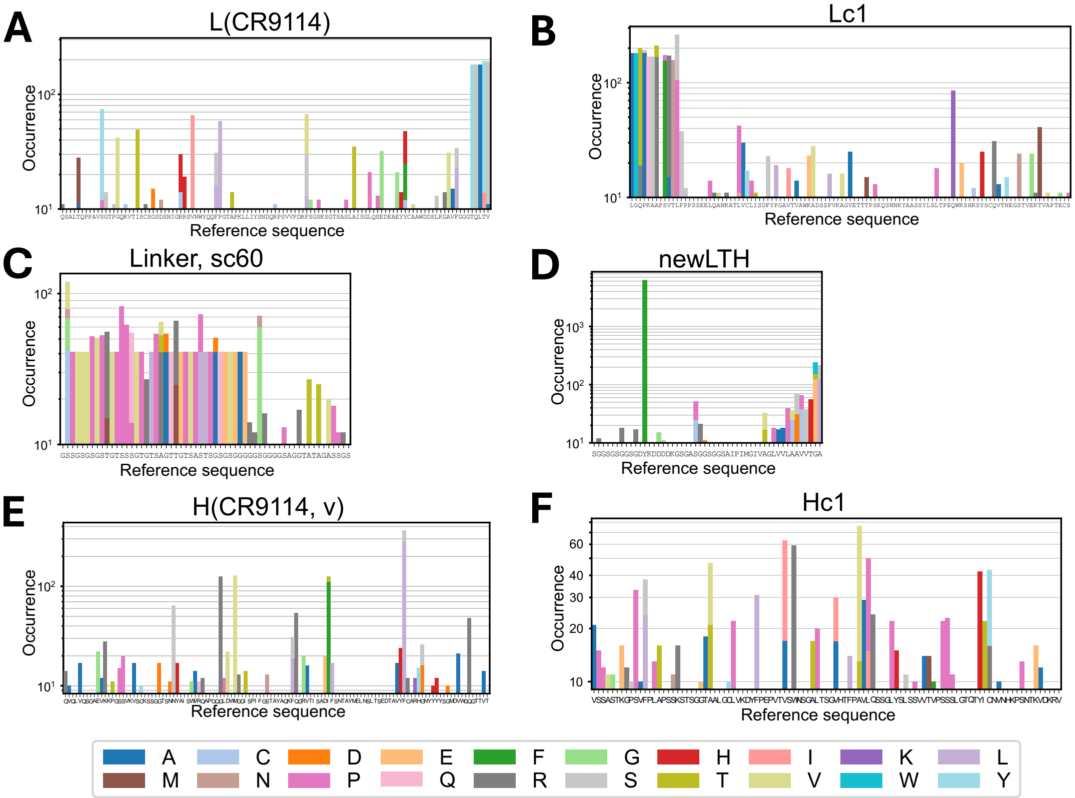


**Supplementary Figure 16: Mutational distribution of CR9114**. (A) Light chain variable region. (B) Light chain constant region. (C) Sc60 linker region. (D) linker-FLAG-MHCI helix region (E) Heavy chain variable region. (F) Heavy chain constant region. The variant labels are as per our $F_{ab}$ display sequence, see Supplementary Table 7 for Kabat numbering.

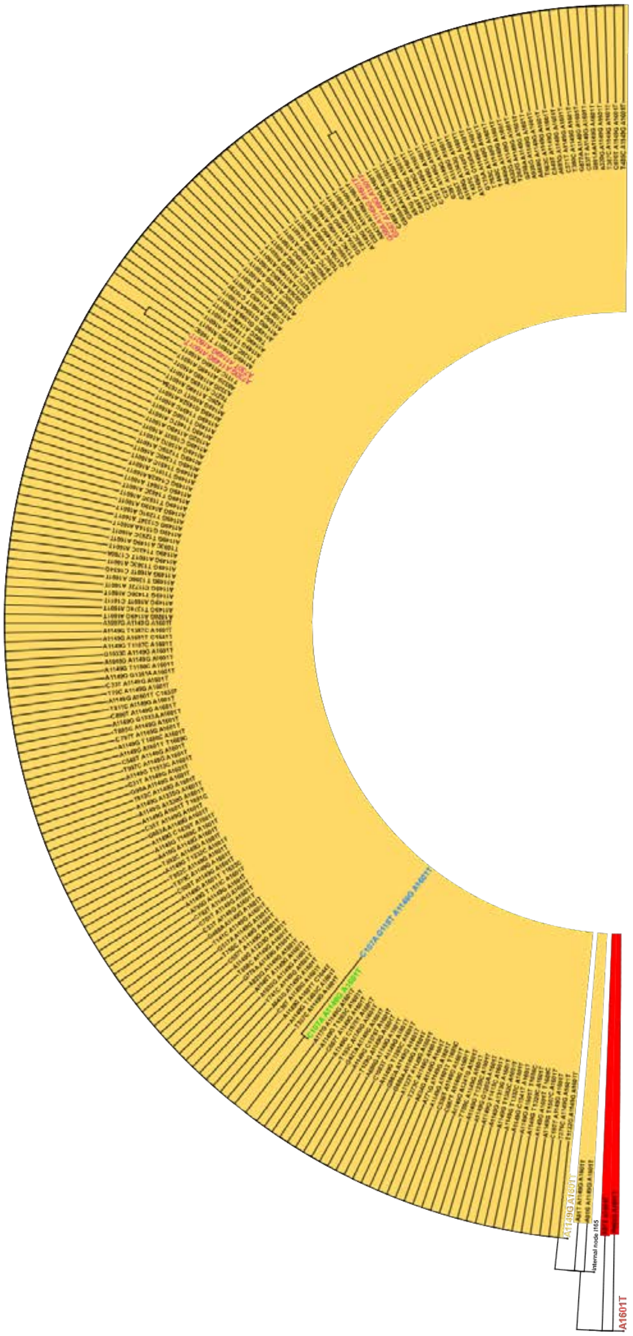

**Supplementary Figure 17: The phylogenetic tree of the Fab library.** Color codes corresponding to the number of mutations (1 red; 2 yellow; 3 green; 4 blue) a mutant harbor and which of them are observed in the subsequent lineages.

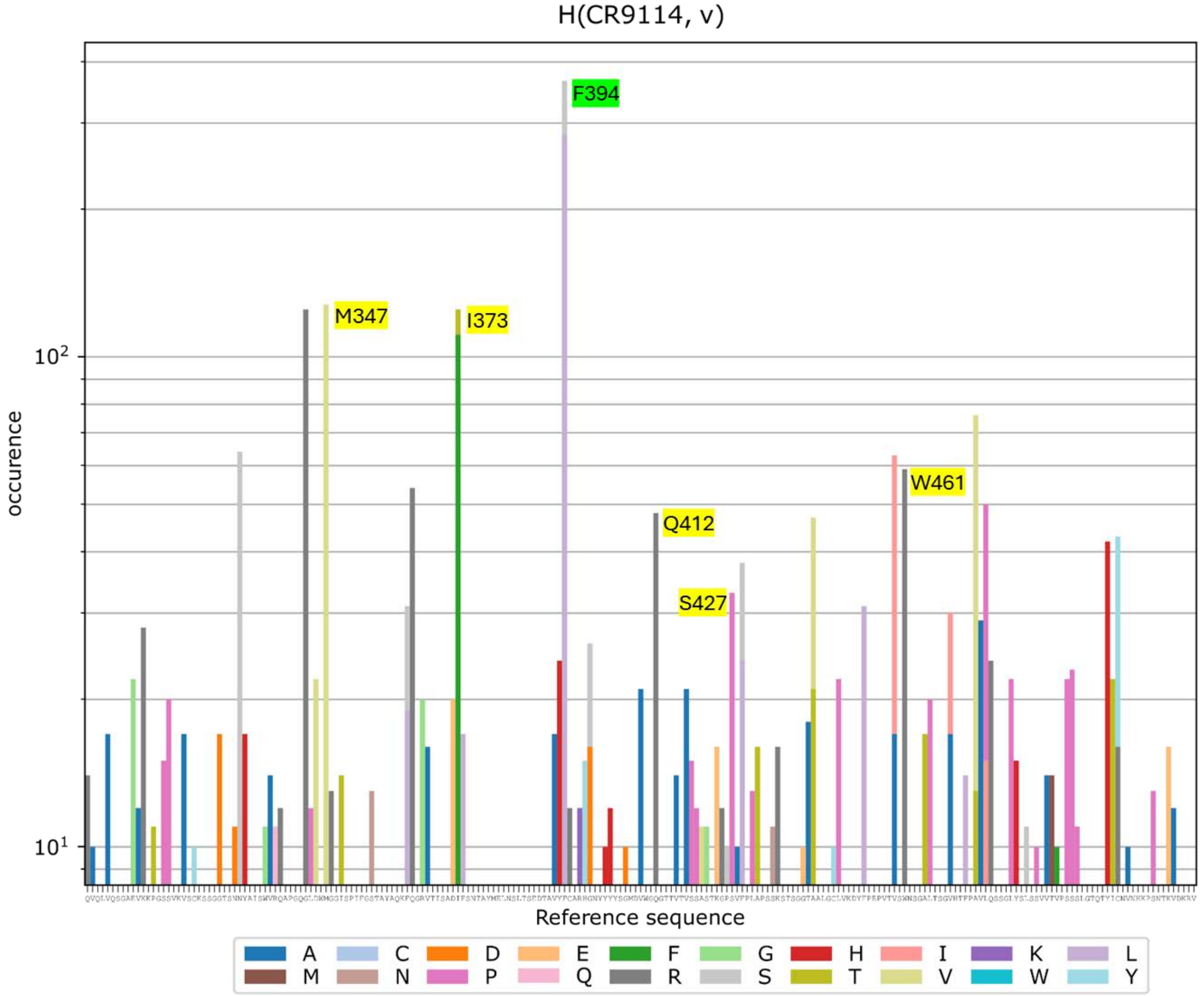


**Supplementary Figure 18:** Mutational distribution of CR9114 heavy chain, with highlighted mutants that showed lower EC50 values, relative to wild-type sequence. The variant labels are as per our $F_{ab}$ display sequence, see Supplementary Table 7 for Kabat numbering.

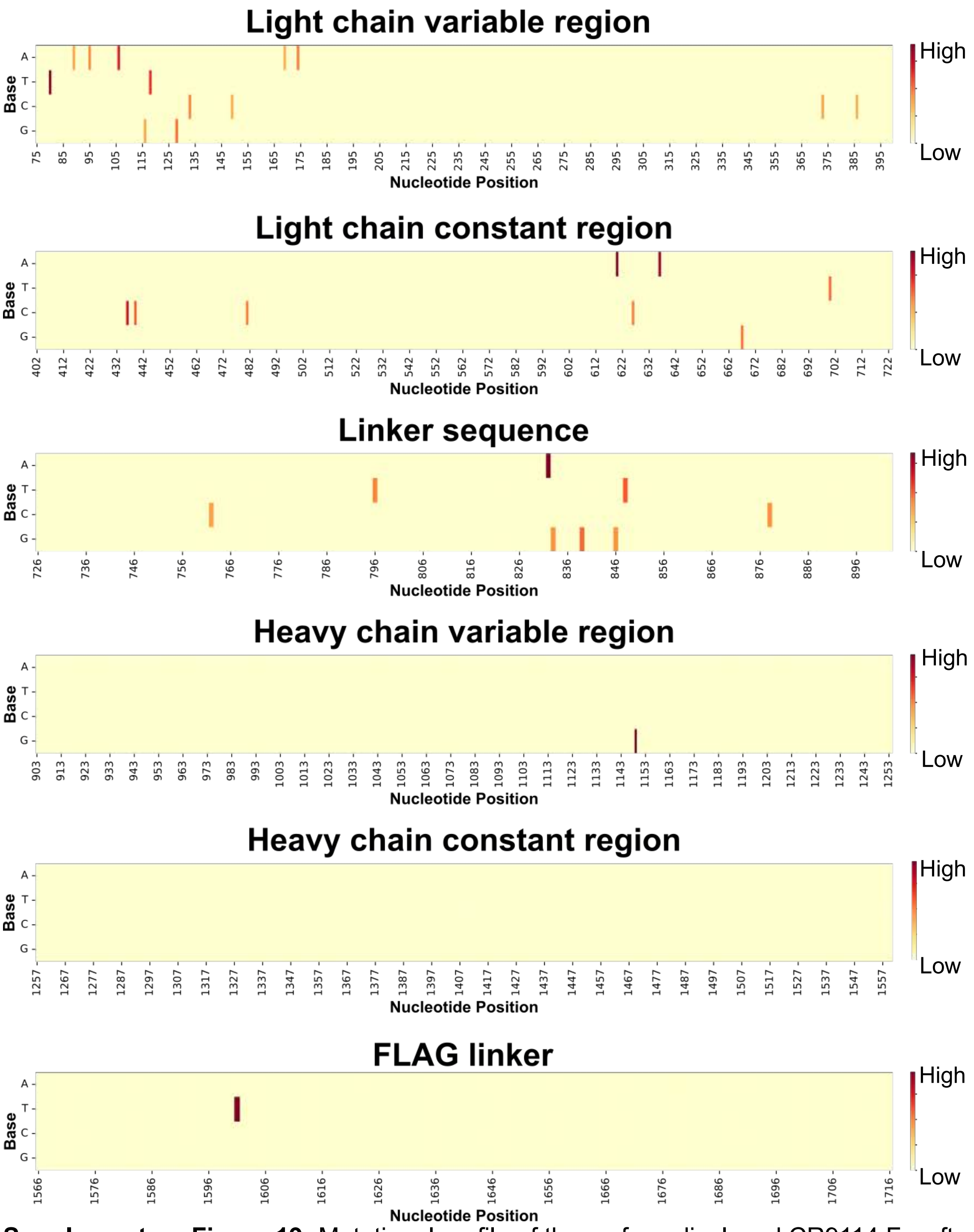


**Supplementary Figure 19:** Mutational profile of the surface displayed CR9114 $F_{ab}$ after 1 round of sorting and enrichment.

**Common Deletion Mutations**

| Mutation Identities | Mutation per 500,000 reads |
|---|---|
| deletion: transmembrane domain 52-55 (AAVA) | 4864 |
| deletion: transmembrane domain 49-54 (VTGAAV) | 1061 |
| deletion: VK 174 (N) | 36 |
| deletion: VH 100a (Y) | 18 |

**Common Single Substitutions in CDRs**

| Mutation Identity | Mutation per 500,000 reads |
|---|---|
| VH N82aK | 283243 |
| transmembrane domain D16V | 281930 |
| VK I21M | 220 |
| VH D46H | 125 |
| VH W154R | 46 |

**Common 2< Substitutions Mutations**

| Mutation Identities | Mutation per 500,000 reads |
|---|---|
| VH N82aK; transmembrane domain D16V | 266997 |
| VH N82aK; transmembrane domain D16V; transmembrane domain A55D | 7146 |
| VK I21M; VHN82aK; transmembrane domain D16V | 220 |
| VH D46H; VHN82aK; transmembrane domain D16V | 125 |
| VH N82aK; VH W154R; transmembrane domain D16V | 46 |

**Supplementary Figure 20: Summary of amino acid level mutations found across CR9114 transcript**. Recurrent in-frame deletions (span shown) are displayed alongside frequent single-nucleotide substitutions; reads with more than two amino-acid substitutions are also highlighted. Counts are normalized to 500,000 reads.

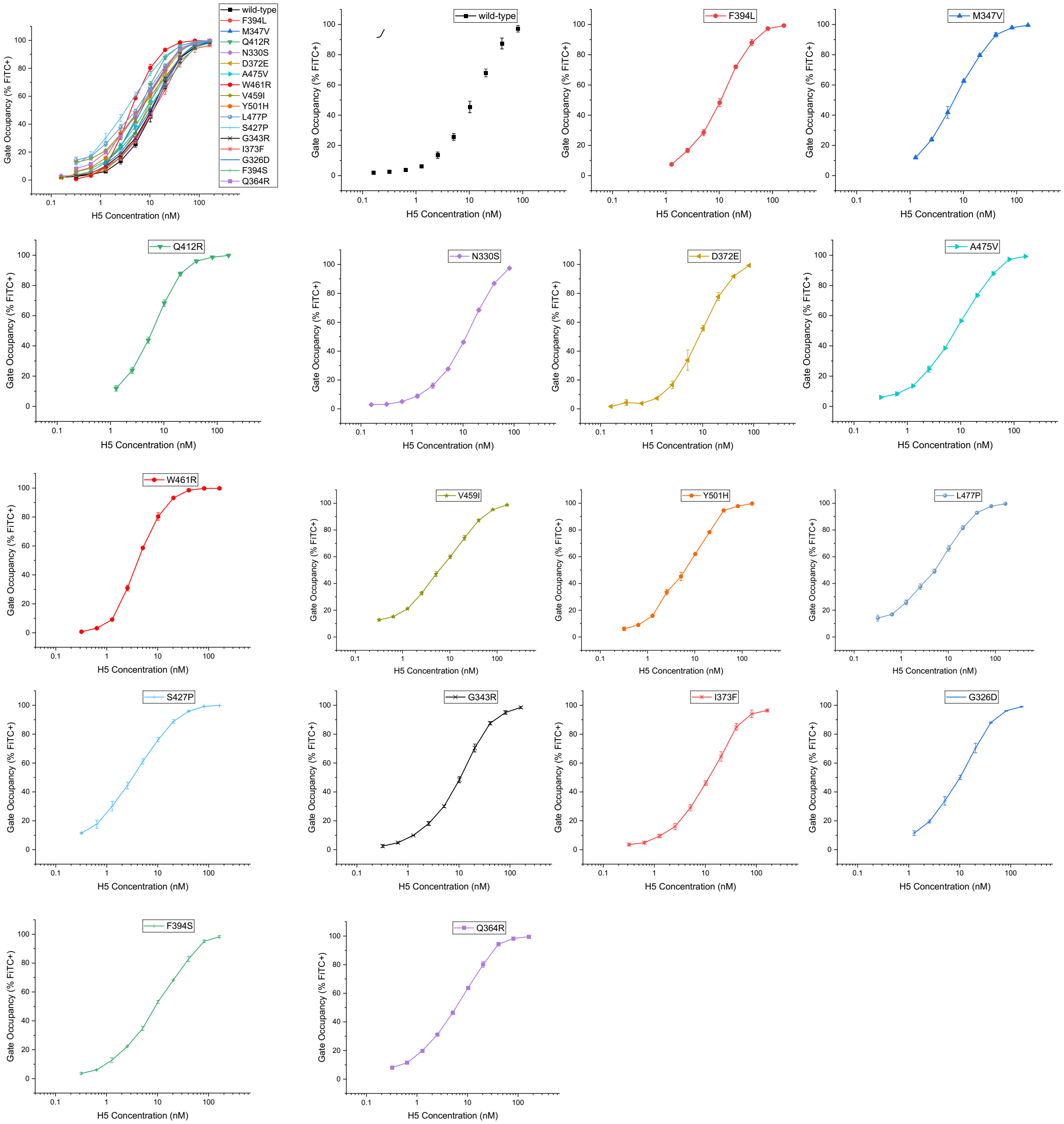


**Supplementary Figure 21: Analysis of H5 binding to evolved Fab variants**. Overlap of H5 binding of key variants is shown in top left panel. The individual traces showing H5 binding of key variants is shown in subsequent panels. Variations are listed in each panel. The variant labels are as per our $F_{ab}$ display sequence, see Supplementary Table 7 for Kabat numbering.

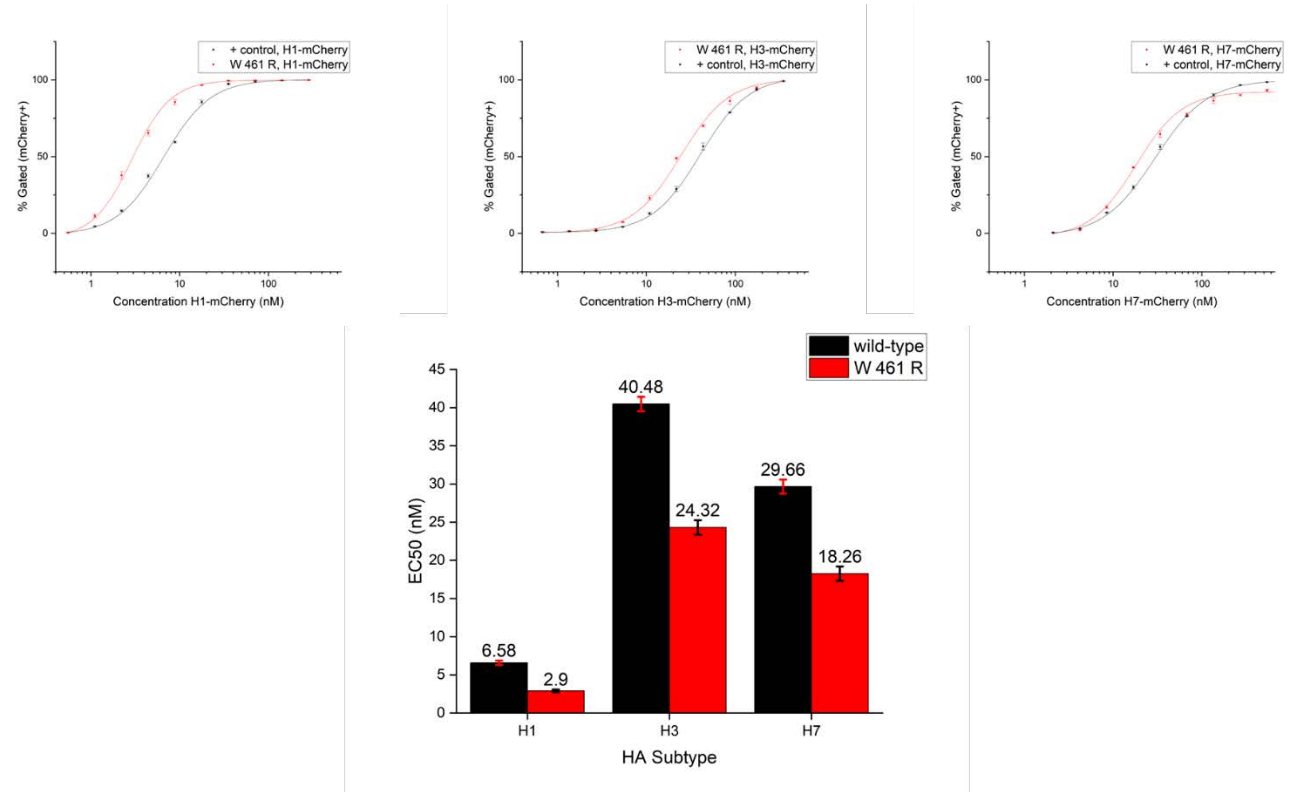


**Supplementary Figure 22:** Titration curves comparing the binding of wild-type CR9114 and the W461R mutant towards other HA variants (H1,H3,H7), along with a bar chart representing their EC50 values. The variant labels are as per our $F_{ab}$ display sequence, see Supplementary Table 7 for Kabat numbering.

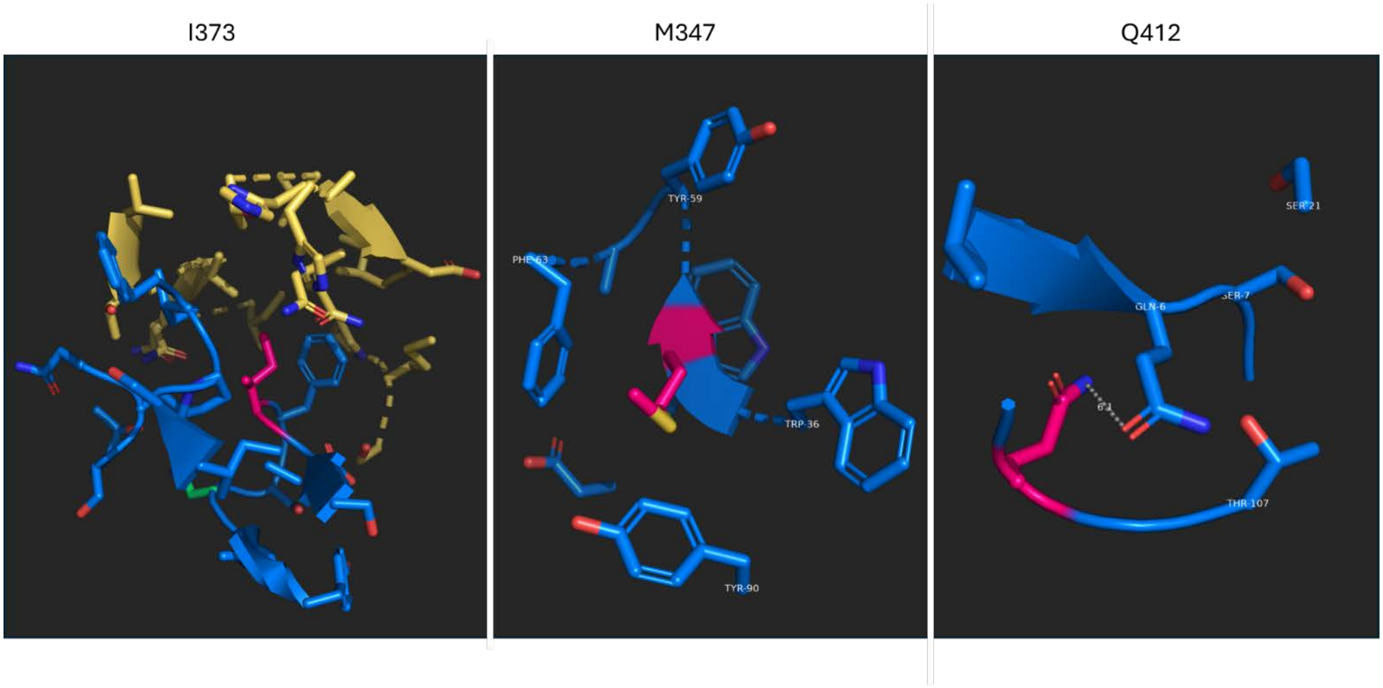


**Supplementary Figure 23:** The crystal structure of CR9114, with highlighted residues (pink) of I373, M347, Q412, the most enriched heavy chain variable region variants with the lowest EC50 values. The variant labels are as per our $F_{ab}$ display sequence, see Supplementary Table 7 for Kabat numbering.

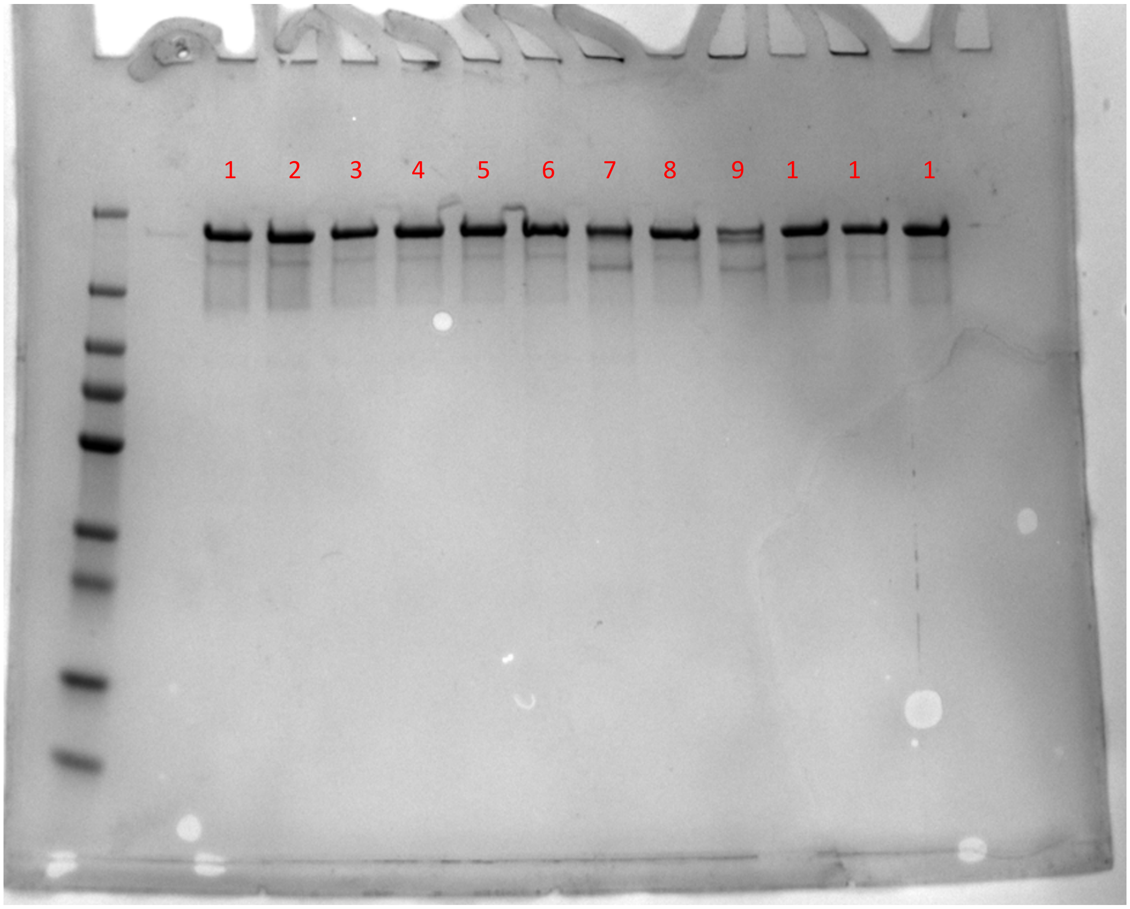


**Supplementary Figure 24: SDS-PAGE of purified CR9114 variants.** Each sample was diluted down to 1.5 ug and run in parallel. From left to right (1-12) the purified CR9114 samples are: 1. WT, 2. VH W154R, 3. VH G27D, 4. VH N31S, 5. VH G44R, 6. VH M48V, 7. VH Q64R, 8. VH D72E, 9. VH D72E-VH I73F–VH F74L, 10. VH I73F, 11. VH F74L, 12. VH F91L.

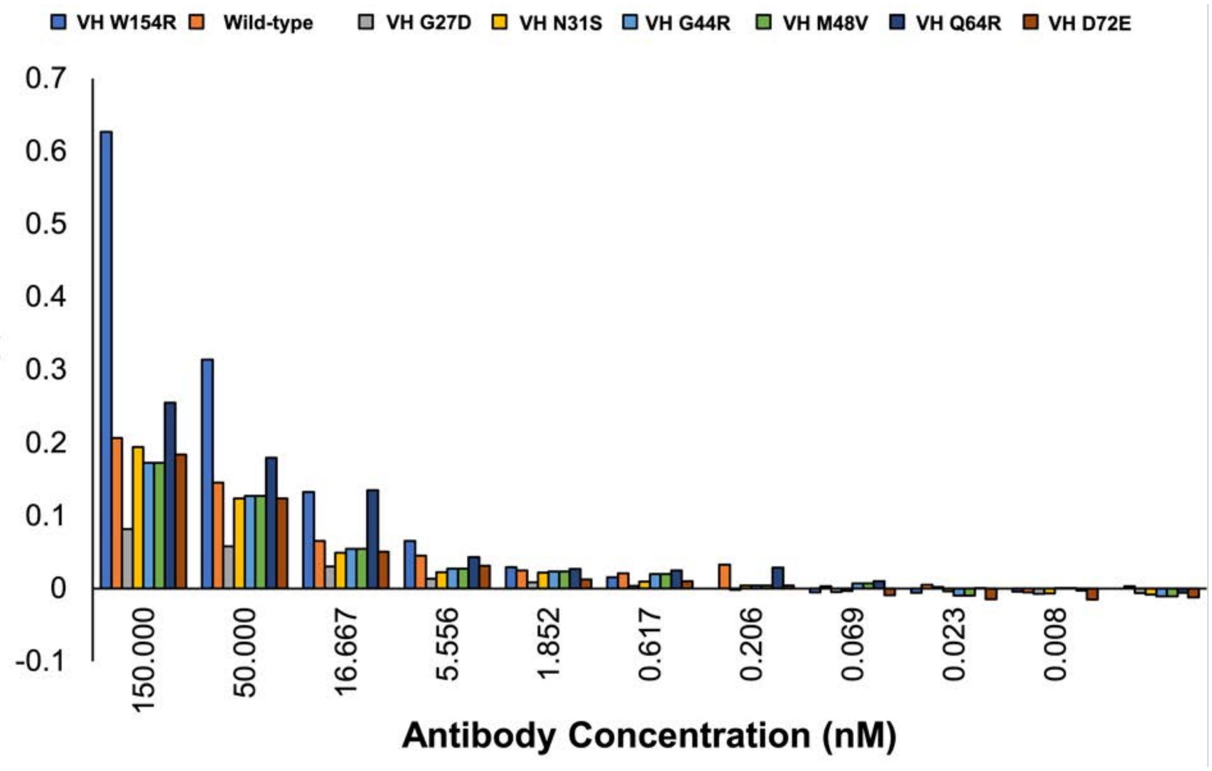


**Supplementary Figure 25: ELISA binding of purified mAbs to H5 hemagglutinin (A/Vietnam/1194/2004).** Dose–response plots from two-fold serial dilutions are shown to summarize relative binding strength of CR9114 variants.

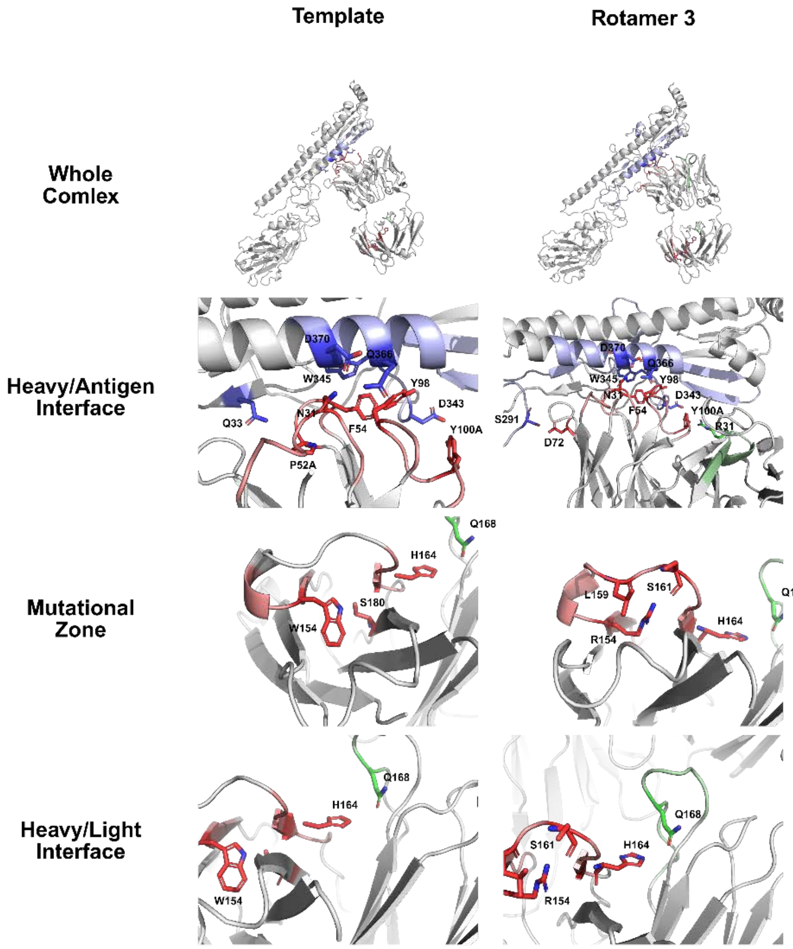


**Supplementary Figure 26: Computationally modeled interactions between H5 HA (blue), CR9114 heavy chain (red), and CR9114 light chain (green).** To understand how they key variant W154R alters the structure of CR9114 antibody, the crystal structure of the CR9114 Fab in complex with the H5 hemagglutinin (PDB ID: 4FQI) was imported into UCSF Chimera, and tryptophan at position 154 was individually substituted with each of 10 predicted arginine rotamers. All non-protein ligands and water molecules were removed prior to energy minimization, which was performed using Chimera’s default parameters. Following minimization, interfacial hydrogen-bond networks were analyzed using Chimera’s built-in hydrogen-bond detection tool, and subsequent docking with HADDOCK revealed minimal change in the interfacial binding energy, given by the HADDOCK score, between the wild type (-132.0 +/- 4.5 units) compared to the W154 variant (-130.6 +/- 1.8 units). Previous studies suggests that the binding contributions made by the light chain also play an important role in antigen binding. While CR9114 exclusively binds to H5 HA using its CDRH loops, significant bond shortening was observed for the preserved hydrogen bond between heavy chain H164 and light chain Q168; this light chain residue is connected to a broader hydrogen bond network in the $L_c$ region that interfaces with the $L_v$ region and potentially transfers this stability to the light chain residue R31, which stabilizes the configuration of CDRH residues.

**Supplementary Table 1: Oligonucleotides used in this study.**

| Name | Sequence (5'-3') | Purpose |
|---|---|---|
| **SB 168A** | AAACGGCTCGAGGTTCTCAGAGGGCACAGCCAG | Amplify proXIV1 & proXIV2 from RA1 genome; add homology for p274 |
| **SB 168B** | GGGCACCGGAGCCGTTCTTGTGCAGGAGGTCCA | Amplify proXIV1 & proXIV2 from RA1 genome; add homology for p274 |
| **SB 169A** | CCTGCACAAGAACGGCTCCGGTGCCCGTCAGTG | Amplify p274 backbone; add homology for proXIV1 & proXIV2 |
| **SB 169B** | TGCCCTCTGAGAACCTCGAGCCGTTTCTTGCAG | Amplify p274 backbone; add homology for proXIV1 & proXIV2 |
| **SB 413A** | GCTGCGCCTTATCCGGTAACTATCGT | Amplify p274 backbone; add homology for Origin of Replication |
| **SB 413B** | ACGATAGTTACCGGATAAGGCGCAGC | Amplify p274 backbone; add homology for Origin of Replication |
| **SB 246A** | CTTGATGTTTGTTTTTCAATAGCAA CTAATGCCTACGCTCAATCAGCTCTC ACGCAACC | Amplify Fab-CR9114; add homology for secretion signal (MLS) |
| **GL 7B** | GCCACCATGAAGAAGAACATAGCGTTTTTG CTCGCCTTGATGTTTGTTTTTCAATAG CA | Amplify Fab-CR9114 SB 246A Amplification; add further homology for secretion signal (MLS) |
| **SB 274A** | AGGGGATCGTCGACCGTACGGCCACCATGAAGAAGAA CATAGCGT | Amplify Fab-CR9114 GL |

| | | |
|---|---|---|
| | | 7B Amplification; add final homology for secretion signal (MLS) |
| **SB 274B** | CCGCTGCCTCCGGAGCTAGCTACTCGTTTGTCAAC CTTCGTATTT | Amplify Fab-CR9114 SB with compete MLS; add homology for p274 |
| **SB 275A** | GCTAGCTCCGGAGGCAGCGGATCCGGAGGCAGCGGAGATT ACAAAGATGACGATGATAAA | Amplify Linker-Tag-MHCI helix; add homology for Fab-CR9114 |
| **SB 252B** | TGCACCTGAGGAGTGCGGCCGCTTTAATCTGAGCTCTTCTTTCTCCA CAGC | Amplify Linker-Tag-MHCI helix; add homology for p274 |
| **SB 209A** | AGCGGCCGCACTCCTCAGGTGCAGG | Amplify p274; add homology for Linker-Tag-MHCI helix |
| **SB 275B** | GGTGGCCGTACGGTCGACGATCCCCTCACGACACCTGAAATGGA AGAAAA | Amplify p274; add homology for MLS |
| **SB 313A** | GGATGGGGTTGGGGGATGCG | Amplify DIVAC; add homology for p274 |
| **SB 314B** | TTTTCAGTTTCGGTCAGCCTCGCCT | Amplify DIVAC; add homology for p274 |
| **SB 316B** | CGCATCCCCCAACCCCATCCCCTCGAGCCGTTTCTTGCAGGAATAC | Amplify p274; add homology for DIVAC |
| **SB 322A** | AGGCGAGGCTGACCGAAACTGAAAAGGCTCCGGTGCCCGTCAGTG | Amplify p274; add homology for DIVAC |
| **SB 171A** | ATCTACGGCGTGCAGTGCTTCAG | Amplify 274-eGFP*; introduce knockout mutation into |

| | | |
|---|---|---|
| | | eGFP; blunt end ligation |
| **SB 171B** | CAGGGTGGTCACGAGGGTGG | Amplify 274-eGFP*; blunt end ligation |
| **SB 408A** | TCAGGATGCCTTCTATATCCTCAGCTTCTC | Confirm 5' Integration of p274-based cassettes |
| **SB 408B** | AATTGCATATGCTAAGTGTGTGAATGAAGT | Confirm 3' Integration of p274-based cassettes |
| **SB 212B** | CCTGCACCTGAGGAGTGCGGCCGCT | Confirm Integration of pp274-based cassettes |
| **DJO 124** | TCTGGATGTCAGCGTAGGCATTAGTTGCTATTGAAAAAACA | Amplify p274; add homology for 047-09_1A02 |
| **DJO 408** | GCCAAAATCATCCGGAGGCAGCGGATCCGGAGGCAGCGGAGATTACAAAG | Amplify p274; add homology for 047-09_1A02 |
| **DJO 475** | AATGCCTACGCTGACATCCAGATGACTCAGTCTCCCTCCAGCCTC | Amplify 047-09_1A02; add homology for p274 |
| **DJO 476** | TGCCTCCGGATGATTTTGGCTCAACTTTTTTGTCCACTTTGGTGTTTG | Amplify 047-09_1A02; add homology for p274 |
| **DJO 725** | AAACGGCTCGAGGTGAATTGGTAAATATGTGGGTACGAATTCTGG | Amplify pro862; add homology for p274 |
| **DJO 726** | ATTTACCAATTCACCTCGAGCCGTTTCTTGCAGGAATACAAGAAA | Amplify p274; add homology for pro862 |
| **DJO 727** | CTCACTTAGGCACGGCTCCGGTGCCCGTCAGTGGGCAGAGCGCA | Amplify p274; add homology for pro862 |
| **DJO 728** | GGGCACCGGAGCCGTGCCTAAGTGAGGATTTGCATGTGGGTGGTG | Amplify pro862; add homology for p274 |
| **DJO 729** | AAACGGCTCGAGGCACAGTGTGGAAACCCACATCCCGAGAGTTTC | Amplify pro936; add |

| | | |
|---|---|---|
| | | homology for p274 |
| **DJO 730** | TTTCCACACTGTGCCTCGAGCCGTTTCTTGCAGGAATACAAGAAA | Amplify p274; add homology for pro936 |
| **DJO 731** | ATCAACCCAACAAGGCTCCGGTGCCCGTCAGTGGGCAGAGCGCAC | Amplify p274; add homology for pro936 |
| **DJO 732** | GGGCACCGGAGCCTTGTTGGGTTGATGCTGCTTCTTCAGAGGGGA | Amplify pro936; add homology for p274 |
| **DJO 733** | AAACGGCTCGAGGGTGAGGAGTTGTTTAAATTCCCCTCTGAAGAAG | Amplify pro1417; add homology for p274 |
| **DJO 734** | AACAACTCCTCACCCTCGAGCCGTTTCTTGCAGGAATACAAGAAA | Amplify p274; add homology for pro1417 |
| **DJO 735** | AGTGTCATTGTCCGGCTCCGGTGCCCGTCAGTGGGCAGAGCGCAC | Amplify p274; add homology for pro1417 |
| **DJO 736** | GGGCACCGGAGCCGGACAATGACACTCAAACCCAGAATTCGTACC | Amplify pro1417; add homology for p274 |

**Supplementary Table 2: Synthesized DNA fragments**. Commercially purchased synthesized DNA fragments used in this study.

| Synthesized DNA Fragment | Sequence (5'-3') |
|---|---|
| CR9114 Fab | CAATCAGCTCTCACGCAACCGCCAGCTGTTTCCGGTACCCCA<br>GGTCAACGCGTTACCATATCATGTAGCGGCTCTGACTCAAAT<br>ATTGGTCGCCGCTCCGTAAACTGGTACCAACAGTTTCCCGGG<br>ACAGCTCCGAAACTCCTGATCTATAGCAACGATCAACGCCCG<br>TCAGTGGTACCAGACAGGTTTAGTGGCTCTAAATCCGGAACA<br>TCAGCTTCTCTCGCTATCAGCGGTCTCCAATCAGAAGACGAA<br>GCTGAGTACTACTGCGCAGCTTGGGATGACTCCTTGAAGGGA<br>GCTGTCTTCGGCGGGGGTACTCAGCTTACAGTGCTTGGACAG<br>CCCAAAGCCGCCCCTTCCGTAACTCTTTTCCCCCCATCAAGC<br>GAGGAACTTCAAGCAAATAAGGCAACCCTTGTCTGCCTGATT<br>AGTGATTTTTATCCGGGTGCCGTTACAGTGGCATGGAAAGCA<br>GACTCTAGTCCAGTCAAGGCAGGAGTTGAGACAACTACGCCA<br>TCCAAGCAATCCAACAACAAATACGCTGCCTCAAGTTATTTG<br>AGCCTTACCCCAGAGCAATGGAAAAGTCATCGCTCATATTCT<br>TGTCAGGTCACACATGAAGGCAGCACAGTGGAAAAGACGGT<br>TGCGCCAACGGAATGTTCCGGGGGGAGTTCCGGTAGTGGTTC<br>CGGTTCCACGGGTACCTCCTCCTCAGGTACCGGGACTTCCGC<br>GGGCACAACGGGAACCAGTGCCTCTACTTCCGGCTCAGGAA<br>GTGGAGGCGGAGGGGGGAGCGGAGGCGGTGGATCCGCGGG<br>CGGAACCGCTACCGCTGGGGCGTCATCCGGATCTCAAGTAC<br>AGCTCGTTCAATCAGGAGCTGAGGTGAAAAAACCCGGTTCT<br>TCTGTAAAAGTTAGCTGCAAATCATCTGGAGGCACTAGCAA<br>CAACTACGCTATAAGTTGGGTACGACAAGCTCCAGGGCAA<br>GGTCTGGACTGGATGGGAGGCATTTCACCGATATTCGGGAG<br>CACAGCTTACGCTCAGAAGTTTCAGGGCAGGGTAACCATCT<br>CAGCTGATATTTTTTCTAACACAGCTTACATGGAACTCAAT<br>AGTCTCACATCAGAAGACACAGCTGTGTACTTTTGTGCTA<br>GGCATGGCAATTATTACTACTACAGTGGCATGGATGTATGG<br>GGCCAGGGAACAACAGTTACAGTCTCTTCAGCCAGTACCAA<br>GGGGCCCTCCGTCTTTCCTCTCGCCCCGAGTAGCAAAAGCA<br>CTTCAGGGGGGACTGCAGCACTCGGATGCCTGGTTAAAGA<br>TTACTTTCCAGAACCTGTCACCGTATCCTGGAATAGTGGCG<br>CTCTCACTAGCGGCGTACATACGTTCCCGGCGGTCCTCCAG<br>AGTAGCGGTCTGTATAGTTTGAGTTCTGTCGTAACGGTTCC<br>CTCATCTTCTCTGGGTACACAGACCTATATCTGCAATGTGAA<br>TCATAAACCATCAAATACGAAGGTTGACAAACGAGTA |
| Linker-FLAG-MHCI helix | GCTAGCTCCGGAGGCAGCGGATCCGGAGGCAGCGGAGATTA<br>CAAAGATGACGATGATAAAGGCTCTGGAGCCTCCGGAGGCA<br>GCGGAGGAAGTGCTATCCCCATCATGGGTATCGTTGCTGGCC<br>TGGTTGTCCTTGCAGCTGTAGTCACTGGAGCTGCGGTCGCTG<br>CTGTGCTGTGGAGAAAGAAGAGCTCAGATTAA |

| | |
|---|---|
| 047-09_1A02 Fab | GACATCCAGATGACTCAGTCTCCCTCCAGCCTCAGTGCCTCCGTGGGTGACCGA<br>GTCACTATCACATGCCGAGCTAGCCAAGACATCAACAATAATTTGAACTGGTATCA<br>ACAGAAGCCAGGAAAAGCCCCCAAATTGCTTATGTACGACGCATCTAATTTGGAA<br>GCTGGCGTCCCTTCTCGCTTCAGTGGCACTGGGTCCGGCACCGACTTCACCTTT<br>ACTATCTCCTCCCTTCAGGCGGAGGACATAGCGACTTACTATTGTCAGCAATATAG<br>GAATCCACTTCTCACGTTCGGAGGTGGAACTAAGGTGGAGATAAAACGGACCGT<br>GGCAGCGCCGTCTGTCTTCATTTTTCCCCCTAGCGACGAGCAGCTCAAATCAGG<br>GACAGCTAGCGTTGTGTGTCTGTTGAACAATTTTTACCCAAGGGAAGCCAAAGTT<br>CAATGGAAGGTGGATAACGCCCTCCAGAGTGGAAATAGTCAAGAAAGTGTGACT<br>GAACAGGATAGTAAGGATTCCACTTATAGTCTTTCATCTACGTTGACGCTGTCTAA<br>GGCCGACTATGAGAAACATAAGGTCTACGCATGTGAGGTAACACATCAGGGGTTG<br>TCATCTCCAGTAACAAAATCCTTCAACAGGGGGGAGTGCGGGGGGAGTTCCGGT<br>AGTGGTTCCGGTTCCACGGGTACCTCCTCCTCAGGTACCGGGACTTCCGCGGG<br>CACAACGGGAACCAGTGCCTCTACTTCCGGCTCAGGAAGTGGAGGCGGAGGGG<br>GGAGCGGAGGCGGTGGATCCGCGGGCGGAACCGCTACCGCTGGGGCGTCATC<br>CGGATCTGAGGTGCAACTCCTTGAGAGCGGAGGAGCGCTGGTACAGCCTGGGG<br>GCTCACTGCGCCTGTCTTGCACCGCAAGTGGCTTCACTCTTGGTCATTTCGCGA<br>TGGCGTGGGTGAGGCAAGCGCCTGGCAAAGGGTTGGAATGGCTGTCAGCTATA<br>AGCGGCGGAGGTGGGACCACGTACTATGCCGATTCCGTCAAGGGGCGATTTACT<br>ATTAGTAGGGACAATTCAAAGAATACTCTGTACTTGCAGCTTTCTGGCTTGAGGGC<br>AGAGGACACCGCCCTTTACTTTTGTGGTAAGTACGATAGTTCAGGGCACCACTAT<br>GTAAGGCGCATGCATTTTTGGGGACAAGGTACGCTTGTCACTGTCTCCTCAGCCT<br>CCACAAAAGGGCCGAGCGTTTTCCCTCTTGCGCCCAGTAGCAAGTCCACATCTG<br>GCGGCACAGCCGCTTTGGGATGCCTTGTAAAGGATTACTTTCCCGAACCAGTGA<br>CTGTCAGTTGGAACTCTGGTGCTCTTACTAGCGGTGTCCATACATTCCCAGCGGT<br>ACTGCAGAGCTCCGGGCTCTATTCACTCTCAAGTGTAGTCACTGTGCCAAGCTCT<br>AGCTTGGGTACCCAAACGTATATCTGTAATGTAAATCATAAGCCTTCAAACACCAA<br>AGTGGACAAAAAAGTTGAGCCAAAATCA |

**Supplementary Table 3: Plasmid DNA constructs used in this study**.

| Plasmid Name | Integration Cassette |
|---|---|
| pSB1 | proXIV-1-EF1a-eGFP*-PGK-puroR |
| pSB2 | EF1a-eGFP*-PGK-puroR |
| pSB3 | EF1a-CR9114-PGK-puroR |
| pSB4 | proXIV-1-EF1a-CR9114-PGK-puroR |
| pSB5 | proXIV-2-EF1a-eGFP*-PGK-puroR |
| pSB6 | DIVAC-EF1a-eGFP*-PGK-puroR |
| pSZ1 | proXIV-2-862-EF1a-eGFP*-PGK-puroR |
| pSZ2 | proVIX-2-1417-EF1a-eGFP*-PGK-puroR |
| pSZ3 | proXIV-2-936-EF1a-eGFP*-PGK-puroR |
| pDJO1 | proXIV-1-EF1a-047-09_1A02-PGK-puroR |

**Supplementary Table 4: B cell lines generated in this study**.

| B Cell Line | Plasmids Used | Integration Cassette |
|---|---|---|
| RA 1-SB1 | p276 + pSB1 | proXIV-1 EF1a-eGFP*-PGK-puroR |
| RA 1-SB2 | p276 + pSB2 | EF1a-eGFP*-PGK-puroR |
| RA 1-SB3 | p276 + pSB3 | EF1a-CR9114-PGK-puroR |
| RA 1-SB4 | p276 + pSB4 | proXIV-1 EF1a-CR9114-PGK-puroR |
| RA 1-SB5 | p276 + pSB5 | proXIV-2 EF1a-eGFP*-PGK-puroR |
| RA 1-SB6 | p276 + pSB6 | DIVAC-EF1a-eGFP*-PGK-puroR |
| RA 1-SZ1 | p276 + pSZ1 | proXIV-2-862-EF1a-eGFP*-PGK-puroR |
| RA 1-SZ2 | p276 + pSZ2 | proXIV-2-1417-EF1a-eGFP*-PGK-puroR |
| RA 1-SZ3 | p276 + pSZ3 | proXIV-2-936-EF1a-eGFP*-PGK-puroR |
| RA 1-DJO1 | p276 + pDJO1 | proXIV-1 EF1a-047-09_1A02-PGK-puroR |

**Supplementary Table 5. Table detailing the most enriched nucleotide substitution obtained through PacBio NGS analysis of the eGFP* library**

| Nucleotide Substitution | Read Frequency | Mutational Frequency (%) | Initial Codon | Initial Amino Acid | Residue Number | Substituted Amino Acid | Substituted Codon |
|---|---|---|---|---|---|---|---|
| T197C | 718103 | 58.52 | ATC | I | 66 | T | ACC |
| T197G | 125082 | 10.19 | ATC | I | 66 | S | AGC |
| G310A | 122486 | 9.98 | GAC | D | 104 | N | AAC |
| A596G | 94611 | 7.71 | AAC | N | 199 | S | AGC |
| T197A | 37893 | 3.09 | ATC | I | 66 | N | AAC |
| C208A | 28447 | 2.32 | CAG | Q | 70 | K | AAG |
| C127G | 23808 | 1.94 | CTG | L | 43 | V | GTG |
| T50A | 5845 | 0.48 | GTC | V | 17 | D | GAC |
| C108G | 1653 | 0.13 | GGC | G | 36 | G | GGG |
| A557C | 1634 | 0.13 | AAC | N | 186 | T | ACC |
| G58T | 1580 | 0.13 | GAC | D | 20 | Y | TAC |
| G14A | 1503 | 0.12 | GGC | G | 5 | D | GAC |
| C46A | 1452 | 0.12 | CTG | L | 16 | M | ATG |
| G13A | 1406 | 0.11 | GGC | G | 5 | S | AGC |
| G575T | 1348 | 0.11 | GGC | G | 192 | V | GTC |
| C646A | 1328 | 0.11 | CGC | R | 216 | S | AGC |
| T491C | 1268 | 0.10 | GTG | V | 164 | A | GCG |
| T701A | 1120 | 0.09 | ATG | M | 234 | K | AAG |
| G16A | 953 | 0.08 | GAG | E | 6 | K | AAG |
| G8A | 875 | 0.07 | AGC | S | 8 | N | AAC |

**Supplementary Table 6**. The average mutational rates per kb of eGFP* and CR9114 Fab Surface Display genes.

| | Average mutation count/per mRNA | Segment Length (kb) | Mutation Rate (Mutations / kb) |
|---|---|---|---|
| eGFP* | 2.10 | 0.76 | 2.76 |
| eGFP* naïve pool early passage | 0.287 | 0.586 | 0.490 |
| eGFP* naïve pool late passage | 0.315 | 0.586 | 0.538 |
| CR9114 Surface Display | 2.98 | 1.72 | 1.73 |

**Supplementary Table 7:** The variant labels are as per the structure, see Supplementary Table 7 for Kabat numbering.

| Heavy | | | Light | | |
|---|---|---|---|---|---|
| AA | Kabat position | Original position | AA | Kabat position | Original position |
| Q | 1 | 300 | Q | 1 | 24 |
| V | 2 | 301 | S | 2 | 25 |
| Q | 3 | 302 | A | 3 | 26 |
| L | 4 | 303 | L | 4 | 27 |
| V | 5 | 304 | T | 5 | 28 |
| Q | 6 | 305 | Q | 6 | 29 |
| S | 7 | 306 | P | 7 | 30 |
| G | 8 | 307 | P | 8 | 31 |
| A | 9 | 308 | A | 9 | 32 |
| E | 10 | 309 | - | 10 | |
| V | 11 | 310 | V | 11 | 33 |
| K | 12 | 311 | S | 12 | 34 |
| K | 13 | 312 | G | 13 | 35 |
| P | 14 | 313 | T | 14 | 36 |
| G | 15 | 314 | P | 15 | 37 |
| S | 16 | 315 | G | 16 | 38 |
| S | 17 | 316 | Q | 17 | 39 |
| V | 18 | 317 | R | 18 | 40 |
| K | 19 | 318 | V | 19 | 41 |
| V | 20 | 319 | T | 20 | 42 |
| S | 21 | 320 | I | 21 | 43 |
| C | 22 | 321 | S | 22 | 44 |
| K | 23 | 322 | C | 23 | 45 |
| S | 24 | 323 | S | 24 | 46 |
| S | 25 | 324 | G | 25 | 47 |
| G | 26 | 325 | S | 26 | 48 |
| G | 27 | 326 | D | 27 | 49 |
| T | 28 | 327 | S | 27A | 50 |
| S | 29 | 328 | N | 27B | 51 |
| N | 30 | 329 | I | 28 | 52 |
| N | 31 | 330 | G | 29 | 53 |
| Y | 32 | 331 | R | 30 | 54 |
| A | 33 | 332 | R | 31 | 55 |
| I | 34 | 333 | S | 32 | 56 |
| S | 35 | 334 | V | 33 | 57 |

| | | |
|---|---|---|
| W | 36 | 335 |
| V | 37 | 336 |
| R | 38 | 337 |
| Q | 39 | 338 |
| A | 40 | 339 |
| P | 41 | 340 |
| G | 42 | 341 |
| Q | 43 | 342 |
| G | 44 | 343 |
| L | 45 | 344 |
| D | 46 | 345 |
| W | 47 | 346 |
| M | 48 | 347 |
| G | 49 | 348 |
| G | 50 | 349 |
| I | 51 | 350 |
| S | 52 | 351 |
| P | 52A | 352 |
| I | 53 | 353 |
| F | 54 | 354 |
| G | 55 | 355 |
| S | 56 | 356 |
| T | 57 | 357 |
| A | 58 | 358 |
| Y | 59 | 359 |
| A | 60 | 360 |
| Q | 61 | 361 |
| K | 62 | 362 |
| F | 63 | 363 |
| Q | 64 | 364 |
| G | 65 | 365 |
| R | 66 | 366 |
| V | 67 | 367 |
| T | 68 | 368 |
| I | 69 | 369 |
| S | 70 | 370 |
| A | 71 | 371 |
| D | 72 | 372 |
| I | 73 | 373 |
| F | 74 | 374 |

| | | |
|---|---|---|
| N | 34 | 58 |
| W | 35 | 59 |
| Y | 36 | 60 |
| Q | 37 | 61 |
| Q | 38 | 62 |
| F | 39 | 63 |
| P | 40 | 64 |
| G | 41 | 65 |
| T | 42 | 66 |
| A | 43 | 67 |
| P | 44 | 68 |
| K | 45 | 69 |
| L | 46 | 70 |
| L | 47 | 71 |
| I | 48 | 72 |
| Y | 49 | 73 |
| S | 50 | 74 |
| N | 51 | 75 |
| D | 52 | 76 |
| Q | 53 | 77 |
| R | 54 | 78 |
| P | 55 | 79 |
| S | 56 | 80 |
| V | 57 | 81 |
| V | 58 | 82 |
| P | 59 | 83 |
| D | 60 | 84 |
| R | 61 | 85 |
| F | 62 | 86 |
| S | 63 | 87 |
| G | 64 | 88 |
| S | 65 | 89 |
| K | 66 | 90 |
| S | 67 | 91 |
| G | 68 | 92 |
| T | 69 | 93 |
| S | 70 | 94 |
| A | 71 | 95 |
| S | 72 | 96 |
| L | 73 | 97 |

| | | | | | |
|---|---|---|---|---|---|
| S | 75 | 375 | A | 74 | 98 |
| N | 76 | 376 | I | 75 | 99 |
| T | 77 | 377 | S | 76 | 100 |
| A | 78 | 378 | G | 77 | 101 |
| Y | 79 | 379 | L | 78 | 102 |
| M | 80 | 380 | Q | 79 | 103 |
| E | 81 | 381 | S | 80 | 104 |
| L | 82 | 382 | E | 81 | 105 |
| N | 82A | 383 | D | 82 | 106 |
| S | 82B | 384 | E | 83 | 107 |
| L | 82C | 385 | A | 84 | 108 |
| T | 83 | 386 | E | 85 | 109 |
| S | 84 | 387 | Y | 86 | 110 |
| E | 85 | 388 | Y | 87 | 111 |
| D | 86 | 389 | C | 88 | 112 |
| T | 87 | 390 | A | 89 | 113 |
| A | 88 | 391 | A | 90 | 114 |
| V | 89 | 392 | W | 91 | 115 |
| Y | 90 | 393 | D | 92 | 116 |
| F | 91 | 394 | D | 93 | 117 |
| C | 92 | 395 | S | 94 | 118 |
| A | 93 | 396 | L | 95 | 119 |
| R | 94 | 397 | K | 95A | 120 |
| H | 95 | 398 | G | 95B | 121 |
| G | 96 | 399 | A | 96 | 122 |
| N | 97 | 400 | V | 97 | 123 |
| Y | 98 | 401 | F | 98 | 124 |
| Y | 99 | 402 | G | 99 | 125 |
| Y | 100 | 403 | G | 100 | 126 |
| Y | 100A | 404 | G | 101 | 127 |
| S | 100B | 405 | T | 102 | 128 |
| G | 100C | 406 | Q | 103 | 129 |
| M | 100D | 407 | L | 104 | 130 |
| D | 101 | 408 | T | 105 | 131 |
| V | 102 | 409 | V | 106 | 132 |
| W | 103 | 410 | L | 107 | 133 |
| G | 104 | 411 | G | 108 | 134 |
| Q | 105 | 412 | Q | 109 | 135 |
| G | 106 | 413 | P | 110 | 136 |
| T | 107 | 414 | K | 111 | 137 |

| | | | | | |
|---|---|---|---|---|---|
| T | 108 | 415 | A | 112 | 138 |
| V | 109 | 416 | A | 113 | 139 |
| T | 110 | 417 | P | 114 | 140 |
| V | 111 | 418 | S | 115 | 141 |
| S | 112 | 419 | V | 116 | 142 |
| S | 113 | 420 | T | 117 | 143 |
| A | 114 | 421 | L | 118 | 144 |
| S | 115 | 422 | F | 119 | 145 |
| T | 116 | 423 | P | 120 | 146 |
| K | 117 | 424 | P | 121 | 147 |
| G | 118 | 425 | S | 122 | 148 |
| P | 119 | 426 | S | 123 | 149 |
| S | 120 | 427 | E | 124 | 150 |
| V | 121 | 428 | E | 125 | 151 |
| F | 122 | 429 | L | 126 | 152 |
| P | 123 | 430 | Q | 127 | 153 |
| L | 124 | 431 | A | 128 | 154 |
| A | 125 | 432 | N | 129 | 155 |
| P | 126 | 433 | K | 130 | 156 |
| S | 127 | 434 | A | 131 | 157 |
| S | 128 | 435 | T | 132 | 158 |
| K | 129 | 436 | L | 133 | 159 |
| S | 130 | 437 | V | 134 | 160 |
| T | 131 | 438 | C | 135 | 161 |
| S | 132 | 439 | L | 136 | 162 |
| G | 133 | 440 | I | 137 | 163 |
| G | 134 | 441 | S | 138 | 164 |
| T | 135 | 442 | D | 139 | 165 |
| A | 136 | 443 | F | 140 | 166 |
| A | 137 | 444 | Y | 141 | 167 |
| L | 138 | 445 | P | 142 | 168 |
| G | 139 | 446 | G | 143 | 169 |
| C | 140 | 447 | A | 144 | 170 |
| L | 141 | 448 | V | 145 | 171 |
| V | 142 | 449 | T | 146 | 172 |
| K | 143 | 450 | V | 147 | 173 |
| D | 144 | 451 | A | 148 | 174 |
| Y | 145 | 452 | W | 149 | 175 |
| F | 146 | 453 | K | 150 | 176 |
| P | 147 | 454 | A | 151 | 177 |

| | | | | | |
|---|---|---|---|---|---|
| E | 148 | 455 | D | 152 | 178 |
| P | 149 | 456 | S | 153 | 179 |
| V | 150 | 457 | S | 154 | 180 |
| T | 151 | 458 | P | 155 | 181 |
| V | 152 | 459 | V | 156 | 182 |
| S | 153 | 460 | K | 157 | 183 |
| W | 154 | 461 | A | 158 | 184 |
| N | 155 | 462 | G | 159 | 185 |
| S | 156 | 463 | V | 160 | 186 |
| G | 157 | 464 | E | 161 | 187 |
| A | 158 | 465 | T | 162 | 188 |
| L | 159 | 466 | T | 163 | 189 |
| T | 160 | 467 | T | 164 | 190 |
| S | 161 | 468 | P | 165 | 191 |
| G | 162 | 469 | S | 166 | 192 |
| V | 163 | 470 | K | 167 | 193 |
| H | 164 | 471 | Q | 168 | 194 |
| T | 165 | 472 | S | 169 | 195 |
| F | 166 | 473 | N | 170 | 196 |
| P | 167 | 474 | N | 171 | 197 |
| A | 168 | 475 | K | 172 | 198 |
| V | 169 | 476 | Y | 173 | 199 |
| L | 170 | 477 | A | 174 | 200 |
| Q | 171 | 478 | A | 175 | 201 |
| S | 172 | 479 | S | 176 | 202 |
| S | 173 | 480 | S | 177 | 203 |
| G | 174 | 481 | Y | 178 | 204 |
| L | 175 | 482 | L | 179 | 205 |
| Y | 176 | 483 | S | 180 | 206 |
| S | 177 | 484 | L | 181 | 207 |
| L | 178 | 485 | T | 182 | 208 |
| S | 179 | 486 | P | 183 | 209 |
| S | 180 | 487 | E | 184 | 210 |
| V | 181 | 488 | Q | 185 | 211 |
| V | 182 | 489 | W | 186 | 212 |
| T | 183 | 490 | K | 187 | 213 |
| V | 184 | 491 | S | 188 | 214 |
| P | 185 | 492 | H | 189 | 215 |
| S | 186 | 493 | R | 190 | 216 |
| S | 187 | 494 | S | 191 | 217 |

| | | |
|---|---|---|
| S | 188 | 495 |
| L | 189 | 496 |
| G | 190 | 497 |
| T | 191 | 498 |
| Q | 192 | 499 |
| T | 193 | 500 |
| Y | 194 | 501 |
| I | 195 | 502 |
| C | 196 | 503 |
| N | 197 | 504 |
| V | 198 | 505 |
| N | 199 | 506 |
| H | 200 | 507 |
| K | 201 | 508 |
| P | 202 | 509 |
| S | 203 | 510 |
| N | 204 | 511 |
| T | 205 | 512 |
| K | 206 | 513 |
| V | 207 | 514 |
| D | 208 | 515 |
| K | 209 | 516 |
| R | 210 | 517 |
| V | 211 | 518 |

| | | |
|---|---|---|
| Y | 192 | 218 |
| S | 193 | 219 |
| C | 194 | 220 |
| Q | 195 | 221 |
| V | 196 | 222 |
| T | 197 | 223 |
| H | 198 | 224 |
| E | 199 | 225 |
| G | 200 | 226 |
| S | 201 | 227 |
| T | 202 | 228 |
| V | 203 | 229 |
| E | 204 | 230 |
| K | 205 | 231 |
| T | 206 | 232 |
| V | 207 | 233 |
| A | 208 | 234 |
| P | 209 | 235 |
| T | 210 | 236 |
| E | 211 | 237 |
| C | 212 | 238 |
| S | 213 | 239 |
